# Adolescent stress impairs postpartum social behavior via anterior insula-prelimbic pathway

**DOI:** 10.1101/2023.01.03.522598

**Authors:** Kyohei Kin, Jose Francis-Oliveira, Shin-ichi Kano, Minae Niwa

## Abstract

Adolescent stress can be a risk factor for abnormal social behavior in the postpartum period, which critically affects the safety of mothers and children. Nonetheless, the underlying mechanisms remain unclear. Using a newly established mouse model with optogenetics and *in vivo* calcium imaging, we found that adolescent psychosocial stress, combined with pregnancy and delivery, caused hypofunction of the glutamatergic pathway from the anterior insula to prelimbic cortex (AI-PrL pathway), which altered PrL neuronal activity, and in turn led to abnormal social behavior. Specifically, the AI-PrL pathway played a crucial role during recognizing the novelty of other mice by modulating “stable neurons” in PrL, which were constantly activated or inhibited by novel mice. We also observed that glucocorticoid receptor signaling in the AI-PrL pathway had a causal role in stress-induced postpartum changes. Our findings provide novel and functional insights into a cortico-cortical pathway underlying adolescent stress-induced postpartum social behavioral deficits.

---

Pregnancy and childbirth are life events that bring significant physiological and psychological changes to the mother in any mammals ^1^. Human studies indicate that exposure to various types of stress in earlier life stages can cause postpartum behavioral changes ^2–4^. It is suggested that even mild early life stress exposure can cause behavioral changes when combined with pregnancy and delivery. This idea may even explain the prevalence of perinatal complications, such as postpartum mental disorders ^5^. However, ambiguity in the definition of early life stress has prevented us from having a detailed understanding of the relationship between early life stress and postpartum behavioral changes. In most human studies, early life stress includes stress exposure from the postnatal to late adolescent periods. Considering the heterogeneity of development during these periods ^6^, stress-induced changes would also likely be heterogeneous. Thus, focusing on the effects of stress exposure during a specific life stage may bring us further insights into the underlying mechanisms.

Postpartum social behavior is important for the mother’s health, as well as for the development of her children ^5, 7–10^. Unfortunately, postpartum social behavior is sensitive to stress that can occur at different points during the mother’s lifespan. Stress during adolescence, a critical time when hormone levels change dramatically and neural pathways are fine-tuned to facilitate the transition from childhood to adulthood ^6, 11, 12^, can be associated with changes in later postpartum behavior, including social behavior ^2–4, 12–16^. Nonetheless, the neural circuit mechanisms by which adolescent stress leads to changes in postpartum social behavior are unclear. Focusing on the adolescent phase in rodents may shed light on this unaddressed question. However, appropriate animal models have not been well established until recently. We have recently found that mice exposed to social isolation in late adolescence (SILA), which alone causes no endocrine or behavioral changes, show long-lasting behavioral changes only when accompanied by pregnancy and delivery ^17^. These behavioral changes are observed one week postpartum , not immediately after delivery, and last for at least three weeks postpartum ^17^. Further studies using this new mouse model will greatly advance our understanding of how SILA affects postpartum behaviors. In the present study, using this new mouse model, we have focused on the prelimbic cortex (PrL), which is a hub brain region that plays a crucial role in social behavior and regulates stress responses ^18–21^, and examined a PrL-related neural mechanism by which adolescent psychosocial stress (i.e., social isolation) affects postpartum social behavior.

## Results

### Postpartum AI-PrL pathway impaired by SILA

We previously found that stressed dams (mice exposed to SILA and then gave birth to pups) showed reduced preference for social novelty in the social interaction test (SIT), consisting of a sociability trial (S-trial) and social novelty trial (SN-trial), which were designed to progressively increase the complexity of the social context within a single day ^22–24^, at one-week postpartum relative to unstressed dams (healthy controls and group-housed mice that gave birth to pups) ^17^. In the present study, we examined the specific characteristics of social behavioral changes in stressed dams in more detail with a modified SIT, in which S- and SN-trials were performed on two consecutive days (Fig. 1A). SILA consisted of no interaction with other mice and confinement in opaque, wire-topped polypropylene cages from five to eight weeks of age.

**Fig. 1.**
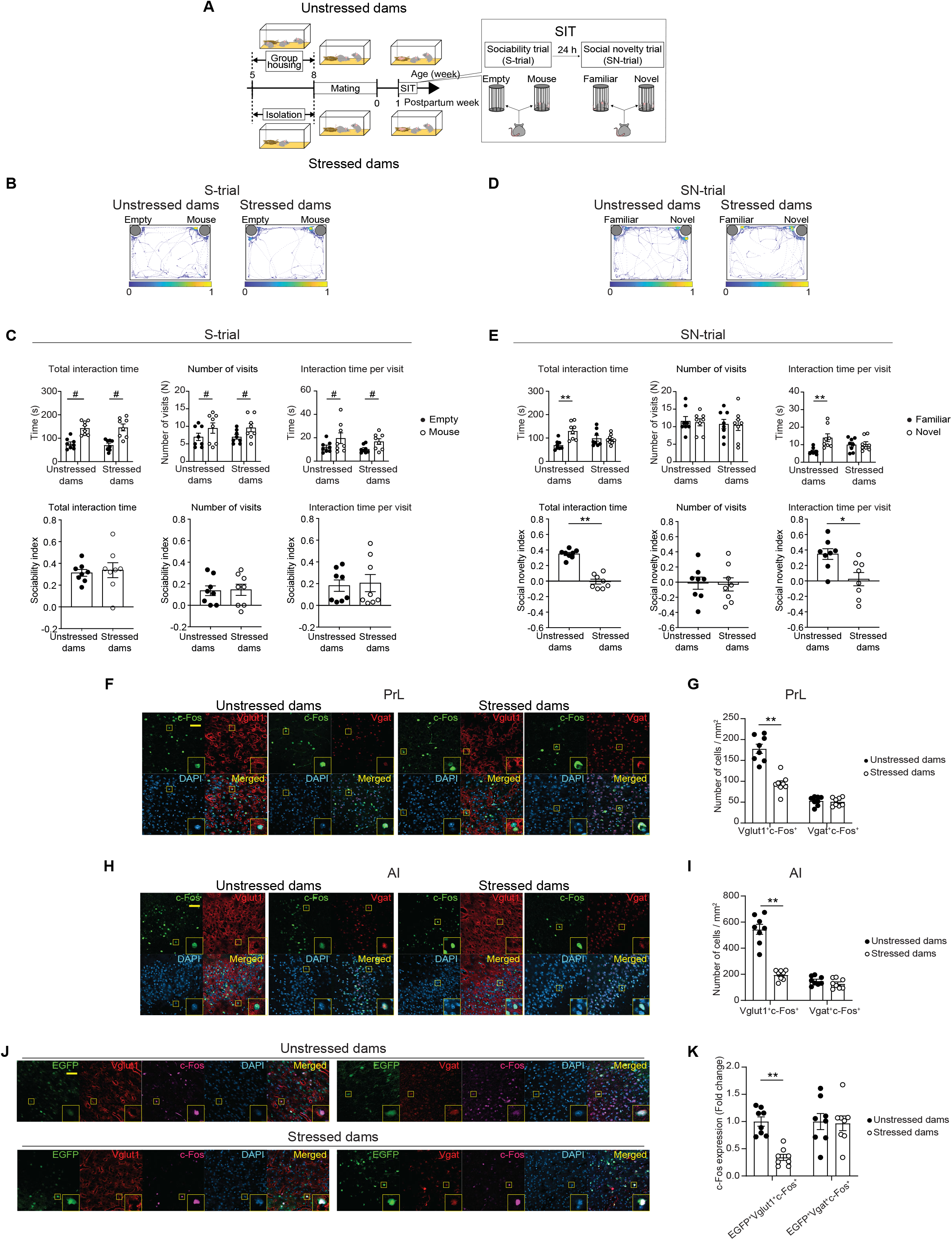
SILA induced postpartum behavioral changes in SN-trial and reduced neural activity in AI and PrL. (**A**) Experimental design for SIT experiments. (**B**) Representative heatmaps of the mouse track during S-trials. (**C**) Both unstressed and stressed dams showed sociability toward a mouse in the S-trials (two-way mixed ANOVA). (**D**) Representative heatmaps of the mouse track during the SN-trials. (**E**) Stressed dams did not show a novel mouse preference in the SN-trials, compared to unstressed dams (two-way mixed ANOVA). Social preference indexes for S- and SN-trials were also shown in **Figs. 1C** and **1E** (Student’s *t* test, Cohen’s D = 4.766 for total interaction time in the SN-trial, and Cohen’s D = 1.478 for interaction time per visit in the SN-trial). (**F**) Representative images of c-Fos^+^, Vglut1^+^ or Vgat^+^, DAPI, and colocalized cells in PrL of unstressed or stressed dams. Scale bar, 50 µm. (**G**) Stressed dams showed decreased number of Vglut1^+^c-Fos^+^ cells, but not Vgat^+^c-Fos^+^ cells, in PrL after the SN-trial (two-way mixed ANOVA). (**H**) Representative images of c-Fos^+^, Vglut1^+^ or Vgat^+^, DAPI, and colocalized cells in AI of unstressed or stressed dams. Scale bar, 50 µm. (**I**) Stressed dams showed decreased number of Vglut1^+^c-Fos^+^ cells, but not Vgat^+^c-Fos^+^ cells, in AI after the SN-trials (two-way mixed ANOVA). (**J**) Representative images of EGFP^+^, c-Fos^+^, Vglut1^+^ or Vgat^+^, DAPI, and colocalized cells in AI of unstressed or stressed dams injected with AAVretro-hSyn-EGFP into PrL. Scale bar, 50 µm. (**K**) Stressed dams showed decreased number of EGFP^+^Vglut1^+^c-Fos^+^ cells, but not EGFP^+^Vgat^+^c-Fos^+^ cells, in AI after the SN-trial. Scale bar, 50 µm. (two-way mixed ANOVA). All data are represented as mean ± SEM. * = *post hoc* Bonferroni, *p* < 0.05. ** = *post hoc* Bonferroni, *p* < 0.01. ^#^ = ANOVA main effect for the brain region, *p* < 0.05.

Immediately after SILA, no behavioral changes were observed in SIT (Supplementary Fig. 1A-C). Each female was then mated with a healthy B6J male mouse at eight weeks of age, and gave birth to pups. Then, SIT was conducted one week after delivery (Fig. 1A). In the S-trials, we observed social preference with no behavioral changes between stressed and unstressed dams in the total interaction time, number of visits, and interaction time per visit with a mouse relative to an empty enclosure (Fig. 1, B and C). In the SN-trials, unstressed dams spent significantly longer time interacting with a novel mouse than a familiar mouse, indicating a novel mouse preference. This result was due to longer interaction times per visit, and not due to any significant differences in the frequency of visits to the familiar and novel mice. In contrast, stressed dams showed no difference in total interaction time between the novel and familiar mice. No differences were also found in the frequency of visits and interaction time per visit (Fig. 1, D and E). The preference indexes for SIT were calculated and indicated a robust behavioral change in the SN-trials in stressed dams compared to unstressed dams (effect size > 1) (Fig. 1E, bottom panels). Thus, we analyzed our subsequent SIT data using the preference indexes for better clarity. These results indicated that SILA, in conjunction with pregnancy and delivery, impaired behavior in social novelty preference during the postpartum period.

As a first step for assessing the underlying neural mechanisms, we investigated whether PrL neuronal activity was altered during the social behavioral changes observed in stressed dams. A decrease in c-Fos immunoreactivity, an indicator of neuronal activity, was observed in PrL after the SN-trials in stressed dams compared to unstressed dams (Supplementary Fig. 1D), and this change was observed primarily in glutamatergic neurons, but not GABAergic neurons (Fig. 1F and G). We next examined neuronal connections in direct upstream regions of PrL, using both B6J male and female mice in normal group-housed conditions. Retrograde tracing was performed by injecting a retrograde adeno-associated virus (AAV) expressing EGFP (AAV-retro-hSyn-EGFP) into PrL (Supplementary Fig. 1E) ^25^. We observed EGFP labeling in many PrL inputs, which were mainly ipsilateral, with no sex-dependent differences (Supplementary Fig. 1, F and G). Systematic examination of c-Fos immunoreactivity in these upstream regions after the SN-trials revealed that the number of c-Fos^+^ neurons was significantly reduced in stressed dams compared to unstressed dams in only two brain regions: mostly in the anterior insula (AI, effect size = 3.6), and to a lesser extent in the basolateral amygdala (BLA, effect size = 1.4) (Supplementary Fig. 1H). Similar to the findings in PrL, decreased activity in AI was observed in stressed dams compared to unstressed dams, mainly in glutamatergic neurons, but not in GABAergic neurons (Fig. 1, H and I). Of note, the pathway from AI to PrL mainly consisted of glutamatergic neurons, in both male and female mice (Supplementary Fig. 1, I and J). Further, c-Fos immunoreactivity was reduced in glutamatergic neurons, but not in GABAergic neurons, in the AI-PrL pathway in stressed dams compared to unstressed dams (Fig. 1, J and K). These results suggested that the activity in glutamatergic neurons of PrL and the AI-PrL pathway was reduced when SILA was accompanied by pregnancy and delivery.

### Anatomical features of the AI-PrL pathway

To characterize the anatomical features of the AI-PrL pathway, anterograde- or dual-viral tracing strategies were employed ^26^. Since our data showed that the AI-PrL pathway was predominantly glutamatergic, we used anterograde tracing via injection of AAV-Flex-tdTomato into AI of vesicular glutamate transporter 1 (*Slc17a7*, Vglut1)-Cre mice. Glutamatergic pathways from AI projected to all PrL layers, but mainly to deep layers V and VI (Supplementary Fig. 2, A and B). Conversely, anterograde tracing via injection of AAV-Flex-tdTomato into PrL of Vglut1-Cre mice did not show any pathways from PrL to AI (Supplementary Fig. 2, C and D). To identify potential collateral projections of the AI-PrL pathway, dual viral tracing was performed by injecting AAV-retro-hSyn-Cre and AAV-flex-tdTomato into PrL and AI of B6J mice, respectively. We only observed tdTomato^+^ cells in AI and fibers in PrL (Supplementary Fig. 2, E and F). Thus, these virus-tracing studies indicated that the AI-PrL pathway was a unidirectional pathway from AI to PrL with no collateral projections.

### Role of the AI-PrL pathway in postpartum social novelty behavior

Based on the findings that stressed dams showed decreased activity in the AI-PrL pathway and reduced preference for social novelty, and reports that AI was responsive to aversive stimuli and a core region of cognitive and motivational processes ^27, 28^, we hypothesized that SILA, in conjunction with pregnancy and delivery, would lead to decreased activity in PrL glutamatergic neurons (hereafter, referred to as PrL neurons) and subsequent social novelty recognition in the postpartum period via hypofunction of the AI-PrL pathway. To test this hypothesis directly, we investigated how modulation of the AI-PrL pathway altered PrL neuronal activity and social behavior using *in vivo* microendoscopic calcium imaging in conjunction with optogenetics (Fig. 2, A and B). We used eNPHR3.0 for optogenetic inhibition of the AI-PrL pathway in unstressed dams (Unstressed dams / eNPHR) and ChrimsonR for optogenetic activation of the AI-PrL pathway in stressed dams (Stressed dams / Chrimson).

**Fig. 2.**
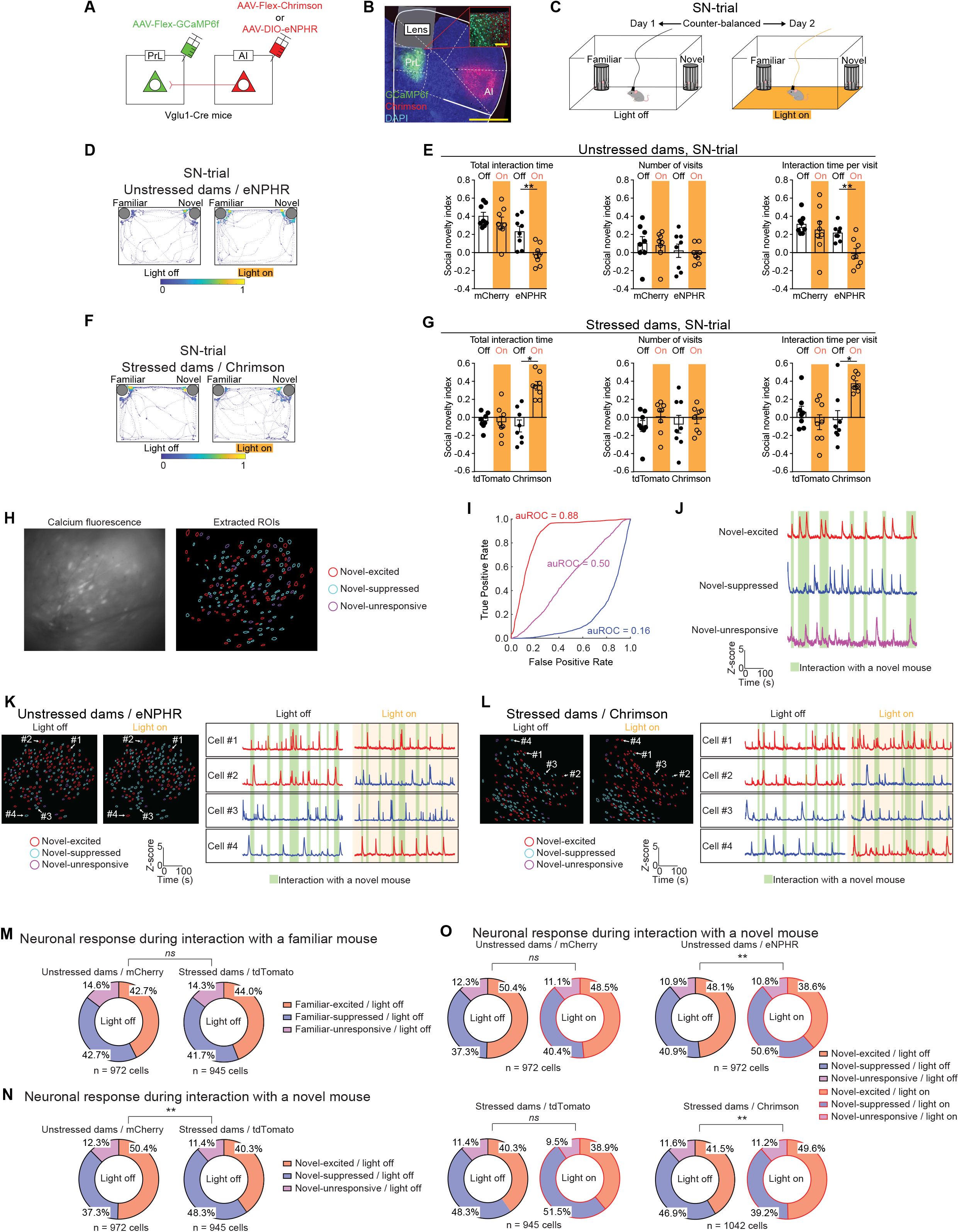
Hypofunction of the AI-PrL pathway underlay SILA-induced reduction of PrL neuronal activity and subsequent behavioral changes in social novelty recognition in the postpartum period. **(A)** Strategy of *in vivo* microendoscopic calcium recording of PrL through optogenetic manipulation of the AI-PrL pathway. **(B)** Representative image showing expression of Chrimson in the AI-PrL pathway and GCaMP6f underneath a GRIN lens. Scale bars, 1 mm and 100 µm. **(C)** Experimental timeline of SIT without and with optogenetic manipulation. **(D)** Representative heatmaps of the mouse track during the SN-trials in unstressed dams expressing eNPHR. **(E)** Optogenetic inhibition of the AI-PrL pathway decreased social novelty preference in total interaction time and interaction time per visit (two-way mixed ANOVA, Wilcoxon signed-rank test). **(F)** Representative heatmaps of the mouse track during the SN-trials in stressed dams expressing Chrimson. **(G)** Optogenetic activation of the AI-PrL pathway increased social novelty preference in total interaction time and interaction time per visit (two-way mixed ANOVA, Wilcoxon signed-rank test). **(H)** Imaging field of view showing raw calcium fluorescence and regions of interest (ROIs) corresponding to single neurons. **(I)** Receiver operating characteristic (ROC) curves computed from three example neurons that were categorized as novel-excited [area under ROC curve (auROC) = 0.88], novel-unresponsive (auROC = 0.50), or novel-suppressed (auROC = 0.16). **(J)** Representative calcium traces from novel-excited (top), novel-suppressed (middle), and novel-unresponsive (bottom) neurons. **(K)** Representative extracted ROIs and calcium traces from PrL in an unstressed dam with optogenetic inhibition of the AI-PrL pathway. **(L)** Representative extracted ROIs and calcium traces from PrL in a stressed dam with optogenetic activation of the AI-PrL pathway. **(M)** PrL activity during interaction with a familiar mouse did not show any differences between stressed and unstressed dams expressing control viruses (Chi-squared test). **(N)** Fractions of novel-excited and novel-suppressed cells in stressed dams expressing control viruses were significantly decreased and increased, respectively, in comparison to unstressed dams expressing control viruses (Chi-squared test). (O) Optogenetic inhibition in unstressed dams decreased and increased the fractions of novel-excited and novel-suppressed neurons in PrL, respectively (Chi-squared test). Optogenetic activation of the AI-PrL pathway in stressed dams increased and decreased the fractions of novel-excited and novel-suppressed neurons in PrL, respectively (Chi-squared test). All data are represented as mean ± SEM. For ANOVAs, * indicates statistical significance for *post hoc* Bonferroni comparisons. * = *post hoc* Bonferroni, *p* < 0.05. ** = *post hoc* Bonferroni, *p* < 0.01. *ns* = non-significant.

First, the functional relevance of the AI-PrL pathway on PrL neuronal activity was confirmed by a combination of optogenetics and *in vivo* microendoscopic calcium imaging in the home cage, according to previously described methods ^29^. Simultaneous monitoring of calcium dynamics in PrL neurons showed that optogenetic inhibition or activation of the AI-PrL pathway decreased or increased the calcium transient rate and its average amplitude in PrL neurons, respectively (Supplementary Fig. 3, A and B). Next, to examine whether hypofunction of the AI-PrL pathway underlay the behavioral changes observed in stressed dams, we used optogenetic manipulation of the AI-PrL pathway during SIT (Fig. 2C and Supplementary Fig. 3C). Optogenetic manipulation of the AI-PrL pathway in unstressed and stressed dams did not affect behavior during the S-trials (Supplementary Fig. 3, D and E). Notably, optogenetic inhibition of the AI-PrL pathway reduced social interaction with novel mice in the SN-trials in unstressed dams (Fig. 2, D and E). In contrast, optogenetic activation of the AI-PrL pathway ameliorated behavioral changes in the SN-trials in stressed dams (Fig. 2, F and G). These results indicated that hypofunction of the AI-PrL pathway was responsible for SILA-induced changes in social novelty recognition that occurred in the postpartum period. Consistent with these behavioral changes induced by optogenetic manipulation, c-Fos immunoreactivity in PrL after SN-trials was decreased by optogenetic inhibition in unstressed dams, and increased by optogenetic activation in stressed dams (Supplementary Fig. 3, F and G).

We then used *in vivo* microendoscopic calcium imaging to visualize the changes in PrL neuronal activity induced by manipulation of the AI-PrL pathway during SIT. Recorded calcium images were analyzed with a receiver operating characteristic (ROC) analysis (Methods and Fig. 2, H to J) ^30–32^. This approach enabled us to identify subsets of PrL neurons that were excited or suppressed during behavioral testing, as well as changes in PrL activity induced by optogenetic manipulation of the AI-PrL pathway (see Supplementary Table 2 for the definitions of subsets, Fig. 2, K and L). PrL activity patterns during the S-trials did not show any differences between unstressed and stressed dams expressing control viruses (Supplementary Fig. 4, A and B).

Optogenetic manipulation of the AI-PrL pathway also did not affect PrL activity patterns during the S-trials (Supplementary Fig. 4, C and D). These findings were consistent with the lack of behavioral differences we observed during the S-trials (Supplementary Fig. 3, D and E). Notably, PrL activity patterns during interactions with a novel mouse, but not a familiar mouse, during the SN-trials were significantly different in stressed dams expressing control viruses compared to unstressed dams expressing control viruses. Specifically, the fractions of novel-excited / light off and novel-suppressed / light off neurons were significantly decreased and increased, respectively, in stressed dams compared to unstressed dams (Fig. 2, M and N). Such changes were not observed following SILA alone, consistent with the finding that no behavioral changes were observed during SIT (Supplementary Fig. 1, B and C, Supplementary Fig. 5). Furthermore, optogenetic manipulation of the AI-PrL pathway during the SN-trials changed PrL activity patterns. We observed a significant decrease and increase in the fractions of novel-excited and novel-suppressed neurons, respectively, by optogenetic inhibition in unstressed dams. In stressed dams, the fractions of novel-excited and novel-suppressed neurons were increased and decreased, respectively, by optogenetic activation (Fig. 2O and Supplementary Fig. 4E). These results suggested that hypofunction of the AI-PrL pathway was causally linked to SILA-induced changes in social novelty recognition, but not sociability, during the postpartum period.

### Specificity of the AI-PrL pathway in regulating postpartum social novelty behavior

As mentioned above, another significantly altered region in stressed dams after SN-trials was BLA (Supplementary Fig. 1H). To evaluate if the AI-PrL pathway specifically led to changes in social novelty recognition during the postpartum period, we also investigated the involvement of the BLA-PrL pathway in such behavior. As in PrL and AI, c-Fos expression was reduced in stressed dams compared to unstressed dams, mainly in glutamatergic neurons, but not in GABAergic neurons (Supplementary Fig. 6, A and B). Since the pathway from BLA to PrL mainly consisted of glutamatergic neurons (Supplementary Fig. 6, C and D), we also conducted optogenetic manipulation of the glutamatergic pathway from BLA to PrL (BLA-PrL pathway) during SIT (Supplementary Fig. 6, E and F). Unlike manipulation of the AI-PrL pathway, optogenetic manipulation of the BLA-PrL pathway did not affect the behavioral outcomes of both S- and SN-trials in SIT when comparing unstressed and stressed dams (Supplementary Fig. 6, G and H). PrL activity patterns were also not affected by optogenetic manipulation of the BLA-PrL pathway (Supplementary Fig. 6, I and J). These data suggested that the BLA-PrL pathway might not modulate SILA-induced changes in PrL neuronal activity and social behavior during the postpartum period. Therefore, we focused our efforts on further examination of the role of the AI-PrL pathway in postpartum social behavior.

### Role of the AI-PrL pathway in PrL stable neurons during social novelty behavior

Next, we examined which neuronal populations in PrL were modulated by optogenetic manipulation of the AI-PrL pathway during the SN-trials. We used longitudinal cell registration of PrL neuronal activity in animals expressing control viruses during the SN-trials on two consecutive days (SN-trials with and without light stimulation) to classify individual neurons into nine categories, based on their activity patterns as defined by ROC analysis on each day (Fig. 3A). In unstressed dams expressing control viruses, about half of the recorded neurons showed the same activity patterns during interaction with a novel mouse on two consecutive days (“stable neurons”), while the other half showed different responses to a novel mouse on two consecutive days (dynamic neurons) (Fig. 3, A and B). The fractions of these categories of neurons were significantly different in stressed dams compared to unstressed dams, both expressing control viruses. The value of the adjusted residual (AR) indicated that these differences were mainly due to changes in the fraction of stable neurons. In stressed dams expressing control viruses, the fraction of neurons excited during interaction with a novel mouse on two consecutive days was decreased, and the fraction of neurons suppressed during interaction with a novel mouse on two consecutive days was increased, compared to unstressed dams expressing control viruses (Fig. 3B). These data suggested that, despite the presence of neurons that could alter their activity patterns in response to novel social stimuli on two consecutive days, SILA induced a significant change in the response types of stable neurons in PrL.

**Fig. 3.**
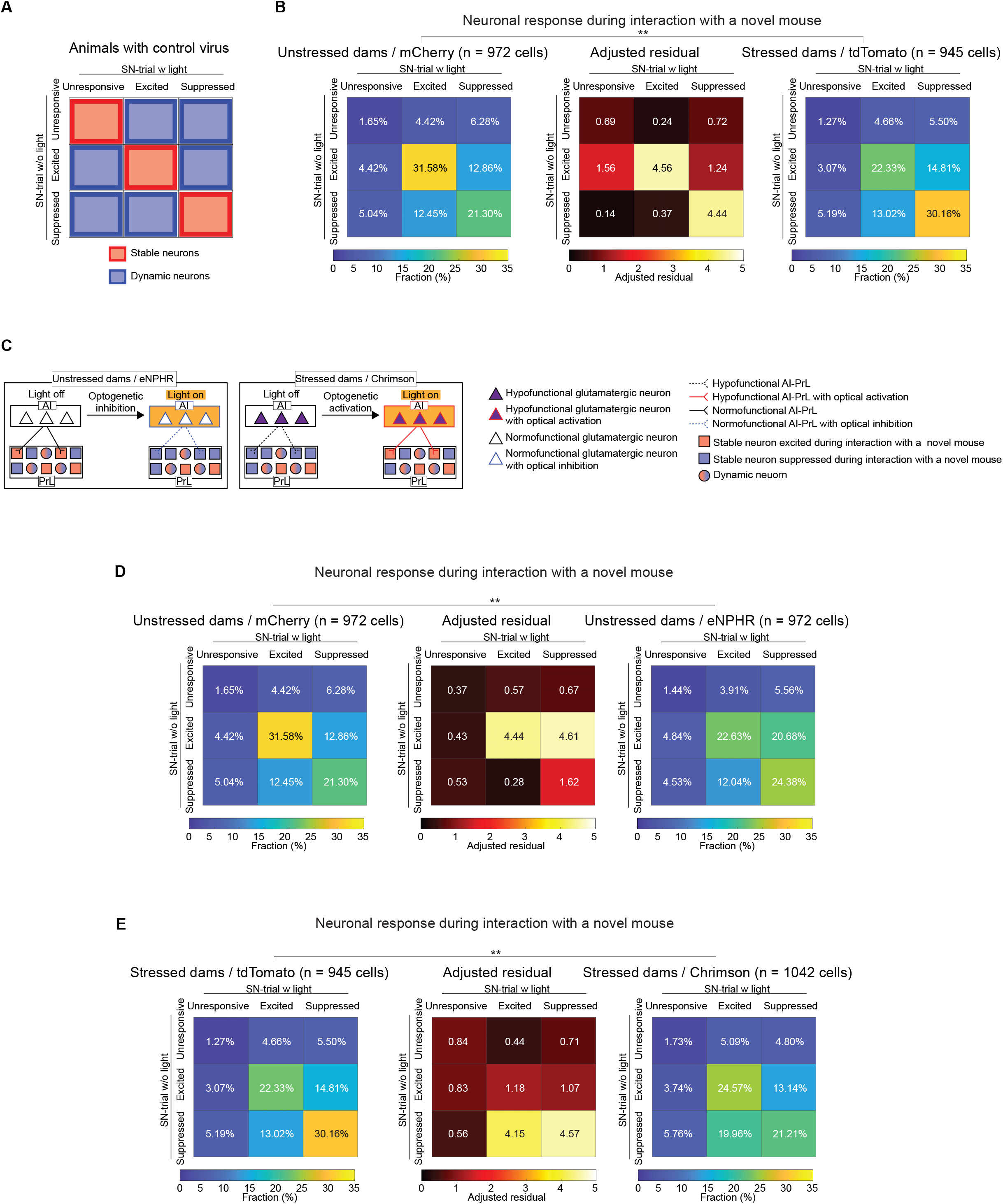
AI-PrL pathway affected the activity of stable neurons in PrL during the SN-trials. (**A**) Stable and dynamic neurons defined by longitudinal registration of calcium imaging over two consecutive days (SN-trials with and without light stimulation) from animals expressing only control viruses, not opsin. Even without optogenetic manipulation, several PrL neurons showed the same neuronal activity patterns during SN-trials on two consecutive days (defined as “stable neurons”), while others showed different activity patterns (defined as “dynamic neurons”). (**B**) Significant differences in the patterns of PrL activity changes between SN-trials on two consecutive days were observed between stressed and unstressed dams expressing control viruses (Chi-squared test). (**C**) Scheme of our hypothesis regarding the effect of the AI-PrL pathway on PrL activity during SN-trials. (**D**) Significant differences in the patterns of PrL activity changes from SN-trials without light stimulation to SN-trials with light stimulation were observed between unstressed dams expressing control viruses and eNPHR (Chi-squared test). (**E**) Significant differences in the patterns of PrL activity changes from SN-trials without light stimulation to SN-trials with light stimulation were observed between stressed dams expressing control viruses and Chrimson (Chi-squared test). ** = *p* < 0.01.

Based on these findings, we hypothesized that the AI-PrL pathway modulated the activity of stable, but not dynamic, neurons in PrL, and was essential for keeping the excitation state of stable neurons during interaction with a novel mouse (Fig. 3C). To address our hypothesis, we analyzed the data from calcium imaging of PrL neurons with optogenetic manipulation of the AI-PrL pathway during the SN-trials in unstressed and stressed dams. Optogenetic manipulation of the AI-PrL pathway resulted in significant differences in the patterns of PrL activity changes between SN-trials without and with light stimulation (Fig. 3, D and E). As light stimulation in animals with opsins would affect the neuronal activity in PrL, stable and dynamic neurons could not be defined directly with data from these animals. Therefore, we evaluated the differences between animals with control viruses and opsins, and speculated on the changes in stable and dynamic neurons. The AR values indicated that optogenetic inhibition in unstressed dams decreased the fraction of neurons that were excited regardless of light stimulation, and increased the fraction of neurons that were excited during light off and suppressed during light on. These data suggested that optogenetic inhibition of the AI-PrL pathway in unstressed dams modulated stable neurons, which were excited during interactions with a novel mouse under a normofunctional AI-PrL pathway [categorized as excited regardless of light stimulation in unstressed dams / mCherry (31.58%)], into suppressed neurons during interactions with a novel mouse [categorized as excited without light stimulation and suppressed with light stimulation in unstressed dams / eNPHR (20.68%)] (Fig. 3D). In contrast, optogenetic activation in stressed dams increased the fraction of neurons that were suppressed during light off and excited during light on, and decreased the fraction of neurons that were suppressed regardless of light stimulation. These data suggested that optogenetic activation of the AI-PrL pathway in stressed dams turned stable neurons, which were suppressed due to a hypofunctional AI-PrL pathway [categorized as suppressed regardless of light stimulation in stressed dams / tdTomato (30.16%)], into excited neurons during interactions with a novel mouse [categorized as suppressed without light stimulation and excited with light stimulation in stressed dams / Chrimson (19.96%)] (Fig. 3E). In addition, the pattern of the PrL activity change in stressed dams / Chrimson from SN-trials with light stimulation (presumably under a normofunctional AI-PrL pathway) to SN-trials without light stimulation (presumably under a hypofunctional AI-PrL pathway) was similar to that from SN-trials in unstressed dams / eNPHR without light stimulation (presumably under a normofunctional AI-PrL pathway) to SN-trials with light stimulation (presumably under a hypofunctional AI-PrL pathway) (Supplementary Fig. 7). These data suggested that optogenetic inhibition of the AI-PrL pathway in unstressed dams and optogenetic activation of the AI-PrL pathway in stressed dams showed opposite effects on the patterns of PrL activity changes between SN-trials with and without light stimulation. These results indicated that the AI-PrL pathway modulated stable neurons in PrL, and was required to excite stable neurons during interactions with a novel mouse.

### Role of the AI-PrL pathway specific to postpartum social novelty behavior

Optogenetic manipulation of the AI-PrL pathway did not affect the patterns of PrL activity changes between S-trials with and without light stimulation, and those during interactions with a familiar mouse between SN-trials with and without light stimulation (Supplementary Fig. 8). These results suggested that modulation of the AI-PrL pathway did not affect PrL neuronal activity during S-trials or interactions with a familiar mouse in SN-trials. These findings indicated that the AI-PrL pathway specifically regulated social novelty behavior, but not sociability behavior, and that activating this pathway was sufficient to ameliorate the behavioral changes observed in stressed dams. It was possible that the AI-PrL pathway regulated social novelty behavior only under pathological conditions. Therefore, we investigated whether this pathway was physiologically relevant in animals under normal conditions. We performed the same experiments as described above in virgin mice without SILA in both sexes (Supplementary Fig. 9A). We found that optogenetic inhibition of the AI-PrL pathway in male and female virgin mice led to behavioral changes in SN-trials, but not S-trials (Supplementary Fig. 9, B to E). Furthermore, calcium imaging of PrL neurons showed that the effect of optogenetic inhibition on PrL activity in virgin mice was similar to that in unstressed dams (Supplementary Fig. 10, A to D, and Supplementary Fig. 11, A and B). Thus, these results suggested that the AI-PrL pathway had a crucial role in social novelty recognition even in physiological conditions, i.e., in mice not exposed to SILA and without pregnancy and delivery.

To investigate whether the AI-PrL pathway contributed to the recognition of non-social information such as objects, we performed a novel objective recognition test (NOR) (Supplementary Fig. 12A). Optogenetic manipulation of the AI-PrL pathway had no effect on preference for a novel object or PrL activity patterns during NOR in both unstressed and stressed dams (Supplementary Fig. 12, B to D). Further, NOR did not identify any behavioral changes in stressed dams, regardless of the modification of the experimental procedure or time point (Supplementary Fig. 12, E-I). These results indicated that normofunction of the AI-PrL pathway was required specifically for social novelty behavior, under both pathological and physiological conditions, and in both sexes.

### AI-PrL pathway function during social interactions

Novel mouse preference requires appropriate recognition of other mice, and elicits positive valence to seek social contact ^33^. We thus examined whether the AI-PrL pathway influenced valence processing in general using the real-time place preference test (RTPP). Optogenetic manipulation of the AI-PrL pathway did not induce either reward or aversion in RTPP (Supplementary Fig. 12, J and K). These results suggested that the AI-PrL pathway might play an important role in encoding the recognition of other mice, but not associated valence, negative or positive.

There were no significant differences in the frequency of visits to familiar and novel mice between stressed and unstressed dams, and optogenetic manipulation of the AI-PrL pathway in unstressed and stressed dams did not change the frequency of visits to familiar or novel mice (Fig. 1E and Fig. 2, E and G). Thus, we hypothesized that the AI-PrL pathway, which was involved in stress-induced changes in social novelty recognition, functioned primarily during interactions with other mice, and not during exploration. To address this hypothesis, we employed behavioral closed-loop optogenetic manipulation in which light stimulation was applied only when the tested animals were exploring or interacting with other mice (Fig. 4A and Supplementary Fig. 13A). Optogenetic inhibition of the AI-PrL pathway in unstressed dams during interaction, but not during exploration, induced behavioral changes in social novelty recognition (Fig. 4, B and C). Optogenetic activation of the AI-PrL pathway in stressed dams during interaction, but not during exploration, ameliorated the behavioral changes during SN-trials (Fig. 4, D and E). During S-trials, behavioral closed-loop optogenetic manipulation did not affect behavior or PrL activity (Supplementary Fig. 13, B to E). Consistent with the behavioral findings, optogenetic inhibition of the AI-PrL pathway in unstressed dams during interaction, but not during exploration, led to similar changes in PrL activity to those observed in stressed dams. Optogenetic activation of the AI-PrL pathway in stressed dams during interaction, but not during exploration, normalized the changes in PrL neuronal activity (Supplementary Fig. 13F and Fig. 4F). Further, optogenetic inhibition during interaction in unstressed dams, and optogenetic activation during interaction in stressed dams, showed opposite effects on the patterns of PrL activity changes between SN-trials with and without light stimulation (Supplementary Fig. 13G). These results were similar to those from optogenetic manipulation during entire SN-trials (Supplementary Fig. 7). Together, our results indicated that the AI-PrL pathway played a crucial role in SILA-induced changes in social novelty recognition during interactions with other mice, rather than exploration, by modulating stable neurons in PrL.

**Fig. 4.**
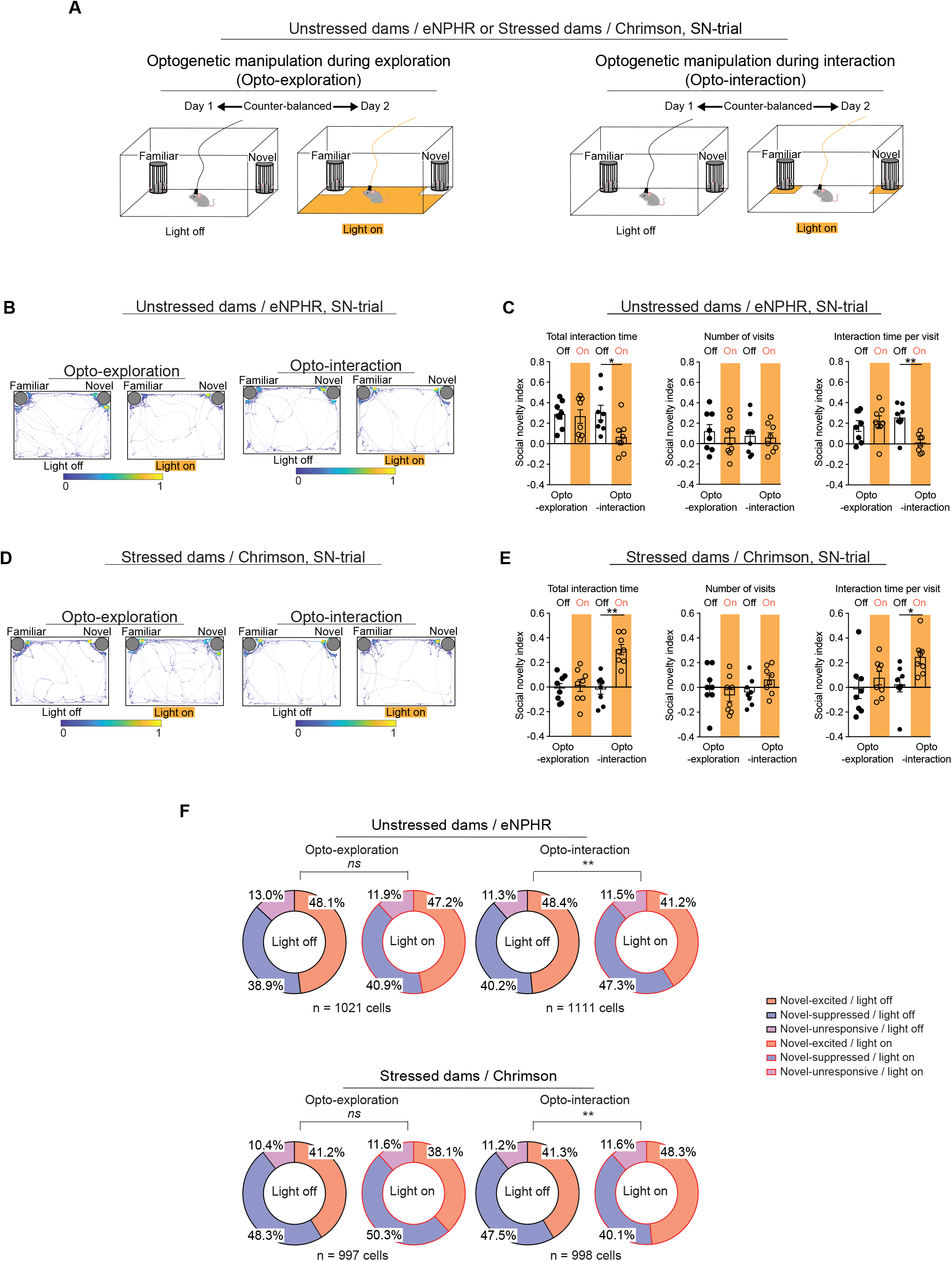
AI-PrL pathway was involved in SILA-induced changes in postpartum PrL neuronal activity and social novelty recognition behavior during interactions. **(A)** Experimental timeline for SIT with behavioral closed-loop optogenetic manipulation. **(B)** Representative heatmaps of the track during SN-trials in unstressed dams expressing eNPHR. **(C)** Optogenetic inhibition during interaction, but not during exploration, in unstressed dams decreased total interaction time and interaction time per visit with novel mice, but not the number of visits (two-way mixed ANOVA, Wilcoxon signed-rank test). **(D)** Representative heatmaps of the track during SN-trials in stressed dams expressing Chrimson. **(E)** Optogenetic activation during interaction, but not during exploration, in stressed dams increased total interaction time and interaction time per visit with novel mice, but not the number of visits (two-way mixed ANOVA). **(F)** Optogenetic activation during interaction in stressed dams increased and decreased the fractions of novel-excited and novel-suppressed neurons in PrL, respectively. Optogenetic inhibition during interaction in unstressed dams decreased and increased the fractions of novel-excited and novel-suppressed neurons in PrL, respectively. These changes were not observed with optogenetic manipulation during exploration (Chi-squared test). For ANOVAs, * indicates statistical significance for *post hoc* Bonferroni comparisons. * = *p* < 0.05, ** = *p* < 0.01. *ns* = non-significant (*p* > 0.05). All data are represented as mean ± SEM.

### Role of GR in the AI-PrL pathway in postpartum social behavior

Previously, we had shown that stressed dams exhibited an aberrantly sustained elevation of corticosterone during the postpartum period ^17^. This elevation was not accompanied by changes in glucocorticoid receptor (GR) expression in PrL and AI, although there was higher GR expression in AI compared to PrL (Supplementary Fig. 14). In the present study, we have shown how the AI-PrL pathway mediated the effects of SILA on changes in postpartum social novelty behavior. Based on these findings, we further examined if enhanced corticosterone signaling in the AI-PrL pathway led to social behavioral changes in stressed dams in the postpartum period. We applied the Cre recombinase dependent on GFP (CRE-DOG) method to mice floxed with the GR gene (GR^fl/fl^ mice). By combining the GFP-dependent Cre recombinase system and AAV-mediated retrograde tracing, pathway-specific GR knock-out (GR-KO) mice were successfully generated ^34–36^. The CRE-DOG method, in which GFP-binding proteins were used for the molecular assembly of Cre recombinase on a GFP scaffold, allowed us to express Cre recombinase in a GFP-dependent manner (Fig. 5A). We validated that Cre recombinase was successfully expressed in the AI-PrL pathway when AAV-retro-CaMKIIα-EGFP and a mix of CRE-DOG viruses (AAV-EF1a-N-Cretrcint G and AAV-EF1a-C-Creint G) were injected into PrL and AI, respectively (Supplementary Fig. 15, A to F). As time passed after the injections, a gradual increase in the expression level of EGFP was observed in AI, accompanied by a gradual decrease in the expression level of GR in AI neurons expressing EGFP (Fig. 5, B and C). Such expression changes were not observed when AAV-retro-CaMKIIα-mCherry was injected instead of AAV2-retro-CaMKIIα-EGFP as a control virus (Supplementary Fig. 16, A to C).

**Fig. 5.**
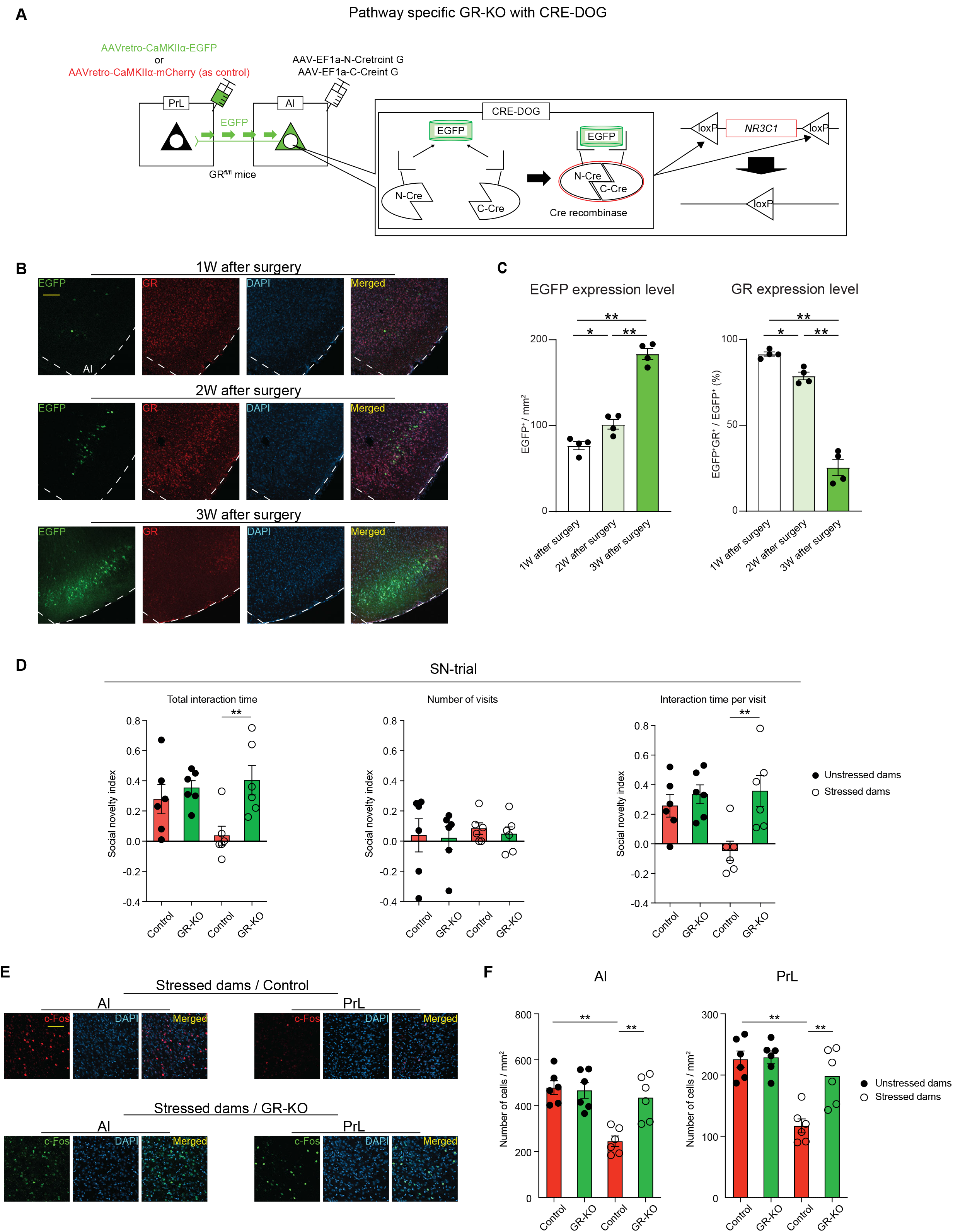
Activation of GR signaling in the AI-PrL pathway played a causal role in postpartum PrL dysfunction and subsequent behavioral changes in social novelty recognition. **(A)** Scheme of AI-PrL pathway specific GR-KO using the CRE-DOG method. **(B)** Representative images of EGFP^+^, GR^+^, DAPI, and colocalized cells in AI of GR^fl/fl^ mice with CRE-DOG. Scale bar, 100 µm. **(C)** GR deletion progressed over time along with expression levels of EGFP (one-way ANOVA, Welch’s ANOVA). **(D)** AI-PrL pathway specific GR-KO ameliorated SILA-induced behavioral changes in SN-trials (two-way ANOVA, Mann Whitney U test). **(E)** Representative images of c-Fos ^+^, DAPI, and colocalized cells in AI and PrL of stressed dams with or without AI-PrL pathway specific GR-KO. Scale bar, 50 μm. **(F)** AI-PrL pathway specific GR-KO normalized the decreased neural activity of AI and PrL in stressed dams (two-way ANOVA).* indicates statistical significance for *post hoc* Bonferroni comparisons. * = *p* < 0.05, ** = *p* < 0.01. All data are represented as mean ± SEM.

Deletion of GR in the AI-PrL pathway ameliorated the behavioral changes in SN-trials, but not S-trials, in stressed dams (Fig. 5D and Supplementary Fig. 16D). AI-PrL pathway-specific GR-KO also normalized the reduced c-Fos immunoreactivity after SN-trials in AI and PrL in stressed dams (Fig. 5, E and F). These findings suggested that activation of GR signaling in the AI neurons projecting to PrL might play a causal role in PrL dysfunction and subsequent behavioral changes in social novelty recognition in the postpartum period.

## Discussion

In the present study, we have found that adolescent psychosocial stress, in conjunction with pregnancy and delivery, leads to hypofunction of the AI-PrL pathway, which in turn alters activity patterns in PrL, resulting in behavioral changes in social novelty recognition. Appropriate social behavior is crucial for the mother to monitor and interpret social signals from others, and behave safely for herself and her children in their new living environment ^5, 37^. Adolescent psychosocial stress may alter behavior during social novelty recognition by decreasing the ability of mothers to recognize social cues via hypofunction of the AI-PrL pathway. Consistent with previous reports ^38–40^, ROC analysis in this study has shown heterogeneity of PrL neurons. Among them, SILA induced postpartum hypofunction of the AI-PrL pathway and affected a specific cell population, namely stable neurons as assessed by *in vivo* calcium imaging. On the other hand, we need to acknowledge the possibility that dynamic neurons are also affected by the AI-PrL pathway. Nevertheless, we believe that the idea that hypofunction of AI-PrL pathway affected stable neurons is reasonable because all analyses in the present study consistently support it.

Although measuring calcium activity and molecular information from the same cells is challenging, elucidating the genetic and molecular features, including the expression level of GR, of these stable neurons would deepen our understanding of their causal role in social novelty recognition. The recently developed calcium and RNA multiplexed activity imaging techniques combined with two-photon calcium imaging in freely moving animals may be used to integrate information from calcium activity and molecular features, and contribute to address this question ^41, 42^. Optogenetic manipulation of the AI-PrL pathway during interaction is sufficient to reproduce or ameliorate the postpartum behavioral changes induced by SILA. PrL is known to be the hub of social behavior and there are various neural circuits involving PrL ^18–20^. Our study has revealed that the AI-PrL pathway contributes to the recognition of novelty of other mice, which is one aspect of social behavior. Exploring upstream and downstream contributions of the AI-PrL pathway would facilitate our understanding of SILA-induced social behavioral changes and the nature of social behavior.

We have elucidated the causal role of the GR-mediated AI-PrL pathway in postpartum behavioral changes during social novelty recognition. Although GR is also well expressed in PrL, GR-KO in the AI-PrL pathway is enough to ameliorate the neuronal activity change in PrL of stressed dams. SILA-induced, aberrantly sustained elevation of corticosterone, which is observed only during the postpartum period ^17^, is important for social behavioral changes in stressed dams. It may account for why SILA affects social behavior only during the postpartum period. Although SILA obstructs normalization of the physiological increase in corticosterone during the postpartum period, SILA itself does not cause corticosterone elevation ^17^. We speculate therefore that SILA-induced social behavioral changes are observed only during the postpartum period^17^. Further research on how GR signaling specifically regulates the AI-PrL pathway function and recognition of social cues would facilitate our mechanistic understanding of the pathological trajectory of adolescent stress leading to abnormal social behavior in the postpartum period.

## Methods

### Mice

C57BL/6J (B6J; JAX 000664), B6;129S-*Slc17a7^tm^*^1^.^1^(cre)*^Hze^*/J (Vglut1-Cre; 023527), B6.Cg-*Gt(ROSA)26Sor^tm^*^14^*^(CAG-tdTomato)Hze^*/J (Ai14; 007914), and B6.Cg-*Nr3c1^tm^*^1^*^.1Jda^*/J (GR^fl/fl^; 021021) mice were purchased from the Jackson Laboratory. All mice used in this study were backcrossed (more than ten generations) to the B6J genetic line. For SILA exposure, virgin, healthy female mice were isolated from five to eight weeks of age. Each female was then mated with a healthy B6J male mouse at eight weeks of age, and gave birth to pups (Fig. 1A). Isolation consisted of no interaction with other mice and confinement to opaque, wire-topped polypropylene cages, whereas group-housed mice were kept in clear, wire-topped plastic cages (18 × 28 × 14 cm). All mice were maintained under a controlled environment (23 ± 3°C; 40 ± 5% humidity; light and dark cycles started at 6 am and 6 pm, respectively) with ad libitum access to food and water. All experimental procedures were performed in accordance with the National Institutes of Health Guidelines for the Care and Use of Laboratory Animals, and under animal protocols approved by the Institutional Animal Care and Use Committees at the University of Alabama at Birmingham.

### Viruses

AAV1-Syn-Flex-GCaMP6f-WPRE-SV40 was a gift from Douglas Kim and the GENIE Project (Addgene viral prep # 100833-AAV1; http://n2t.net/addgene:100833; RRID: Addgene_100833)^43^. AAVretro-hSyn-EGFP was a gift from Bryan Roth (Addgene viral prep # 114469-AAVrg; http://n2t.net/addgene:50465; RRID: Addgene_50465). AAVretro-CaMKIIa-mCherry was a gift from Karl Deisseroth (Addgene viral prep # 114469-AAVrg; http://n2t.net/addgene:114469; RRID: Addgene_114469). AAV1-EF1a-N-CretrcintG and AAV1-EF1a-C-CreintG were gifts from Connie Cepko (Addgene viral prep # 69570-AAV1 and 69571-AAV1; http://n2t.net/addgene:69570 and http://n2t.net/addgene:69571; RRID: Addgene_69570 and Addgene_69571) ^34^. AAVretro-hSyn-Cre-WPRE-hGH was a gift from James M. Wilson (Addgene viral prep # 105553-AAV1; http://n2t.net/addgene:105553; RRID: Addgene_105553). AAV5-Syn-Flex-ChrimsonR-tdTomato, AAV5-Syn-Flex-tdTomato, AAV5-EF1a-DIO-eNpHR3.0-mCherry, and AAV5-EF1a-DIO-mCherry were purchased through the Vector Core at the University of North Carolina at Chapel Hill. AAVretro-CAMKIIa-EGFP was produced at the Vector Core at the University of Alabama at Birmingham.

### Surgical procedures

All surgery was performed under aseptic conditions, and body temperature was maintained with a heating pad. Catoprofen [Rimadyl, Zoetis; 5 mg/kg body weight (BW)] was administrated subcutaneously as a preoperative pain medication. Mice were anesthetized using isoflurane mixed with oxygen (5% for induction, 2-2.5% for maintenance, 1 L/min oxygen flow rate). Eyes were protected with ophthalmic ointment (LubriFresh P.M., Major Pharmaceuticals).

For injections into specific brain regions, the following bregma coordinates were used: PrL, +0.4 mm medial-lateral (ML), +2.1 mm anterior-posterior (AP), -2.2 mm dorsal-ventral (DV); AI, +2.4 mm ML, +2.22 mm AP, -2.95 mm DV; BLA, +2.7 mm ML, -1.12 mm AP, -4.9 mm DV. Injections were performed using a 33-gauge beveled microinjection needle with a 10 µL micro syringe (Nanofil; World Precision Instruments), delivering viruses at a rate of 10 nL/min using a microsyringe pump (UMP3; World Precision Instruments) and controller (Micro2T; World Precision Instruments). After completion of the injections, 15 min were allowed to pass before the needle was slowly withdrawn. As a postoperative pain medication, Catoprofen (5 mg/kg BW) was administrated subcutaneously every 24 hours for three days.

For tracing projections to PrL, AAVretro-hSyn-EGFP (200 nL) was injected into PrL of male and female virgin B6J mice (seven to ten weeks old at the time of virus injections). For anterograde tracing of the AI-PrL pathway, AAV5-Flex-tdTomato (200 nL) was injected into AI or PrL of female virgin Vglut1-Cre mice (seven to ten weeks old at the time of virus injections). For tracing collateral projections of the AI-PrL pathway, AAVrg-hSyn-Cre-WPRE-hGH (200 nL) and AAV5-Syn-Flex-tdTomato (200 nL) were injected into PrL and AI of female virgin B6J mice (seven to ten weeks old at the time of virus injection), respectively.

For optogenetic manipulation of the AI-PrL or BLA-PrL pathway, virus (AAV5-Syn-Flex-ChrimsonR-tdTomato or AAV5-Syn-Flex-tdTomato for stressed dam group, AAV5-EF1a-DIO-eNpHR3.0-mCherry or AAV5-EF1a-DIO-mCherry for unstressed dam group and virgin group, 200 nL each) was unilaterally injected into AI or BLA of virgin Vglut1-Cre mice (five weeks old at the time of virus injections for stressed and unstressed dam groups, eight weeks old at the time of virus injection for virgin group).

Surgery for *in vivo* microendoscopic calcium imaging of PrL was conducted just after surgery for optogenetics. A 1.1 mm-diameter craniotomy was first made at the coordinates, +0.6 mm ML, +2.1 mm AP. After aspiration of brain tissue above PrL using a 30-gauge blunted needle, AAV1-Syn-Flex-GCaMP6f-WPRE-SV40 (200 nL) was injected into PrL. The gradient refractive index lens (GRIN lens; GLP-1040, Inscopix) was then implanted into the dorsal region of PrL (+1.25 mm ML, +2.1 mm AP, -1.8 mm DV) and secured to the skull using dental cement (Metabond S380, Parkell). Following three weeks of virus incubation, baseplates (Inscopix) were cemented around the lens to support the connection of the miniaturized microscope.

For CRE-DOG studies, AAVretro-CaMKIIa-EGFP and a mix of CRE-DOG viruses (AAV1-EF1a-N-Cretrcint G and AAV-EF1a-C-Creint G) were bilaterally injected into PrL and AI of GR^fl/fl^, B6J or Ai14 mice (eight weeks old). AAV1-Flex-tdTomato was also injected into AI for a validation study with B6J mice.

For all experiments involving viral or tracer injections, animals containing mistargeted injection(s) were excluded after histological verification.

### Behavioral tests

Social interaction test (SIT). SIT was conducted as previously described with minor modifications ^17, 44^. Of note, the data in **Fig. 1C and E** in the present study, in which the experiments were conducted on two consecutive days, were different to previous data, in which the experiments were conducted on a single day ^17^. The subject mouse was introduced into a chamber apparatus (60 cm × 40 cm × 35 cm), and was allowed to habituate to the chamber for 30 min for three consecutive days before testing. During habituation, the subject mouse for SIT with calcium imaging was connected to a plastic “dummy” microscope for training. In the S-trial, the subject mouse encountered a novel mouse in a wire cage and an empty wire cage in a chamber for 10 min. In the SN-trial, the subject mouse encountered a familiar mouse, which was co-housed with the subject animal on the day before the SN-trial, and a novel mouse in the wire cage. The parameters analyzed were time spent sniffing each cage (total interaction time), number of visits to each cage, and interaction time per visit to the cage. Measurements were taken using the Ethovision XT 15 software (Noldus). Age- and sex-matched unstressed mice were used as familiar and novel mice. Sociability and social novelty indexes were calculated as follows: (time interacting with mouse or novel mouse cage – time interacting with empty or familiar mouse cage) / (time interacting with mouse or novel mouse cage + time interacting with empty or familiar mouse cage), meaning that indexes of 0 indicated no preference, while positive indexes indicated increased sociability or social novelty behavior, and negative indexes indicated social avoidance or deficits in social novelty behavior.

For SIT without calcium imaging, S- and SN-trials were conducted on postpartum days seven and eight (Fig. 1A). For SIT with calcium imaging, S-trials were conducted on postpartum days seven and eight, and SN-trials were conducted on postpartum days nine and ten. The trials on one of these two days were conducted with optogenetic stimulation and counterbalanced as described in Supplementary Figures S3C and S12A.

Novel objective recognition test (NOR). NOR in Supplementary Figures 12 A-D was conducted on postpartum days seven, 11/12, or 12/13 as previously described with minor modifications ^46, 47^. The trials on one of the two days were conducted with the light stimulation and counterbalanced. Each trial consisted of three phases: habituation, training, and test phases (10 min each). During the habituation phase, the subject mouse was introduced into a chamber apparatus (60 cm × 40 cm × 35 cm) and was allowed to habituate for 10 min. During the training phase, two identical objects were introduced into the chamber apparatus. In the test phase, one of two objects was replaced by a novel object (Supplementary Fig. 12 E). For NOR in Supplementary Figures 12H, only one object was introduced into the chamber apparatus during the training phase. This modification was made to be analogous to the method for SIT. The parameters analyzed were time spent sniffing the object (total interaction time), number of visits to each object, and interaction time per visit to the object. The time spent sniffing each object was analyzed using the Ethovision XT 15 software. The discrimination index was calculated as follows: (time sniffing novel object – time sniffing familiar object / (time sniffing novel object + time sniffing familiar object), meaning that indexes of 0 indicated no recognition of a novel object, while positive indexes indicated increased novel object recognition and negative indexes indicated deficits in novel object recognition.

Real-time place preference test (RTPP). RTPP was conducted on postpartum day 13 or 14, as previously described with minor modifications ^45^. The subject mouse was introduced into a chamber apparatus (60 cm × 40 cm × 35 cm) and was allowed to habituate for 10 min. Each test consisted of two consecutive 15-min sessions. In the first session, mice were allowed to freely explore the two compartments for 15 min, during which entering into one half of the chamber triggered the light stimulation. Exiting the stimulated chamber immediately terminated the light stimulation. During the second session, the opposite side was paired with optogenetic stimulation. The results of these two sessions were combined, and time spent in the unstimulated and stimulated sides was evaluated using the Ethovision XT 15 software.

### Optogenetic manipulation

Optogenetic manipulation was performed using nVoke 2.0 (Inscopix). For optogenetic activation, 20 Hz, 60-pulse trains (5 ms each) of LED light (620 ± 30 nm, 20 mW/mm^2^) were initiated every 30 s during the light on epoch ^48^. For optogenetic inhibition, constant LED light (620 ± 30 nm, 5 mW/mm^2^) was delivered during the light on epoch ^49^. The EthoVision XT 15 software and a mini-IO box system (Noldus) were used to record live tracking of mice, and to trigger the laser on during specific behaviors.

### *In vivo* microendoscopic calcium imaging and analysis

Calcium imaging was performed using a head-mounted microscope to image through a chronically implanted GRIN lens placed above PrL with nVoke 2.0 (Inscopix). Imaging and analyses were conducted with the Inscopix Data Processing Software (IDPS, Inscopix), as previously described with minor modifications and custom MATLAB scripts (MathWorks) ^29–32, 48^. GCaMP6f emission signals were acquired continuously at a frame rate of 20 Hz with blue LED light (455 ± 8 nm, power of 10-60%, analog gain of 1), and spatially down-sampled (2x) prior to motion correction. The motion-corrected video data were then converted to [fluorescence (F) - background fluorescence (F_0_) / F_0_] (ΔF / F_0_), using the mean projection images of the entire movie as F_0_. Calcium signals arising from individual regions of interest (ROIs, that is, cells) were identified using principal and independent component analyses (PCA / ICA), as previously described ^50^. Identified ROIs were then screened for neuronal morphology, and only accepted if they included an area between seven to 70 pixels. The accepted ROI filters were then reapplied to the motion-corrected videos to extract (F - F_0_) / F_0_ traces for each ROI. To match individual neurons across recording sessions, we used the longitudinal registration function of IDPS.

To evaluate the effects of optogenetic manipulation of the AI-PrL pathway on PrL neurons, the modulation index and relative peak amplitude were calculated with calcium imaging in the home cage, as previously described ^29^. Calcium imaging was performed in the home cage for 10 min with and without light stimulation (5 min each, counterbalanced). Calcium transients were detected using the Detect Events function of IDPS (Event Threshold Factor: 4.00, Event Smallest Decay Time: 0.20 sec). The modulation index was evaluated by calculating the number of calcium transients occurring with (Light On) / (Light On + Light Off). Relative peak amplitude was evaluated by calculating the average peak amplitude occurring with (Light On) / (Light Off). Responses of single neurons during behavioral events in SIT and NOR were quantified using an ROC analysis, which is commonly used to characterize neuronal response in calcium imaging data ^30–32^. The definitions of PrL activity patterns are provided in Supplementary Table 2.

Upon application of a binary threshold to ΔF / F_0_ signals and comparison with a binary event vector denoting behavior (“interaction with a mouse” and “sniffing an empty cage” for S-trials, “interaction with a familiar mouse” and “interaction with a novel mouse” for SN-trials, “sniffing familiar objects” and “sniffing novel objects” for NOR), the true positive rate (TPR) and false positive rate (FPR) over all time points were used for measuring behavioral event detection based on neural activity. An ROC curve was generated by plotting TPR against FPR over a binary threshold range of minimum and maximum values of the neural signal, representing the behavioral event detectability of the neural signal at each threshold value. Based on the area under the ROC curve (auROC) value, we classified PrL neurons into three mutually exclusive categories (excited, suppressed, and unresponsive) for each behavior. For each neuron / behavior category, the observed auROC was compared to a null distribution of 1,000 auROC values generated from constructing ROC curves over randomly permuted calcium signals (that is, traces that were circularly permuted using a random time shift). A neuron was considered significantly responsive if its auROC value exceeded the 95th percentile of the random distribution (auROC < 2.5th percentile for suppressed responses, auROC > 97.5th percentile for excited responses). It should be noted that suppressed responses that were defined with this method did not necessarily exhibit an immediate decrease of activity during the corresponding behavior, but displayed an overall negative correlation with the corresponding behavior.

To analyze the pattern of PrL activity changes during the S- or SN-trials on two consecutive days, every single cell was assigned to one cell of a 3 x 3 matrix table based on the results of the ROC analysis. The differences between two matrix tables were evaluated with a chi-squared test, and AR was used to measure the contribution of each cell to significant differences. Stable or dynamic neurons were defined only by the results from animals with control viruses, not opsins. For the interpretation of the data from animals with opsins, we compared them with the data from animals with control viruses and evaluated AR values.

### Immunohistochemistry

Mice were deeply anesthetized with isoflurane, and subsequently intracardially with 1x PBS (Fisher Scientific, BP3991) followed by 4% paraformaldehyde (PFA) (Electron Microscopy Science, 15714-S). The brains were extracted and post-fixed overnight in 4% PFA at 4°C. Coronal sections were obtained with a cryostat at 30 μm. The sections were blocked with 5% normal goat serum containing 0.1% Triton-X 100 (Sigma-Aldrich, 93443) in 1x PBS for 1 h, and then incubated with the following primary antibodies (12 h at 4°C): anti-Vglut1 (1:200, rabbit, Abcam, ab227805), anti-Vgat (1:200, rabbit, Millipore, AB5062P), and anti-c-Fos (1:1000, rabbit, Cell Signaling Technology, 2250 or 1:1000, mouse, Abcam, ab208942). After rinsing, sections were incubated with the following fluorophore-conjugated secondary antibodies for 2 h at room temperature: anti-rabbit Alexa Fluor 488 (1:400, goat, Invitrogen, A11008), anti-rabbit Alexa Fluor 568 (1:400, goat, Invitrogen, A11011), anti-mouse Alexa Fluor 594 (1:400, goat, Invitrogen, A11032), and anti-mouse Alexa Fluor 488 (1:400, goat, Invitrogen, A11001). For double immunohistochemistry using a mouse antibody, the sections were incubated with goat anti-mouse F(ab) fragment (Abcam, ab6668) for 1 h at room temperature after blocking. Antibodies were diluted in the same block solution.

All immunohistochemistry samples were imaged at every third 30 μm interval using the LSM-800 Airyscan confocal microscope (Zeiss), analyzed using Image J Cell Counter plugin, and assigned to brain areas based on classifications according to the Paxinos Mouse Brain Atlas ^51^, using anatomical landmarks in the sections visualized by DAPI staining and tissue autofluorescence, as previously described with minor modifications ^52^. We also confirmed manually if the elongated signals were not included for the results.

For the whole-brain retrograde tracing study, mice were perfused at four weeks following virus injections, and EGFP^+^, EGFP^+^Vglut1^+^, or EGFP^+^Vgat^+^ cells were counted. For the c-Fos study, mice were perfused 90 min after the SN-trial, and c-Fos^+^, Vglut1^+^c-Fos^+^, or Vgat^+^c-Fos^+^ cells were counted. For the dual viral and anterograde tracing studies, mice were perfused at four weeks after virus injections. For the quantification of fluorescence intensity as a proxy for the terminal density of the AI-PrL pathway, 100 (w) × 200 (h) μm sections across the PrL layer based on DAPI density / morphology were analyzed using Fiji, as previously described ^48^. For systematic examination of c-Fos immunoreactivity after the SN-trial, we counted the number of c-Fos^+^ cells and normalized by the mean of those in unstressed dams for each region to evaluate the fold changes ^53^.

For the CRE-DOG validation studies, mice were perfused at one, two, or three weeks after virus injections. EGFP^+^, tdTomato^+^EGFP^+^, EGFP^+^GR^+^, mCherry^+^, or mCherry^+^GR^+^ cells were counted.

### Statistics and reproducibility

All statistical analyses were conducted using SPSS 28 (IBM) and MATLAB R2020a, b, and 2021a (MathWorks). Experiments were randomized whenever possible. All behavioral, imaging, and optogenetics experiments were replicated in multiple subject animals with similar results (the exact numbers of animals and trials for each experiment are provided in **Supplementary Table 3**). For continuous data, normality and homogeneity of variances were tested using the Shapiro-Wilk normality test and Levene’s test, respectively. For comparisons between two groups, we used Student’s *t* tests or Paired Student’s *t* tests. ANOVAs and mixed ANOVAs (meaning at least one variable was a paired measure) were performed as appropriate according to the statistical design of each type of experiment, followed by relevant pairwise comparisons using Bonferroni adjustments. In some cases, data that failed only in the homogeneity of variances were analyzed using Welch *t* tests or Welch ANOVA. When data failed in the assumptions of normality and/or homogeneity of variances, we performed pairwise comparisons using non-parametric tests. The Wilcoxon signed-rank test was used for paired samples, and Mann Whitney U test was used for independent samples. For categorical data, we performed Chi-Squared tests.

Statistical significance was established by *p-*values < 0.05. We also reported statistical tendencies in some cases, defined as *p* values < 0.10. Values of η^2^ and *r* were included in the interpretation of results when relevant to discussing effect sizes. Detailed information for statistical analyses was provided in **Supplementary Table 3**, organized by Figures, with reports of sample sizes, factors used in analysis, type of tests employed for each data set, results from Shapiro-Wilk and Levene’s tests, F values, *U* values, z values, and η^2^ and *r* values for effect sizes. All exact *p* values were also reported.

## Acknowledgements

**Acknowledgments:** We thank Ms. Dania Mallah for assisting with the calcium imaging analysis; members of the Niwa laboratory and Kano laboratory and Dr. Akira Sawa for helpful discussions; Dr. Nao J. Gamo, Dr. J. Andrew Hardaway, and Ms. Phoebe Garcia for their critical reading; Dr. Patrick Stemkowski, Dr. Vardhan Dani, and Dr. Waylin Yu of Inscopix for technical assistance; the Civitan Cellular Imaging Core for use of the Airyscan confocal microscope; and the UAB Vector and Virus Core for virus preparation. This work was supported by the National Institutes of Health (NIH) R01MH-116869 (M.N.), NIH R21MH-128708 (S.K.), and UAB Psychiatry startup funds (M.N.). K.K. was supported by the Takeda Science Foundation Fellowship Program for Young Japanese MDs & PhDs Studying Abroad.

**Author contributions:** MN conceived, designed, and supervised the project with inputs from SK. KK performed the experiments. KK and JFO analyzed the data. KK and MN wrote the first manuscript with inputs from SK. All authors revised and edited the paper.

## Competing interests

The authors declare no competing interests.

## Figure legends

## Notes

### Competing Interest Statement

The authors have declared no competing interest.

### Summary of Updates

Updated the statistical results. Add one co-author.

