## Supplemental material for "Adolescent stress impairs postpartum social behavior via anterior insula-prelimbic pathway"

**This PDF file includes:**

Supplementary Figs. 1 to 16

Supplementary Tables 1 to 3

Supplementary Fig. 1

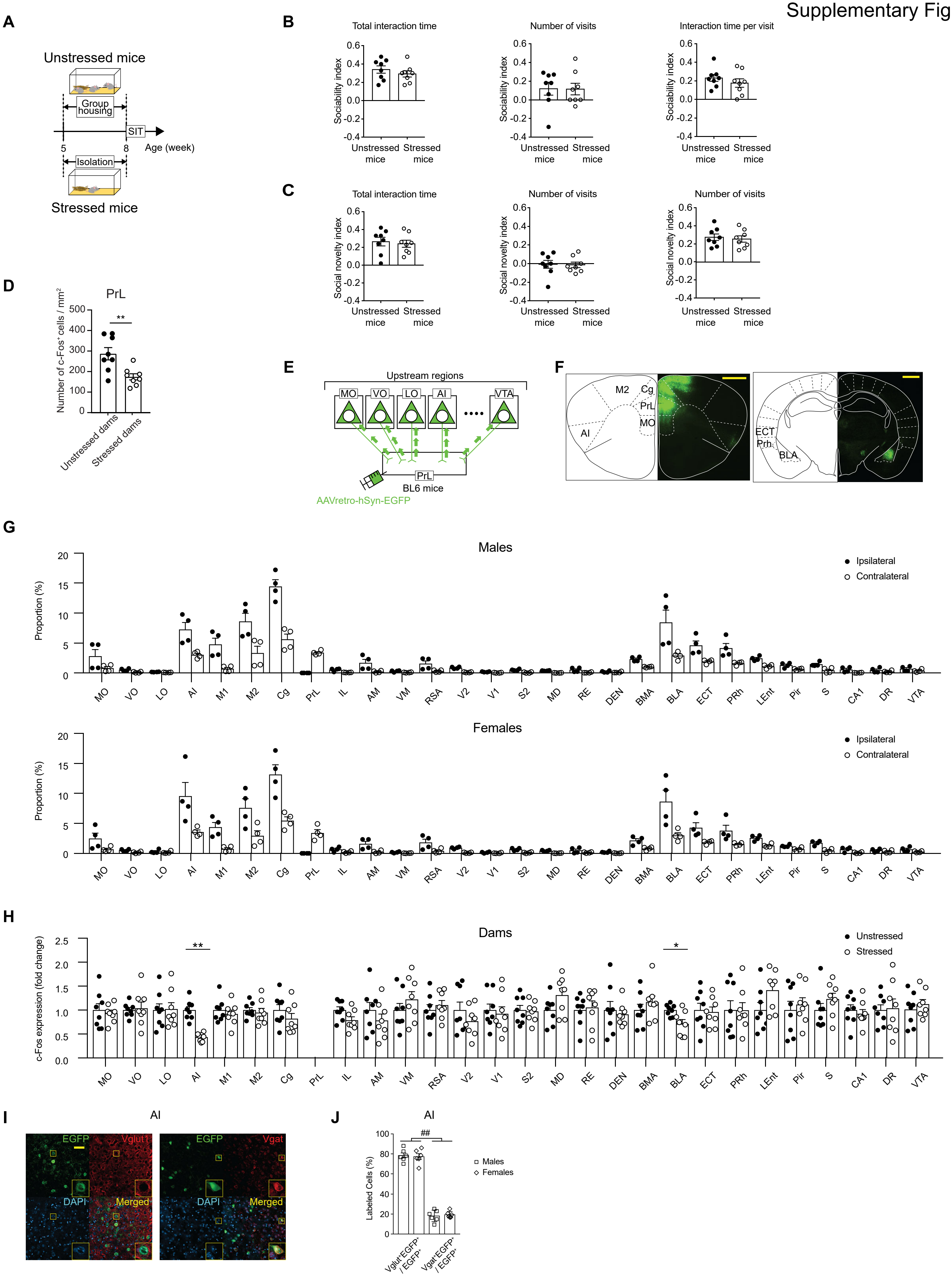

**Supplementary Fig. 1. SILA led to reduced postpartum neuronal activity in PrL and AI after SN-trial.** (**A**) Experimental timeline of the social interaction test (SIT) immediately after SILA. (**B** and **C**) No behavioral changes were observed during S- and SN-trials immediately after SILA (Student's *t* test). (**D**) c-Fos expression after the SN-trial was decreased in PrL of stressed dams in comparison to unstressed dams (Student's *t* test). (**E**) Scheme of retrograde tracing in BL6 mice injected with AAVretro-hSyn-EGFP into PrL. (**F**) Representative images showing labeled neurons in cingulate cortex (Cg), second motor cortex (M2), AI, and BLA. Scale bars, 1 mm. (**G**) Whole-brain quantification of projections to PrL, in male and female mice. For definitions of the abbreviations, see **Supplementary Table 1** (Multiple Paired Student's *t* test and Wilcoxon signed-rank test. Statistical significance not shown for clarity. See **Supplementary Table 3** for *p* values for each region analyzed). (**H**) c-Fos expression levels in dams upstream regions of PrL after SN-trials. c-Fos expression was decreased only in AI and BLA of stressed dams in comparison to unstressed dams. For definitions of the abbreviations, see **Supplementary Table 1** (Multiple Student's *t* test and Mann-Whitney U test. $p$ values <0.10 are also indicated). (**I**) Representative images of $EGFP^+$, $Vglut1^+$ or $Vgat^+$, DAPI, and colocalized cells in AI of mice injected with AAVretro-hSyn-EGFP into PrL. Scale bar, 50 µm. (**J**) AI-PrL pathway mainly consisted of $Vglut1^+$ neurons, not $Vgat^+$ neurons (two-way mixed ANOVA). All data are represented as mean ± SEM. For ANOVAs, * indicates statistical significance for *post hoc* Bonferroni comparisons, and # indicates statistical significance for the main effect. * = $p < 0.05$, ** = $p < 0.01$, ## = $p < 0.01$.

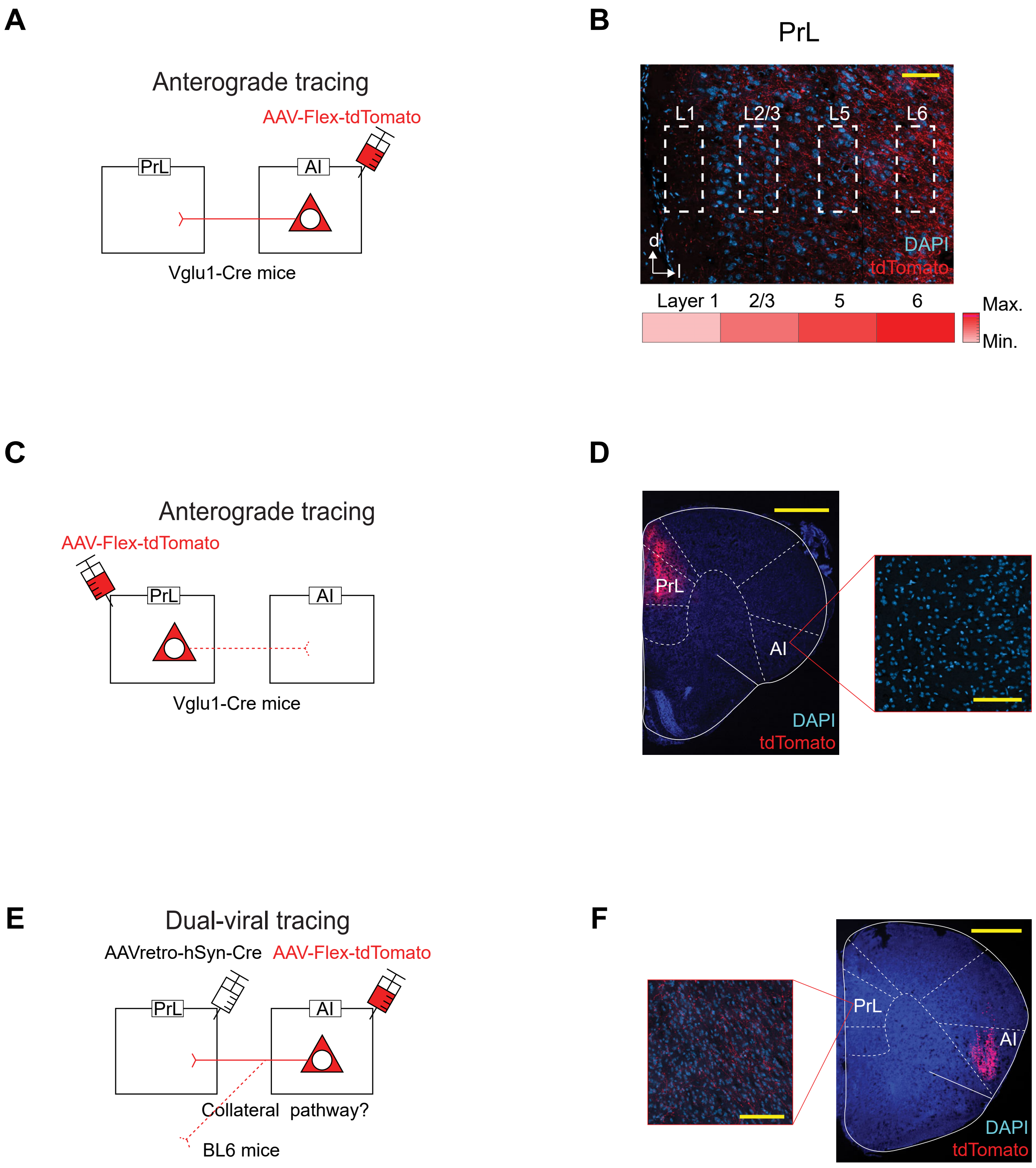
A
Anterograde tracing
AAV-Flex-tdTomato
PrL
AI
Vglu1-Cre mice
B
PrL
L1
L2/3
L5
L6
d
l
DAPI
tdTomato
Layer 1
2/3
5
6
Max.
Min.
C
Anterograde tracing
AAV-Flex-tdTomato
PrL
AI
Vglu1-Cre mice
D
PrL
AI
DAPI
tdTomato
E
Dual-viral tracing
AAVretro-hSyn-Cre
AAV-Flex-tdTomato
PrL
AI
Collateral pathway?
BL6 mice
F
PrL
AI
DAPI
tdTomato

**Supplementary Fig. 2. AI-PrL pathway was a unidirectional pathway from AI to PrL with no collateral projections.** (**A**) Scheme of anterograde tracing in Vglut1-Cre female mice injected with AAV-Flex-tdTomato into AI. (**B**) Representative image showing expression of tdTomato at terminals of the AI-PrL pathway in PrL. Terminals of the AI-PrL pathway had the highest density in deep layers. Scale bar, 100 μm. (**C**) Scheme of anterograde tracing in Vglut1-Cre mice injected with AAV-Flex-tdTomato into PrL. (**D**) Representative images showing expression of tdTomato in PrL, but not in AI. Scale bars, 1 mm and 100 μm. (**E**) Scheme of dual-virus tracing in BL6 female mice injected with AAV-Flex-tdTomato and AAVretro-hSyn-Cre into AI and PrL, respectively, to explore the collateral pathway of the AI-PrL pathway. (**F**) Representative images showing expression of tdTomato in only AI and PrL. Scale bars, 100 μm and 1 mm.

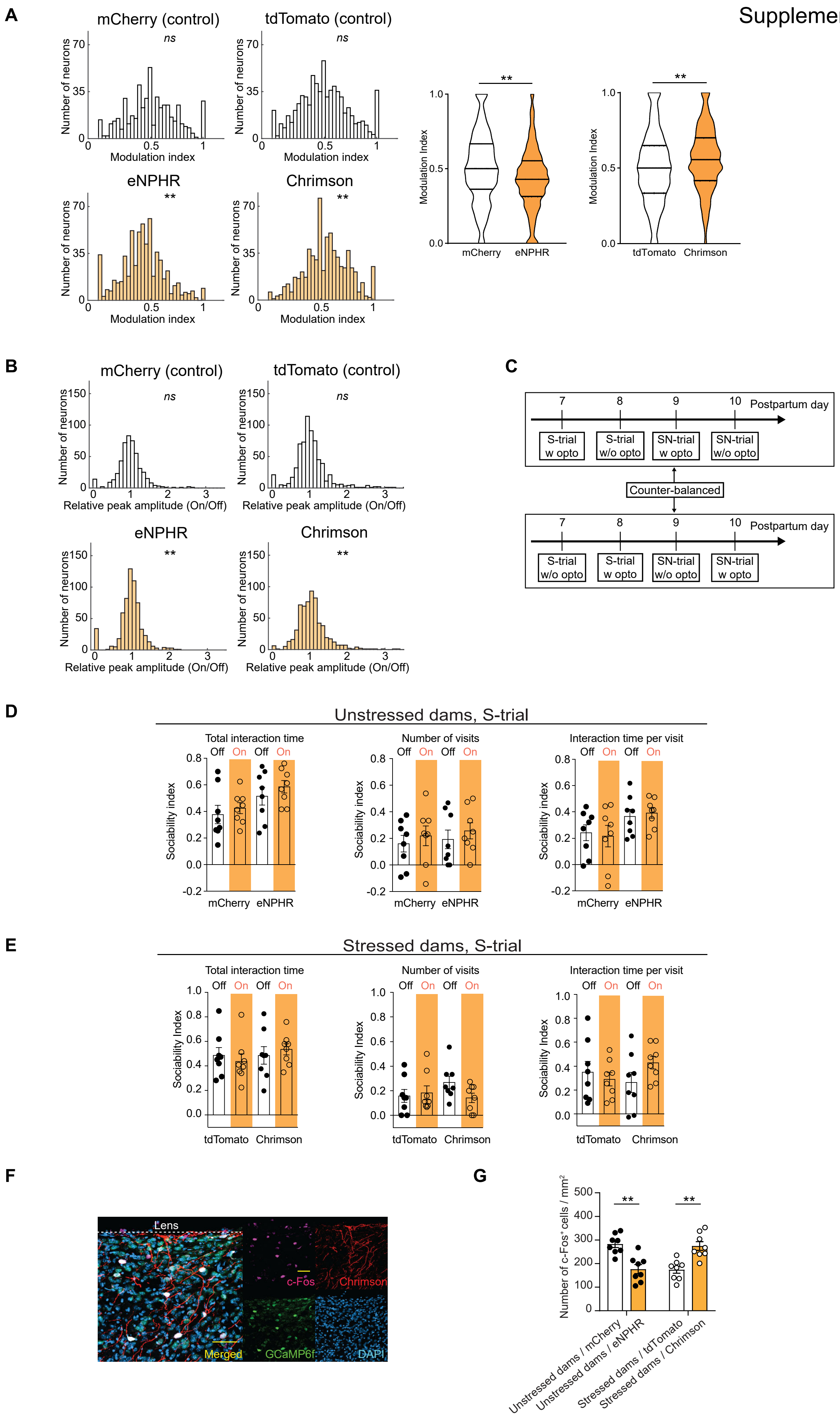
A
mCherry (control)
tdTomato (control)
eNPHR
Chrimson
ns
**
Number of neurons
Modulation index
Modulation Index
mCherry
eNPHR
tdTomato
Chrimson
B
mCherry (control)
tdTomato (control)
eNPHR
Chrimson
Relative peak amplitude (On/Off)
C
Postpartum day
S-trial w opto
S-trial w/o opto
SN-trial w opto
SN-trial w/o opto
Counter-balanced
D
Unstressed dams, S-trial
Total interaction time
Number of visits
Interaction time per visit
Off
On
Sociability index
E
Stressed dams, S-trial
Sociability Index
F
Lens
c-Fos
Chrimson
Merged
GCaMP6f
DAPI
G
Number of c-Fos+ cells / mm2
Unstressed dams / mCherry
Unstressed dams / eNPHR
Stressed dams / tdTomato
Stressed dams / Chrimson

**Supplementary Fig. 3. AI-PrL pathway did not affect sociability in the postpartum period.** (**A**) Modulation index for PrL neurons showed inhibition or activation of calcium transients during optogenetic activation or suppression of the AI-PrL pathway, respectively (Wilcoxon signed-rank test for one sample, $r > 0.25$ indicating small to intermediate effect size). Comparison between control and opsin-expressing group of cells, showing the median ± interquartile range in a violin plot (Mann Whitney U test, $r > 0.10$ indicating small effect size). (**B**) Calcium transient peak amplitudes of PrL were increased or decreased by optogenetic activation or inhibition of AI-PrL pathway, respectively (Wilcoxon signed-rank test for one sample, $r > 0.1$ indicating small effect size). (**C**) Experimental timeline of SIT without and with optogenetic manipulation. (**D** and **E**) Optogenetic manipulation of the AI-PrL pathway did not affect sociability in both stressed and unstressed dams (two-way mixed ANOVA, Wilcoxon signed-rank test). (**F**) Representative images showing expression of GCaMP6f underneath a GRIN lens, pathway from AI expressing Chrimson, and labeled c-Fos$^{+}$ cells in PrL. Scale bars, 50 µm. (**G**) c-Fos expression in PrL was decreased by optogenetic inhibition of the AI-PrL pathway in unstressed dams, and increased by optogenetic activation of the AI-PrL pathway in stressed dams (two-way ANOVA, $p < 0.01$ for interaction). Except for the modulation indexes in **Supplementary Figure 3A**, data are represented as mean ± SEM. For ANOVAs, * indicates statistical significance for *post hoc* Bonferroni comparisons. * = $p < 0.05$, ** = $p < 0.01$. *ns* = non-significant ($p > 0.05$).

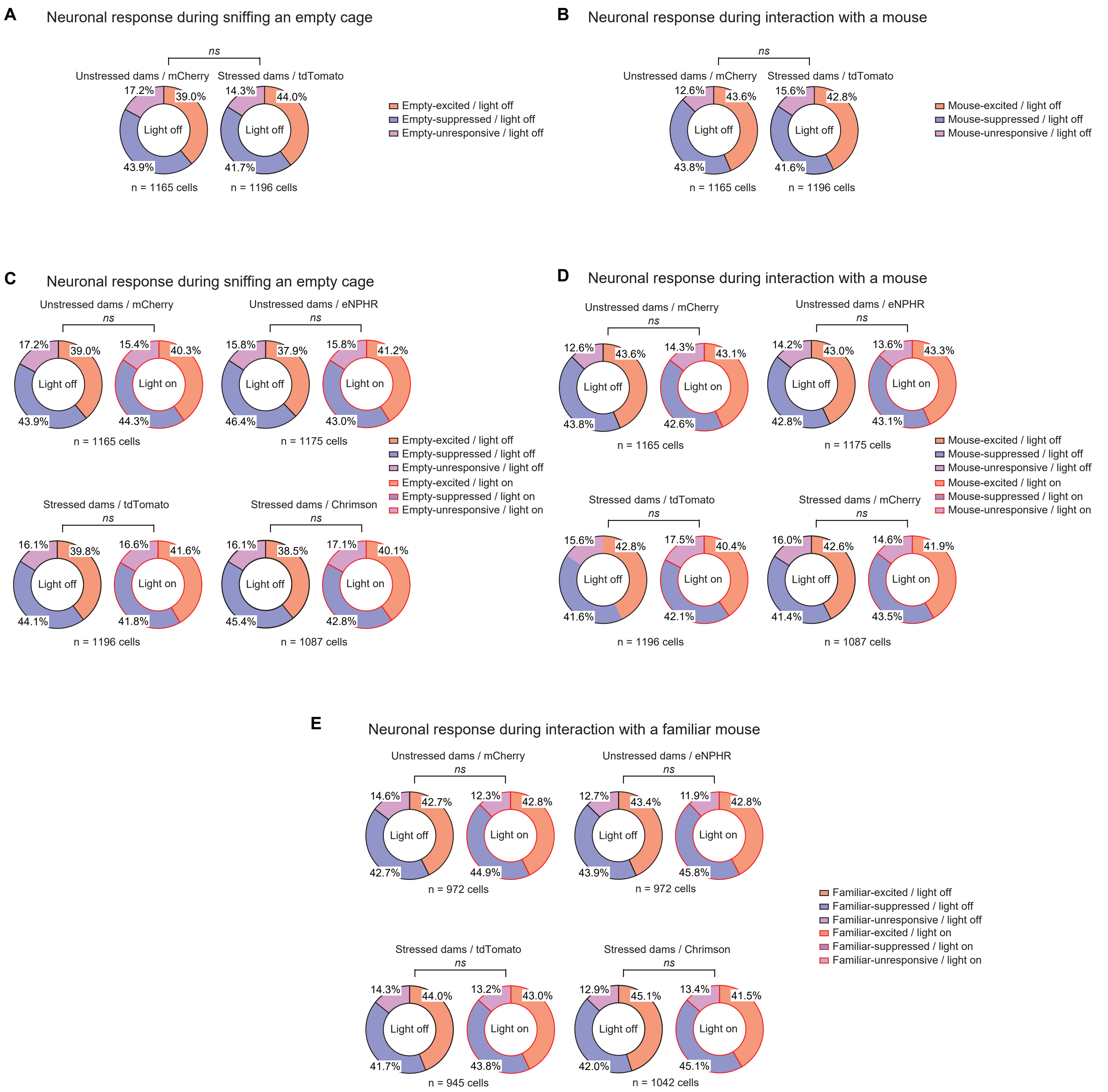
A
Neuronal response during sniffing an empty cage
ns
Unstressed dams / mCherry
Stressed dams / tdTomato
17.2%
39.0%
Light off
43.9%
n = 1165 cells
14.3%
44.0%
Light off
41.7%
n = 1196 cells
Empty-excited / light off
Empty-suppressed / light off
Empty-unresponsive / light off
B
Neuronal response during interaction with a mouse
ns
Unstressed dams / mCherry
Stressed dams / tdTomato
12.6%
43.6%
Light off
43.8%
n = 1165 cells
15.6%
42.8%
Light off
41.6%
n = 1196 cells
Mouse-excited / light off
Mouse-suppressed / light off
Mouse-unresponsive / light off
C
Neuronal response during sniffing an empty cage
Unstressed dams / mCherry
ns
17.2%
39.0%
Light off
43.9%
15.4%
40.3%
Light on
44.3%
n = 1165 cells
Unstressed dams / eNPHR
ns
15.8%
37.9%
Light off
46.4%
15.8%
41.2%
Light on
43.0%
n = 1175 cells
Stressed dams / tdTomato
ns
16.1%
39.8%
Light off
44.1%
16.6%
41.6%
Light on
41.8%
n = 1196 cells
Stressed dams / Chrimson
ns
16.1%
38.5%
Light off
45.4%
17.1%
40.1%
Light on
42.8%
n = 1087 cells
Empty-excited / light off
Empty-suppressed / light off
Empty-unresponsive / light off
Empty-excited / light on
Empty-suppressed / light on
Empty-unresponsive / light on
D
Neuronal response during interaction with a mouse
Unstressed dams / mCherry
ns
12.6%
43.6%
Light off
43.8%
14.3%
43.1%
Light on
42.6%
n = 1165 cells
Unstressed dams / eNPHR
ns
14.2%
43.0%
Light off
42.8%
13.6%
43.3%
Light on
43.1%
n = 1175 cells
Stressed dams / tdTomato
ns
15.6%
42.8%
Light off
41.6%
17.5%
40.4%
Light on
42.1%
n = 1196 cells
Stressed dams / mCherry
ns
16.0%
42.6%
Light off
41.4%
14.6%
41.9%
Light on
43.5%
n = 1087 cells
Mouse-excited / light off
Mouse-suppressed / light off
Mouse-unresponsive / light off
Mouse-excited / light on
Mouse-suppressed / light on
Mouse-unresponsive / light on
E
Neuronal response during interaction with a familiar mouse
Unstressed dams / mCherry
ns
14.6%
42.7%
Light off
42.7%
12.3%
42.8%
Light on
44.9%
n = 972 cells
Unstressed dams / eNPHR
ns
12.7%
43.4%
Light off
43.9%
11.9%
42.8%
Light on
45.8%
n = 972 cells
Stressed dams / tdTomato
ns
14.3%
44.0%
Light off
41.7%
13.2%
43.0%
Light on
43.8%
n = 945 cells
Stressed dams / Chrimson
ns
12.9%
45.1%
Light off
42.0%
13.4%
41.5%
Light on
45.1%
n = 1042 cells
Familiar-excited / light off
Familiar-suppressed / light off
Familiar-unresponsive / light off
Familiar-excited / light on
Familiar-suppressed / light on
Familiar-unresponsive / light on

**Supplementary Fig. 4. AI-PrL pathway did not affect neuronal activity in PrL and sociability in the postpartum period.** (**A** and **B**) PrL activity during S-trials did not show any differences between unstressed and stressed dams expressing control viruses (Chi-squared test). (**C** and **D**) Optogenetic manipulation of the AI-PrL pathway did not affect PrL activity patterns during S-trials in both stressed and unstressed dams (Chi-squared test). (**E**) Optogenetic manipulation of the AI-PrL pathway in both stressed and unstressed dams did not affect PrL activity patterns during interaction with familiar mice (Chi-squared test). *ns* = non-significant ($p > 0.05$).

Supplementary Fig. 5

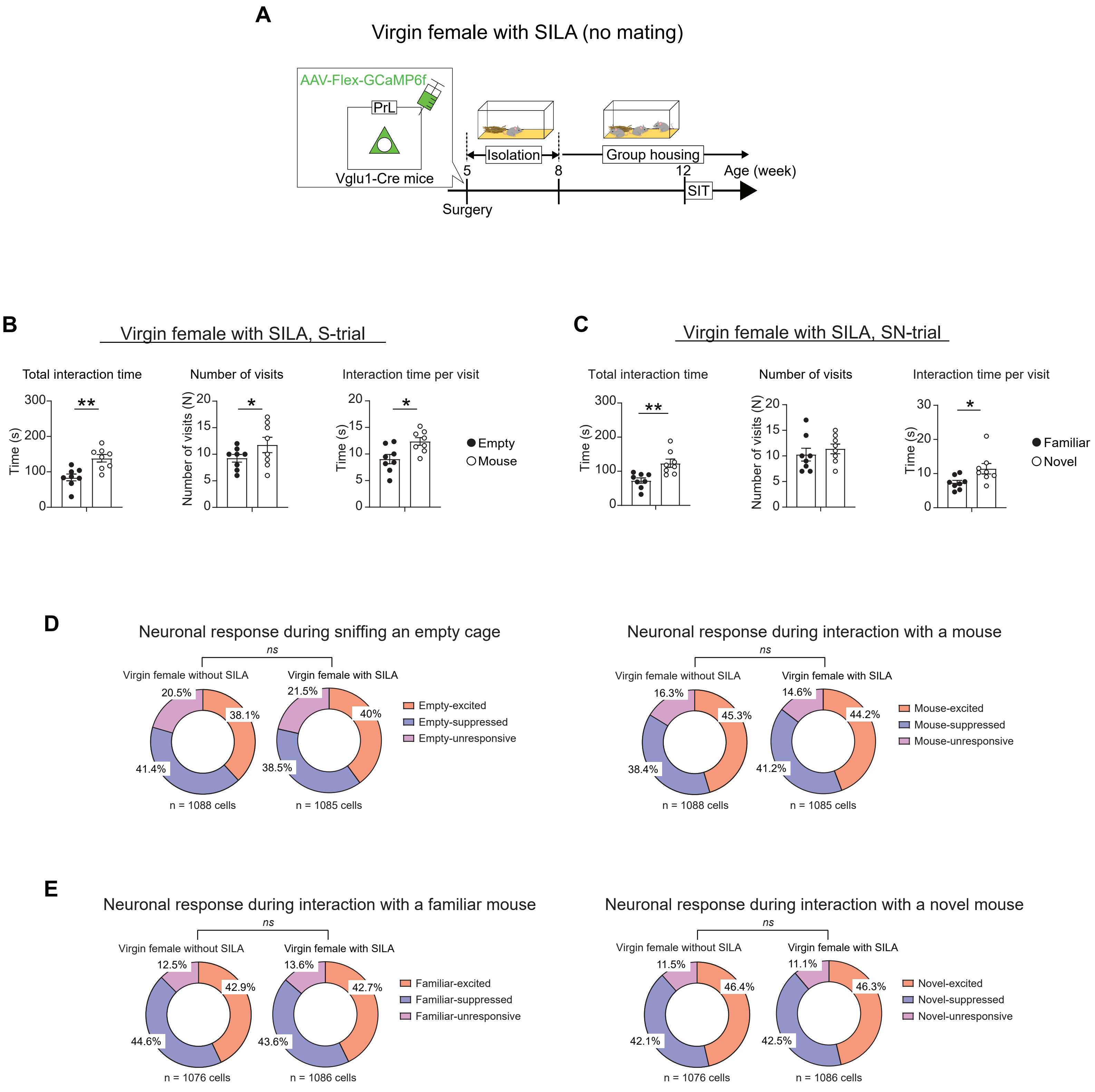

**Supplementary Fig. 5. SILA alone did not affect PrL activity patterns during SIT.** (**A**) Scheme of experimental timeline and strategy for *in vivo* microendoscopic calcium imaging in PrL. (**B** and **C**) In virgin females, SILA alone did not cause behavioral changes both in S- and SN-trials, in which they showed strong sociability and social novelty preference (Paired Student's $t$ test, Cohen's D > 0.8 for all significant $p$ values, and Wilcoxon signed rank test, $r$ > 0.8). (**D** and **E**) In virgin females, SILA alone did not affect PrL activity patterns during S- and SN-trials (Chi-squared test). *$p < 0.05$, **$p < 0.01$. *ns* = non-significant ($p > 0.05$). All data are represented as mean ± SEM.

Supplementary Fig. 6

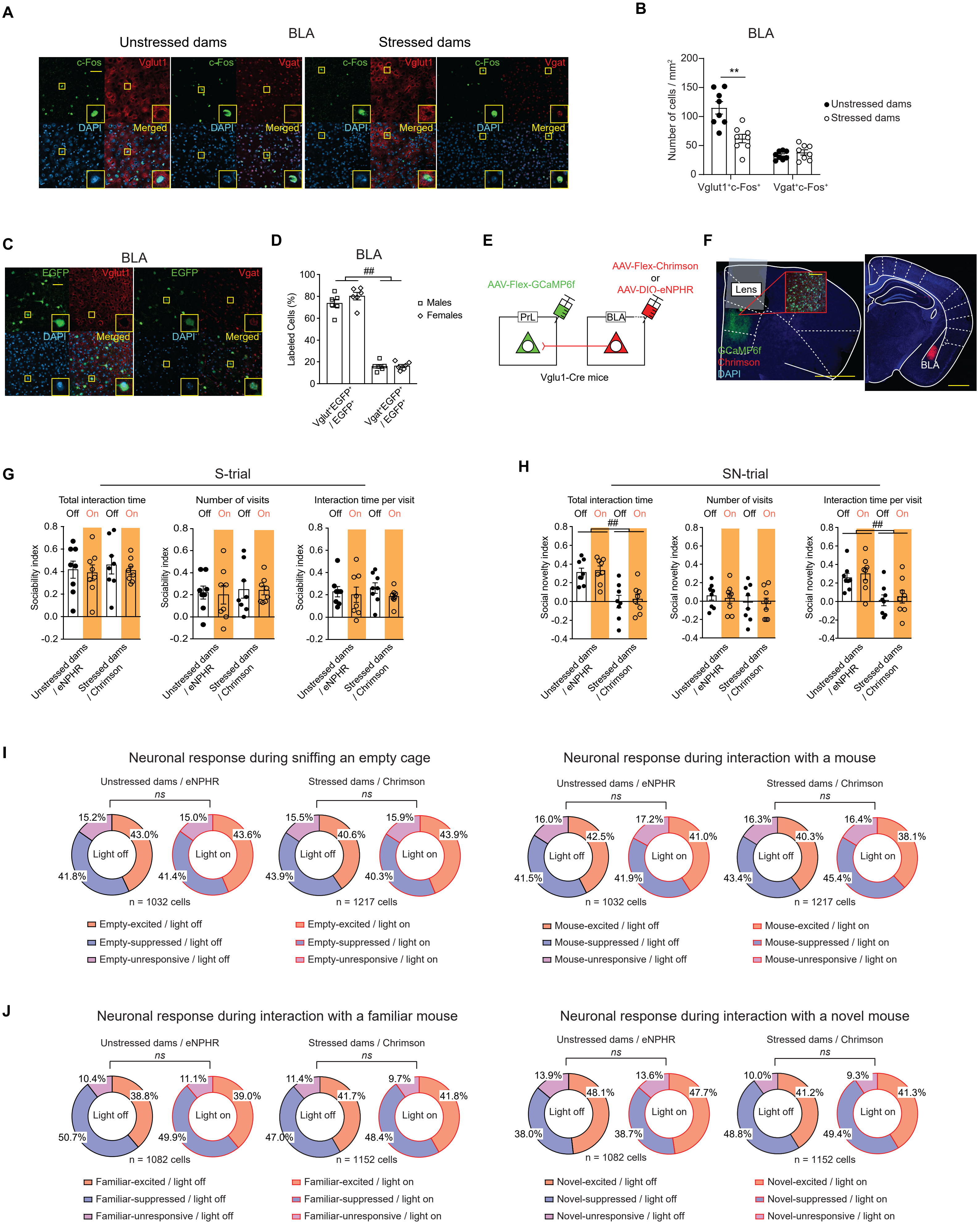

**Supplementary Fig. 6. BLA-PrL pathway did not affect PrL activity patterns and social behavior in the postpartum period.** (**A**) Representative images of c-Fos$^+$, Vglut1$^+$ or Vgat$^+$, DAPI, and colocalized cells in BLA of unstressed or stressed dams. Scale bar, 50 μm. (**B**) Stressed dams showed decreased number of Vglut1$^+$c-Fos$^+$ cells, but not Vgat$^+$c-Fos$^+$, in BLA after SN-trials (Student's *t* test, Welch *t* test). (**C**) Representative images of EGFP$^+$, Vglut1$^+$ or Vgat$^+$, DAPI, and colocalized cells in BLA from mice injected with AAVretro-hSyn-EGFP into PrL. Scale bar, 50 μm. (**D**) BLA-PrL pathway mainly consisted of Vglut1$^+$ neurons, but not Vgat$^+$ neurons (two-way ANOVA, main effect partial $\eta^2 = 0.98$). (**E**) Strategy of *in vivo* microendoscopic calcium imaging of PrL through optogenetic manipulation of the BLA-PrL pathway. (**F**) Representative images showing expressions of Chrimson in the BLA-PrL pathway and GCaMP6f underneath a GRIN lens. Scale bars, 1 mm and 100 μm. (**G** and **H**) Optogenetic manipulation of the BLA-PrL pathway did not affect social behavior in S- and SN-trials (two-way mixed ANOVA, Wilcoxon signed-rank test). (**I** and **J**) Optogenetic manipulation of the BLA-PrL pathway did not affect PrL activity patterns during S- and SN-trials (Chi-squared test). All data are represented as mean ± SEM. For ANOVAs, * indicates statistical significance for *post hoc* Bonferroni comparisons, and $^{\#}$ indicates statistical significance for the main effect. ** = $p < 0.01$, $^{\#\#} = p < 0.01$. *ns* = non-significant ($p > 0.05$).

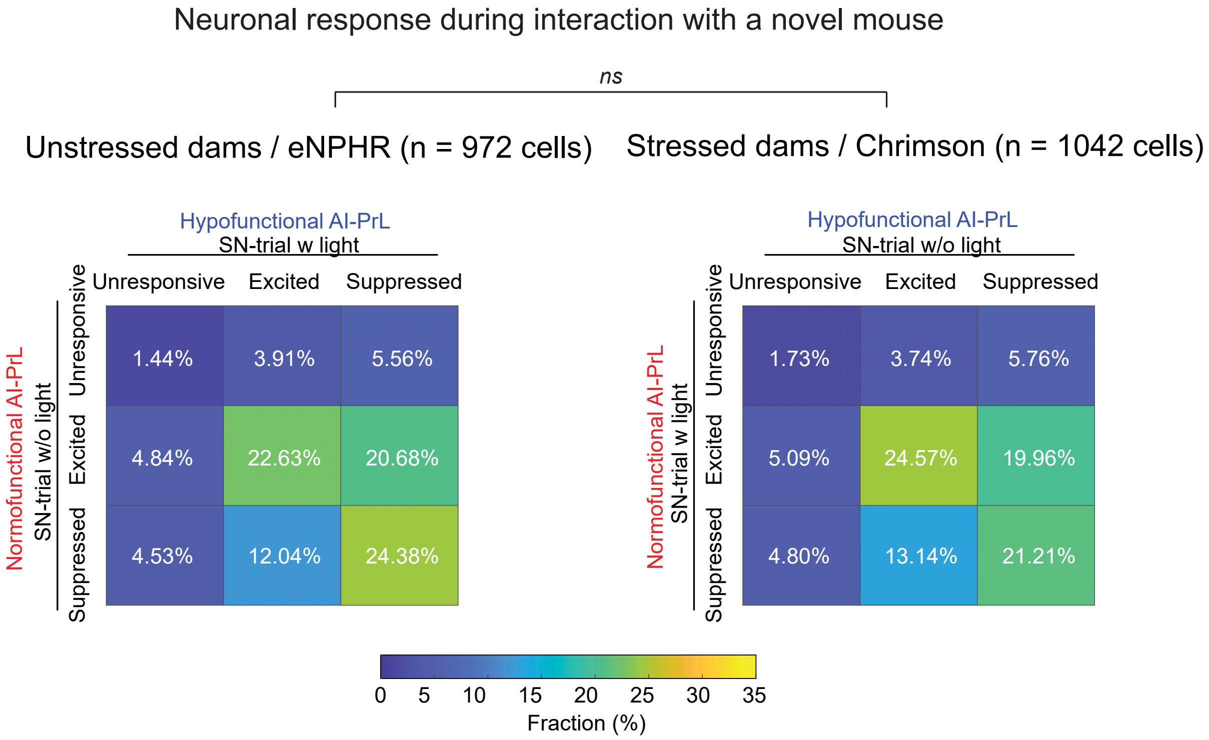
Neuronal response during interaction with a novel mouse
ns
Unstressed dams / eNPHR (n = 972 cells)
Stressed dams / Chrimson (n = 1042 cells)
Hypofunctional AI-PrL
SN-trial w light
Unresponsive
Excited
Suppressed
Normofunctional AI-PrL
SN-trial w/o light
1.44%
3.91%
5.56%
4.84%
22.63%
20.68%
4.53%
12.04%
24.38%
Hypofunctional AI-PrL
SN-trial w/o light
Normofunctional AI-PrL
SN-trial w light
1.73%
3.74%
5.76%
5.09%
24.57%
19.96%
4.80%
13.14%
21.21%
0 5 10 15 20 25 30 35
Fraction (%)

**Supplementary Fig. 7. Optogenetic activation of the AI-PrL pathway in stressed dams and optogenetic inhibition of AI-PrL pathway in unstressed dams showed opposite effects on patterns of PrL activity changes between SN-trials with and without light stimulation.** There were no significant differences between patterns of PrL activity changes in stressed dams from SN-trials with optogenetic activation to SN-trials without optogenetic activation, and those in unstressed dams from SN-trials without optogenetic inhibition to SN-trials with optogenetic inhibition (Chi-squared test). *ns* = non-significant ($p > 0.05$).

Supplementary Fig. 8

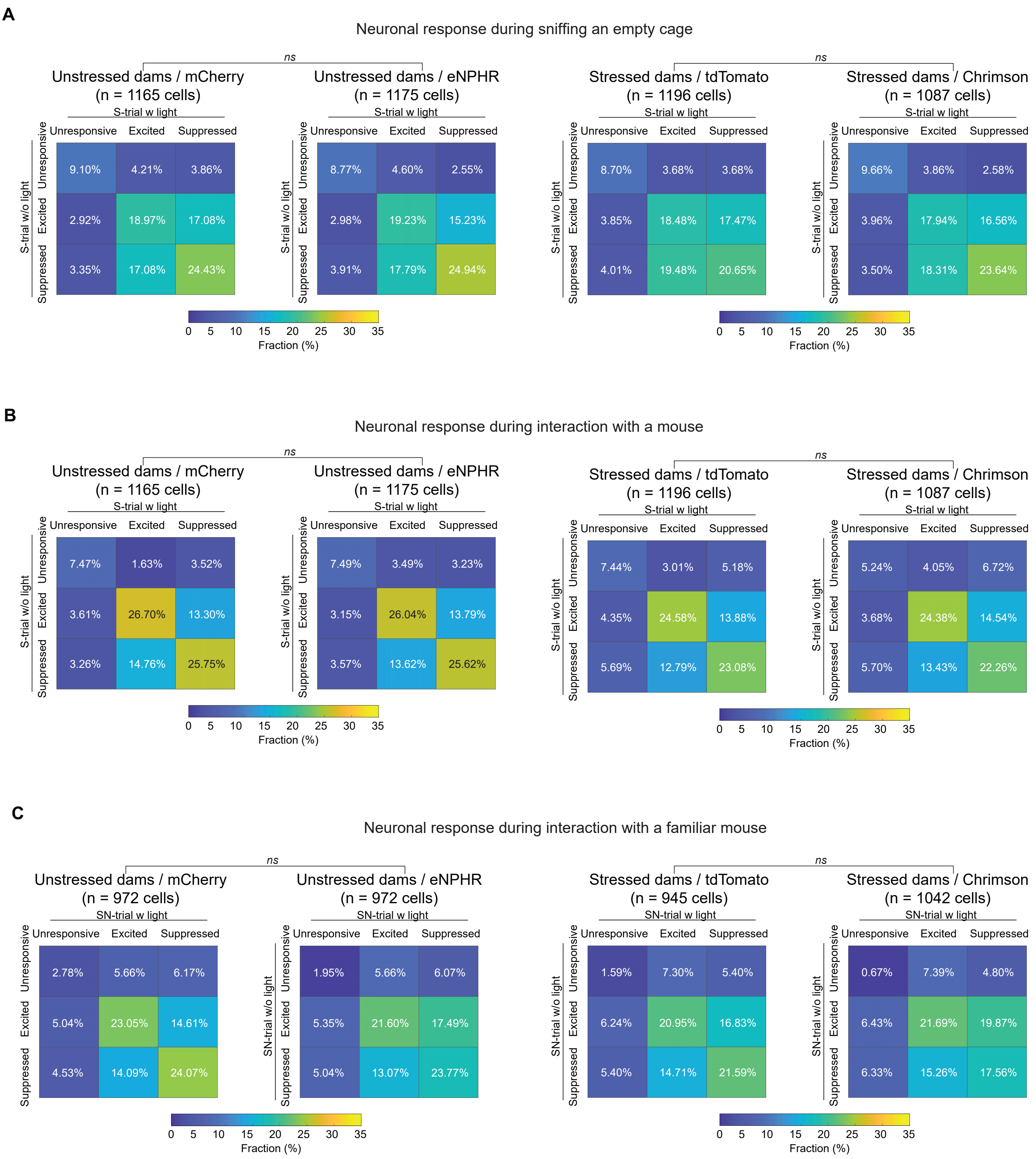

**Supplementary Fig. 8. AI-PrL pathway did not affect patterns of PrL activity changes during S-trials and interaction with a familiar mouse in SN-trials.** (**A** and **B**) Optogenetic manipulation of the AI-PrL pathway did not affect patterns of PrL activity changes from S-trials without light stimulation to S-trials with light stimulation (Chi-squared test). (**C**) Optogenetic manipulation of the AI-PrL pathway did not affect patterns of PrL activity changes during interaction with a familiar mouse from SN-trials without light stimulation to SN-trials with light stimulation (Chi-squared test). *ns* = non-significant ($p > 0.05$).

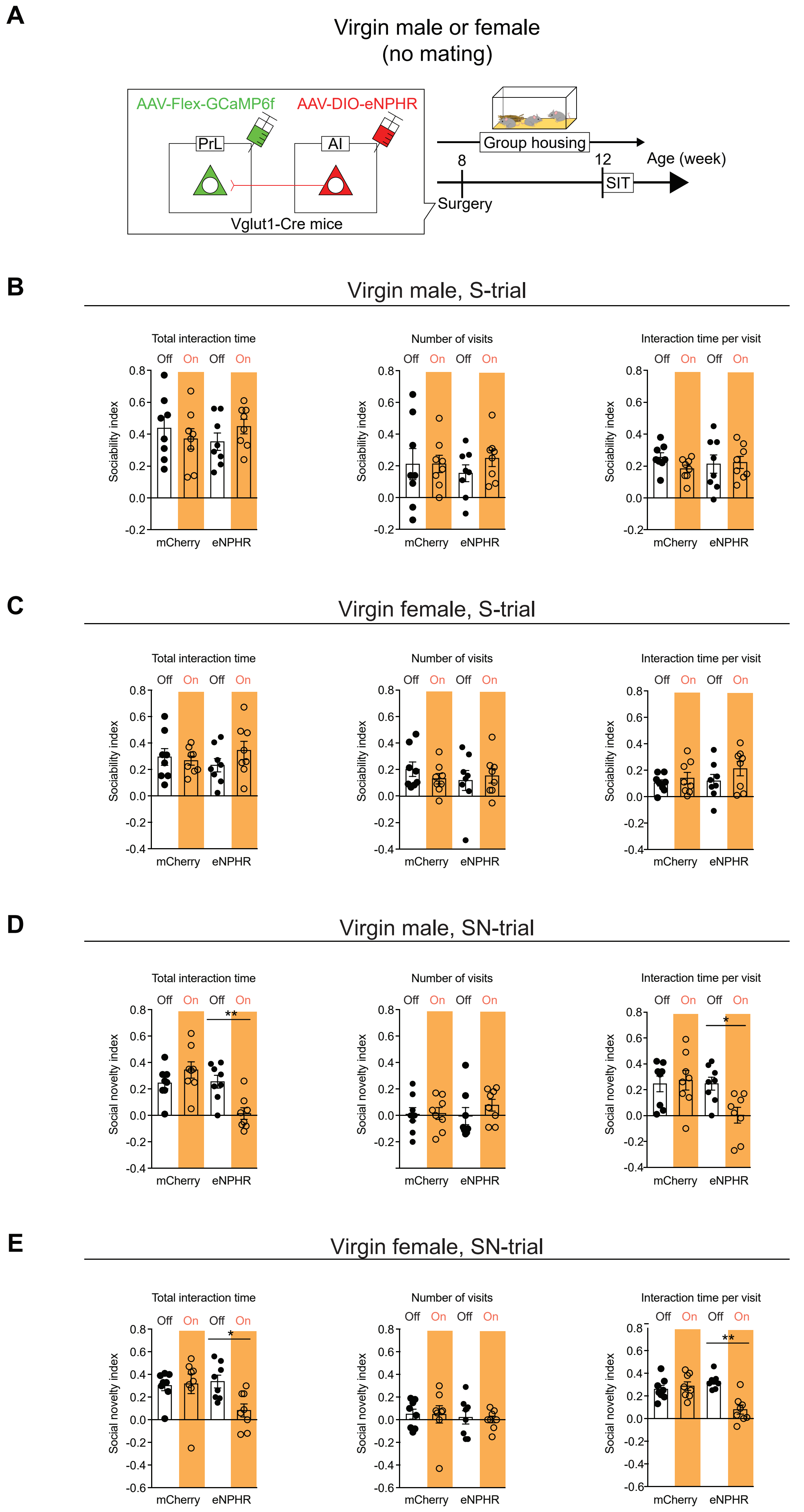
A
Virgin male or female
(no mating)
AAV-Flex-GCaMP6f
AAV-DIO-eNPHR
PrL
AI
Vglut1-Cre mice
Group housing
8
12
Age (week)
Surgery
SIT
B
Virgin male, S-trial
C
Virgin female, S-trial
D
Virgin male, SN-trial
E
Virgin female, SN-trial
Total interaction time
Number of visits
Interaction time per visit
Off On Off On
Sociability index
Social novelty index
mCherry
eNPHR
*
**

**Supplementary Fig. 9. Hypofunction of the AI-PrL pathway in virgin mice led to behavioral changes in social novelty recognition, but not sociability.** (**A**) Experimental timeline of SIT in virgin male and female mice. (**B** and **C**) Optogenetic inhibition of the AI-PrL pathway in virgin male and female mice did not affect sociability (two-way mixed ANOVA, Related samples sign test). (**D** and **E**) Optogenetic inhibition of the AI-PrL pathway in virgin male and female mice decreased total interaction time and interaction time per visit with novel mouse, but not the number of visits (two-way mixed ANOVA, Wilcoxon signed-rank test). All data are represented as mean ± SEM. For ANOVAs, * indicates statistical significance for *post hoc* Bonferroni comparisons. * = $p < 0.05$, ** = $p < 0.01$.

Supplementary Fig. 10

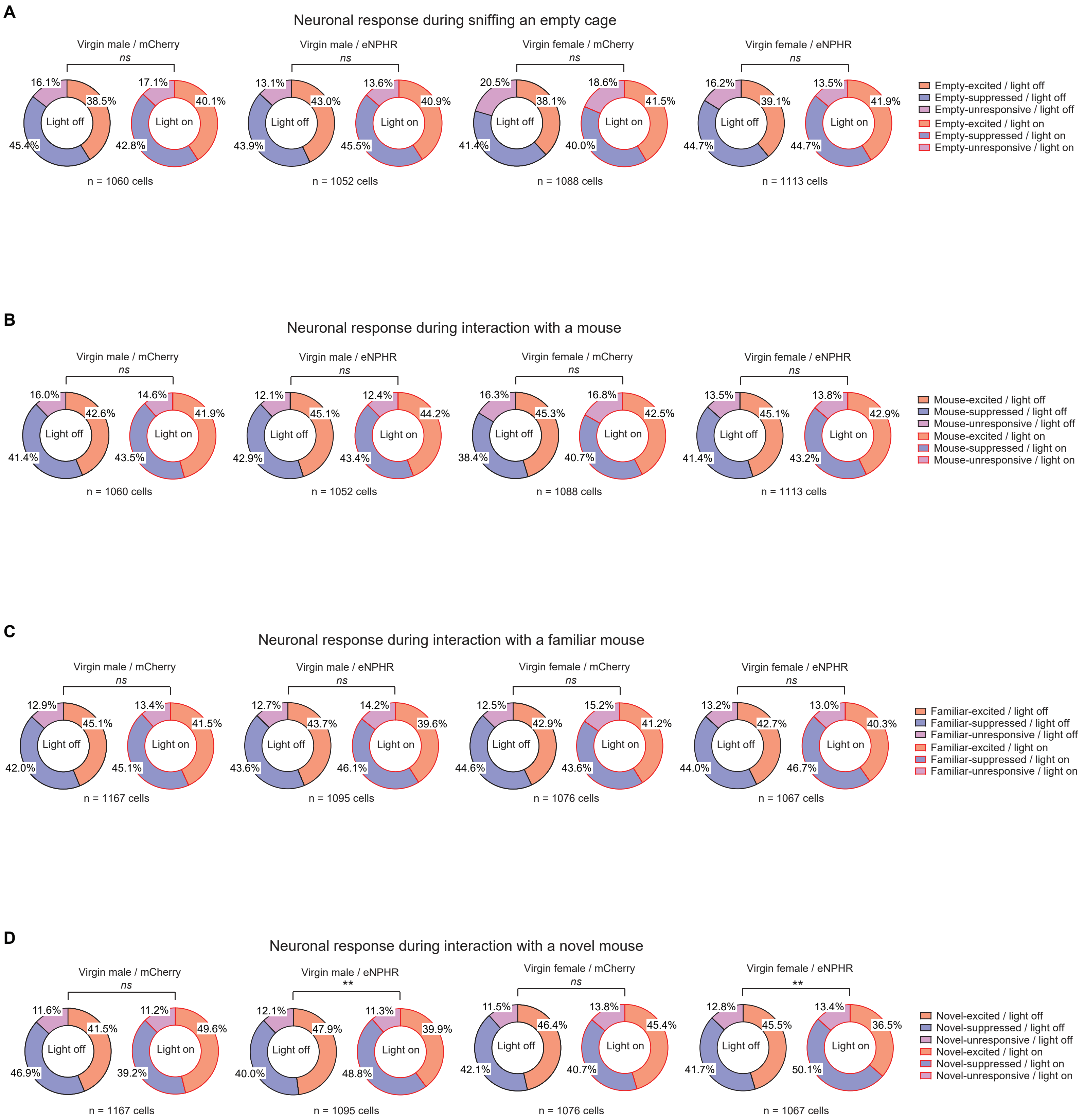

**Supplementary Fig. 10. Effects of optogenetic inhibition of the AI-PrL pathway on PrL activity in virgin mice were similar to those in unstressed dams.** (**A** and **B**) Optogenetic inhibition of the AI-PrL pathway in virgin mice did not affect PrL activity patterns during S-trials (Chi-squared test). (**C**) Optogenetic inhibition of the AI-PrL in virgin mice did not affect PrL activity patterns during interaction with a familiar mouse (Chi-squared test). (**D**) Optogenetic inhibition of the AI-PrL pathway in virgin mice resulted in similar patterns of PrL activity during interaction with a novel mouse to those observed in unstressed dams (Chi-squared test). *ns* = non-significant ($p > 0.05$). ** = $p < 0.01$.

A

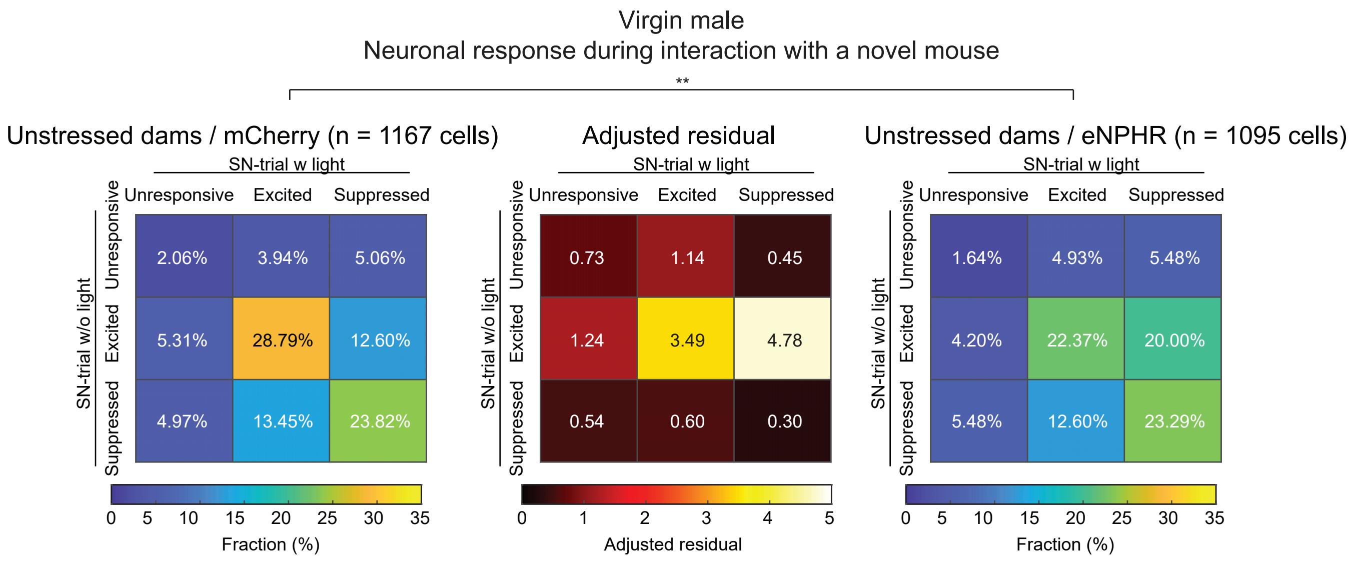

Virgin male
Neuronal response during interaction with a novel mouse
**
Unstressed dams / mCherry (n = 1167 cells)
Adjusted residual
Unstressed dams / eNPHR (n = 1095 cells)
SN-trial w light
SN-trial w/o light
Unresponsive
Excited
Suppressed
2.06%
3.94%
5.06%
5.31%
28.79%
12.60%
4.97%
13.45%
23.82%
0.73
1.14
0.45
1.24
3.49
4.78
0.54
0.60
0.30
1.64%
4.93%
5.48%
4.20%
22.37%
20.00%
5.48%
12.60%
23.29%
Fraction (%)
Adjusted residual

B

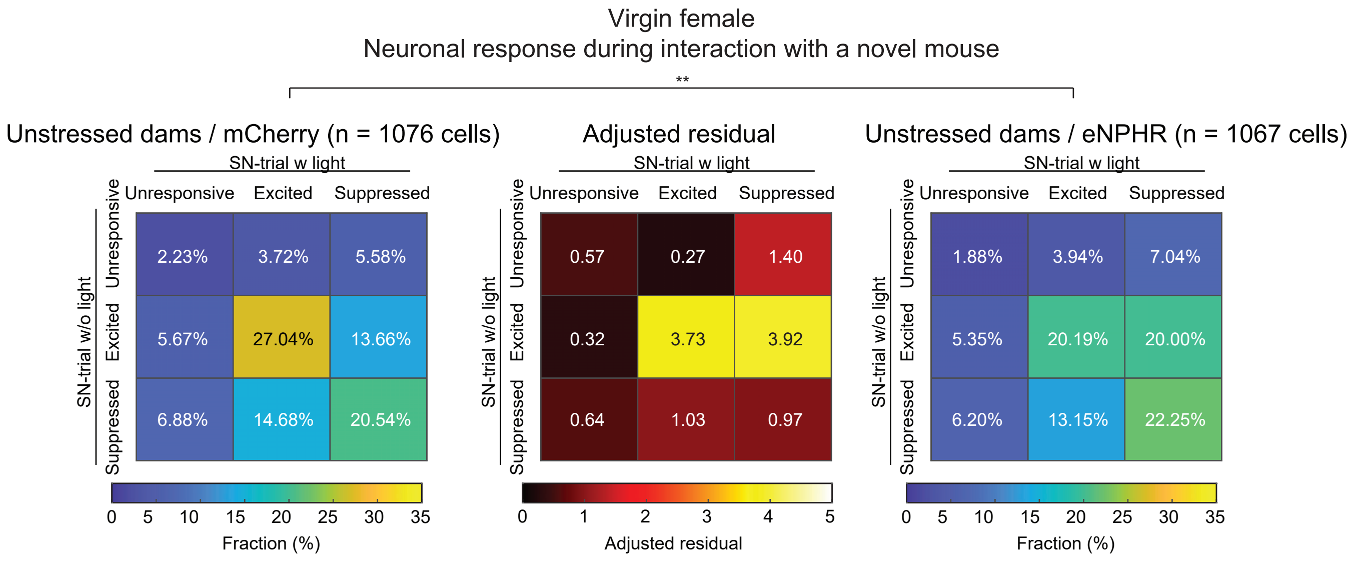

Virgin female
Neuronal response during interaction with a novel mouse
**
Unstressed dams / mCherry (n = 1076 cells)
Adjusted residual
Unstressed dams / eNPHR (n = 1067 cells)
SN-trial w light
SN-trial w/o light
Unresponsive
Excited
Suppressed
2.23%
3.72%
5.58%
5.67%
27.04%
13.66%
6.88%
14.68%
20.54%
0.57
0.27
1.40
0.32
3.73
3.92
0.64
1.03
0.97
1.88%
3.94%
7.04%
5.35%
20.19%
20.00%
6.20%
13.15%
22.25%
Fraction (%)
Adjusted residual

**Supplementary Fig. 11. Optogenetic inhibition of the AI-PrL pathway in virgin mice resulted in similar changes in PrL activity patterns to those observed in unstressed dams.** (**A** and **B**) Significant differences in the patterns of PrL activity changes from SN-trials without light stimulation to SN-trials with light stimulation were observed between virgin mice expressing control viruses and eNPHR (Chi-squared test). ** = $p < 0.01$.

Supplementary Fig. 12

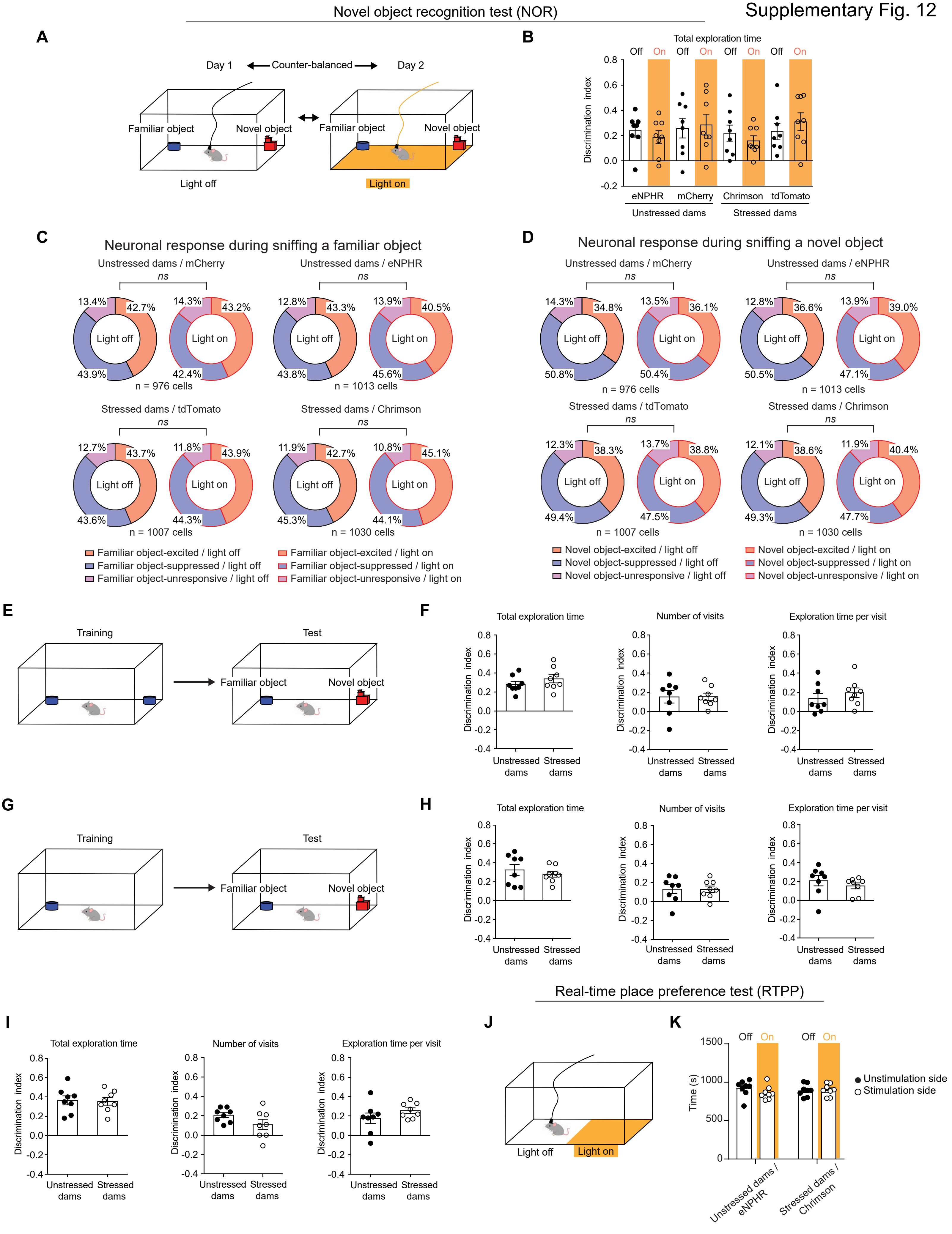

**Supplementary Fig. 12. AI-PrL pathway did not affect non-social object recognition and valence processing.** (**A**) Experimental timeline of NOR without and with optogenetic manipulation. (**B**) Optogenetic manipulation of the AI-PrL pathway did not affect novel object preference in both stressed and unstressed dams (three-way mixed ANOVA). (**C** and **D**) Optogenetic manipulation of the AI-PrL pathway did not affect PrL activity patterns during interaction with familiar or novel objects (Chi-squared test). **(E)** Scheme for NOR test. **(F)** NOR on postpartum day seven did not show significant differences between unstressed and stressed dams (Student's *t* test). **(G)** Scheme for modified NOR, analogous to SIT. **(H)** Modified NOR on postpartum day seven did not show significant differences between unstressed and stressed dams (Student's *t* test, Welch *t* test, Mann Whitney U test). **(I)** Modified NOR on postpartum day 13 did not show significant differences between unstressed and stressed dams (Student's *t* test). (**J**) Scheme for RTPP test. (**K**) Optogenetic manipulation of the AI-PrL pathway did not induce either reward or aversion for the side of the chamber (two-way mixed ANOVA). All data are represented as mean ± SEM. *ns* = non-significant ($p > 0.05$).

Supplementary Fig. 13

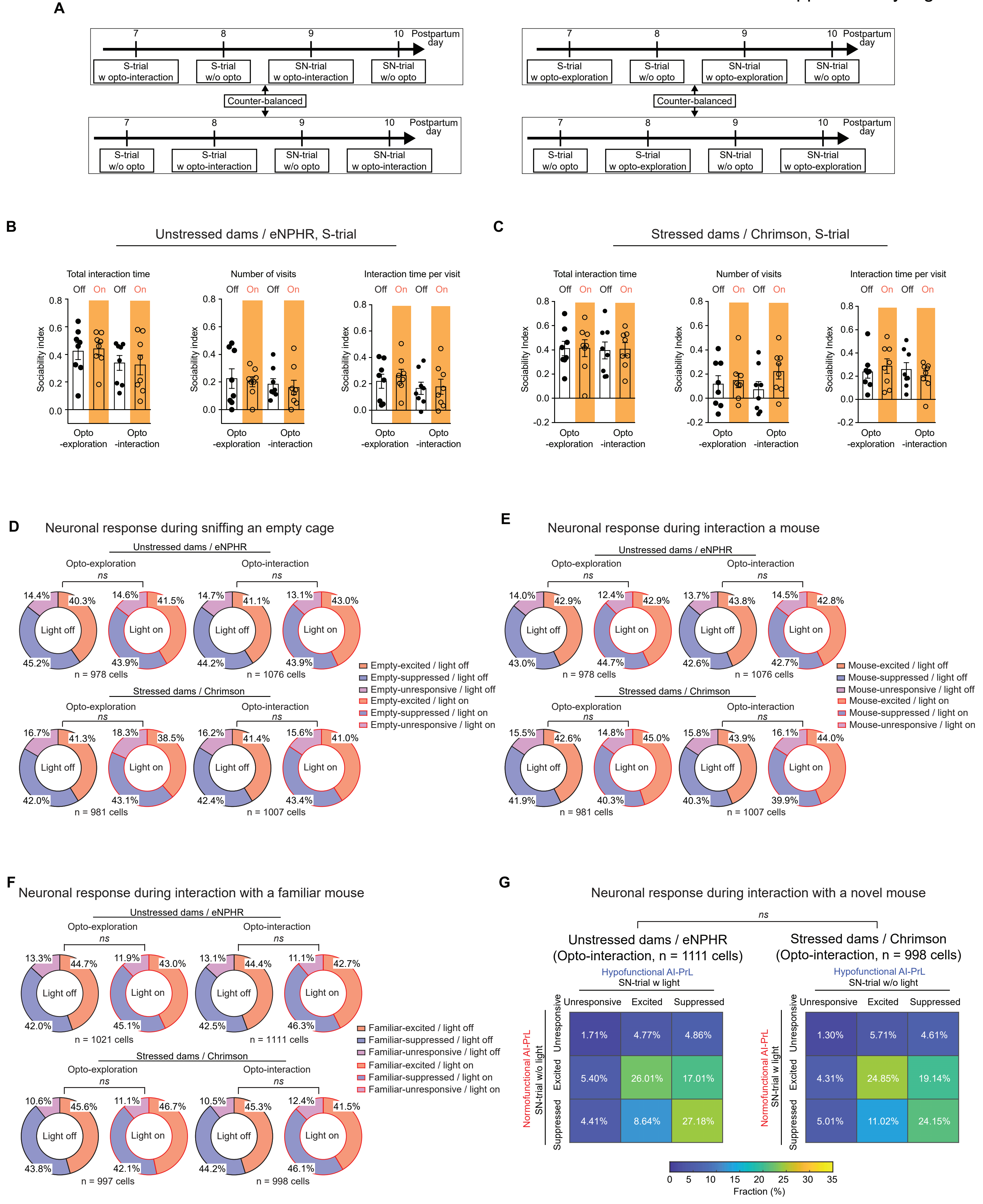

| Unstressed dams / eNPHR | Unresponsive | Excited | Suppressed |
|---|---|---|---|
| Unresponsive | 1.71% | 4.77% | 4.86% |
| Excited | 5.40% | 26.01% | 17.01% |
| Suppressed | 4.41% | 8.64% | 27.18% |

| Stressed dams / Chrimson | Unresponsive | Excited | Suppressed |
|---|---|---|---|
| Unresponsive | 1.30% | 5.71% | 4.61% |
| Excited | 4.31% | 24.85% | 19.14% |
| Suppressed | 5.01% | 11.02% | 24.15% |

**Supplementary Fig. 13. Behavioral closed-loop optogenetic manipulation of the AI-PrL pathway did not affect sociability and PrL activity during S-trials.** (**A**) Experimental timeline of SIT with and without behavioral closed-loop optogenetic manipulation. (**B** and **C**) Behavioral closed-loop optogenetic manipulation did not affect sociability in both unstressed and stressed dams (two-way mixed ANOVA, Wilcoxon signed-rank test). (**D** and **E**) Behavioral closed-loop optogenetic manipulation did not affect PrL activity patterns during S-trials (Chi-squared test). (**F**) Behavioral closed-loop optogenetic manipulation did not affect PrL activity patterns during interaction with a familiar mouse in SN-trials (Chi-squared test). (**G**) There were no significant differences between patterns of PrL activity changes in stressed dams from SN-trials with optogenetic activation during interaction to SN-trials without optogenetic activation, and those in unstressed dams from SN-trials without optogenetic inhibition to SN-trials with optogenetic inhibition during interaction (Chi-squared test). All data are represented as mean ± SEM. *ns* = non-significant ($p > 0.05$).

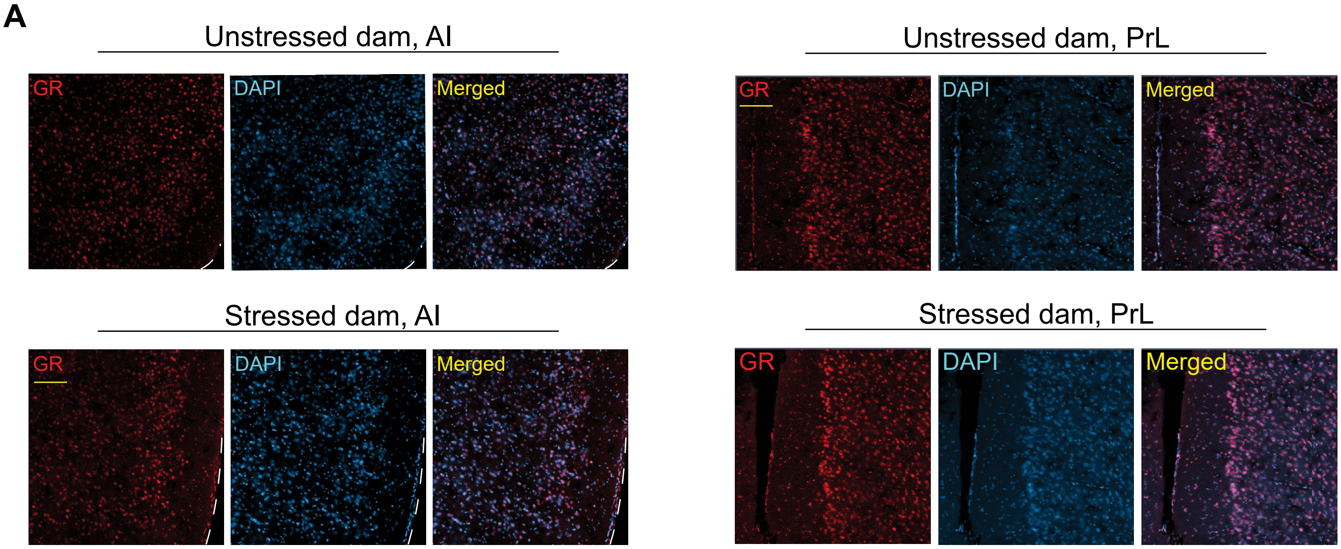
A
Unstressed dam, AI
GR
DAPI
Merged
Unstressed dam, PrL
GR
DAPI
Merged
Stressed dam, AI
GR
DAPI
Merged
Stressed dam, PrL
GR
DAPI
Merged

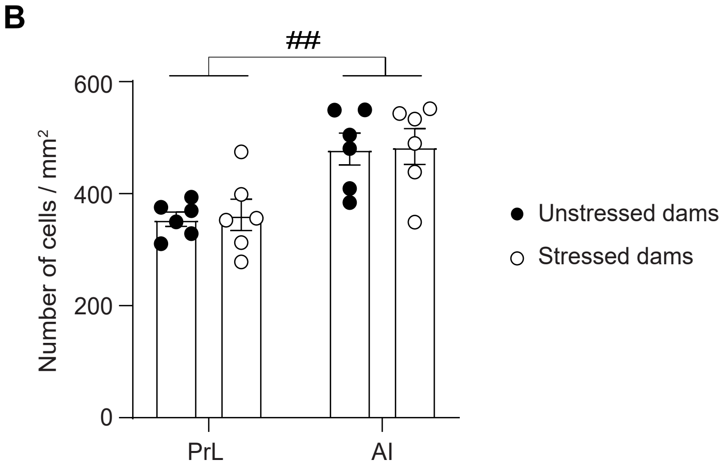
B
##
600
400
200
0
Number of cells / mm²
PrL
AI
Unstressed dams
Stressed dams

**Supplementary Fig. 14. Expression level of the glucocorticoid receptor (GR) was not affected by SILA. (A)** Representative images of $GR^+$ cells in PrL and AI of unstressed and stressed dams. **(B)** There were no significant differences in the expression level of GR both in PrL and AI between unstressed and stressed dams (two-way mixed ANOVA). All data are represented as mean ± SEM. $^{\#\#}$ = ANOVA main effect for brain region, $p < 0.01$.

Supplementary Fig. 15

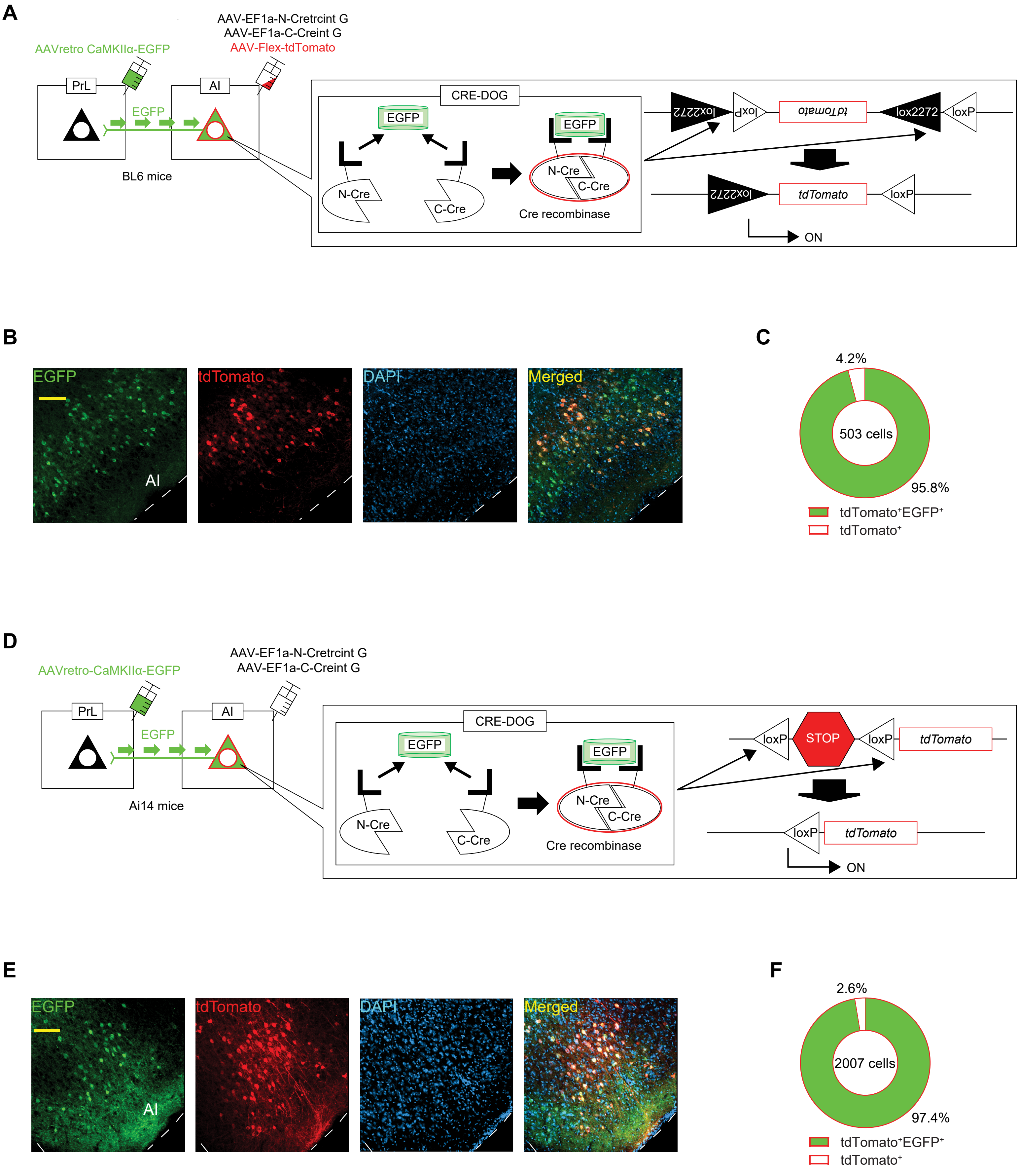

**Supplementary Fig. 15. Cre recombinase was successfully expressed in the AI-PrL pathway by the CRE-DOG method.** (**A**) Scheme of the CRE-DOG validation study with B6J mice injected with AAV-Flex-tdTomato. (**B**) Representative images of EGFP$^+$, tdTomato$^+$, DAPI, and colocalized cells in AI of B6J mice. Scale bar, 100 μm. (**C**) CRE-DOG method successfully expressed tdTomato in cells expressing EGFP in B6J mice. (**D**) Scheme of the CRE-DOG validation study with Ai14 mice. (**E**) Representative images of EGFP$^+$, tdTomato$^+$, DAPI, and colocalized cells in AI of Ai14 mice. (**F**) CRE-DOG method successfully expressed tdTomato in cells expressing EGFP in Ai14 mice. Scale bar, 100 μm.

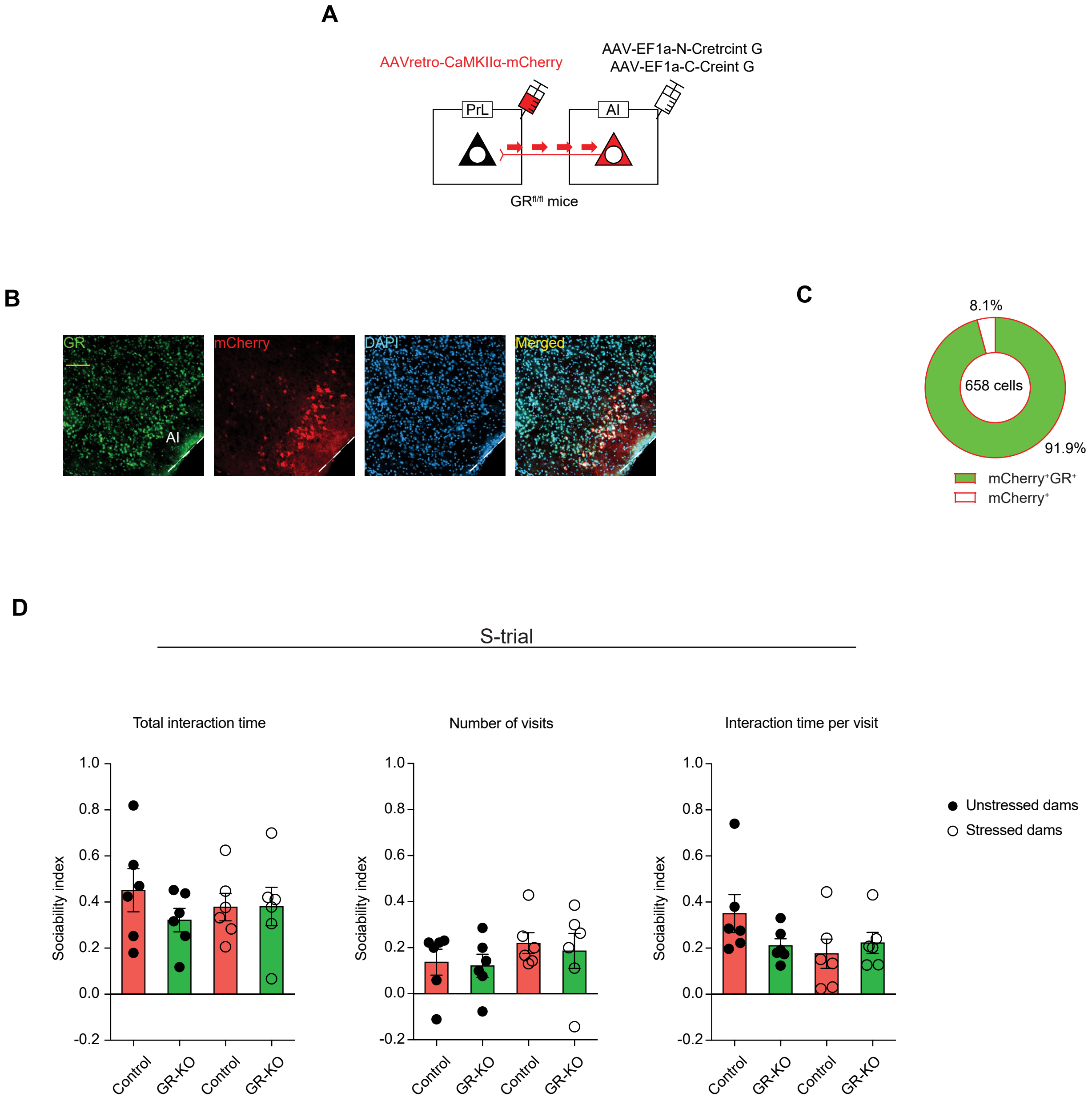
A
AAVretro-CaMKIIα-mCherry
AAV-EF1a-N-Cretrcint G
AAV-EF1a-C-Creint G
PrL
AI
GR fl/fl mice
B
GR
mCherry
DAPI
Merged
AI
C
8.1%
658 cells
91.9%
mCherry+GR+
mCherry+
D
S-trial
Total interaction time
Number of visits
Interaction time per visit
Sociability index
Control
GR-KO
Unstressed dams
Stressed dams

**Supplementary Fig. 16. Pathway-specific GR-KO did not affect the results of S-trials.** (**A**) Scheme of the virus strategy in $GR^{fl/fl}$ mice injected with AAV-retro-CaMKIIα-mCherry instead of AAV2-retro CaMKIIα-EGFP. (**B**) Representative images of $GR^{+}$, $mCherry^{+}$, DAPI, and colocalized cells in AI of $GR^{fl/fl}$ mice without CRE-DOG (no Cre expression in the AI-PrL pathway). (**C**) GR expression was not deleted in $GR^{fl/fl}$ mice without CRE-DOG. (**D**) No significant change was identified in sociability in AI-PrL pathway-specific GR-KO mice (two-way ANOVA). All data are represented as mean ± SEM.

**Supplementary Table 1. Abbreviations of brain areas.**

| Abbreviation | Brain area |
|---|---|
| MO | Medial orbital cortex |
| VO | Ventral orbital cortex |
| LO | Lateral orbital cortex |
| M1 | Primary motor cortex |
| M2 | Secondary motor cortex |
| Cg | Cingulate cortex |
| IL | Infralimbic cortex |
| AM | Anteromedial nucleus of thalamus |
| VM | Ventromedial nucleus of thalamus |
| RSA | Retrosplenial agran cortex |
| V2 | Secondary visual cortex |
| V1 | Primary visual cortex |
| S2 | Secondary somatosensory cortex |
| MD | Mediodorsal nucleus of thalamus |
| RE | Nucleus reuniens of thalamus |
| DEN | Dorsal endopiriform nucleus |
| BMA | Basomedial nucleus of amygdala |
| BLA | Basolateral nucleus of amygdala |
| ECT | Ectorhinal cortex |
| PRh | Perirhinal cortex |
| LEnt | Lateral entorhinal cortex |
| Pir | Piriform cortex |
| S | Subiculum |
| CA1 | CA1 region of hippocampus |
| DR | Dorsal raphe nucleus |
| VTA | Ventral tegmental area |

**Supplementary Table 2. Definitions of subsets of PrL neurons.**

| Behavioral test | Event used for ROC analysis | Name | Definition |
|---|---|---|---|
| S-trial of SIT | Sniffing an empty cage | Empty-excited / light off | Neurons which that are excited during sniffing an empty cage in S-trials without light stimulation |
| | | Empty-suppressed / light off | Neurons that are suppressed during sniffing an empty cage in S-trials without light stimulation |
| | | Empty-unresponsive / light off | Neurons that are not excited or suppressed during sniffing an empty cage in S-trials without light stimulation |
| | Interaction with a mouse | Mouse-excited / light off | Neurons that are excited during interaction with a mouse in S-trials without light stimulation |
| | | Mouse-suppressed / light off | Neurons that are suppressed during interaction with a mouse in S-trials without light stimulation |
| | | Mouse-unresponsive / light off | Neurons that are not excited or suppressed during interaction with a mouse in S-trials without light stimulation |
| | Sniffing an empty cage | Empty-excited / light on | Neurons that are excited during sniffing an empty cage in S-trials with light stimulation |
| | | Empty-suppressed / light on | Neurons that are suppressed during sniffing an empty cage in S-trials with light stimulation |
| | | Empty-unresponsive / light on | Neurons that are not excited or suppressed during sniffing an empty cage in S-trials with light stimulation |
| | Interaction with a mouse | Mouse-excited / light on | Neurons that are excited during interaction with a mouse in S-trials with light stimulation |
| | | Mouse-suppressed / light on | Neurons that are suppressed during interaction with a mouse in S-trials with light stimulation |
| | | Mouse-unresponsive / light on | Neurons that are not excited or suppressed during interaction with a mouse in S-trials with light stimulation |
| SN-trial of SIT | Interaction with a familiar mouse | Familiar-excited / light off | Neurons that are excited during interaction with a familiar mouse in SN-trials without light stimulation |
| | | Familiar-suppressed / light off | Neurons that are suppressed during interaction with a familiar mouse in SN-trials without light stimulation |
| | | Familiar-unresponsive / light off | Neurons that are not excited or suppressed during interaction with a familiar mouse in SN-trials without light stimulation |

| | | | |
|---|---|---|---|
| | Interaction with a novel mouse | Novel-excited / light off | Neurons that are excited during interaction with a novel mouse in SN-trials without light stimulation |
| | | Novel-suppressed / light off | Neurons that are suppressed during interaction with a novel mouse in SN-trials without light stimulation |
| | | Novel-unresponsive / light off | Neurons that are not excited or suppressed during interaction with a novel mouse in SN-trials without light stimulation |
| | Interaction with a familiar mouse | Familiar-excited / light on | Neurons that are excited during interaction with a familiar mouse in SN-trials with light stimulation |
| | | Familiar-suppressed / light on | Neurons that are suppressed during interaction with a familiar mouse in SN-trials with light stimulation |
| | | Familiar-unresponsive / light on | Neurons that are not excited or suppressed during interaction with a familiar mouse in SN-trials with light stimulation |
| | Interaction with a novel mouse | Novel-excited / light on | Neurons that are excited during interaction with a novel mouse in SN-trials with light stimulation |
| | | Novel-suppressed / light on | Neurons that are suppressed during interaction with a novel mouse in SN-trials with light stimulation |
| | | Novel-unresponsive / light on | Neurons that are not excited or suppressed during interaction with a novel mouse in SN-trials with light stimulation |
| NOR | Sniffing a familiar object | Familiar object-excited / light off | Neurons that are excited during sniffing a familiar object in NOR without light stimulation |
| | | Familiar object-suppressed / light off | Neurons that are suppressed during sniffing a familiar object in NOR without light stimulation |
| | | Familiar object-unresponsive / light off | Neurons that are not excited or suppressed during sniffing a familiar object in NOR without light stimulation |
| | Sniffing a novel object | Novel object-excited / light off | Neurons that are excited during sniffing a novel object in NOR without light stimulation |
| | | Novel object-suppressed / light off | Neurons that are suppressed during sniffing a novel object in NOR without light stimulation |
| | | Novel object-unresponsive / light off | Neurons that are not excited or suppressed during sniffing a novel object in NOR without light stimulation |

| | | | |
|---|---|---|---|
| | Sniffing a familiar object | Familiar object-excited / light on | Neurons that are excited during sniffing a familiar object in NOR with light stimulation |
| | | Familiar object-suppressed / light on | Neurons that are suppressed during sniffing a familiar object in NOR with light stimulation |
| | | Familiar object-unresponsive / light on | Neurons that are not excited or suppressed during sniffing a familiar object in NOR with light stimulation |
| | Sniffing a novel object | Novel object-excited / light on | Neurons that are excited during sniffing a novel object in NOR with light stimulation |
| | | Novel object-suppressed / light on | Neurons that are suppressed during sniffing a novel object in NOR with light stimulation |
| | | Novel object-unresponsive / light on | Neurons that are not excited or suppressed during sniffing a novel object in NOR with light stimulation |
| Note that neurons with a suppressed response as defined with this method do not necessarily exhibit an immediate decrease of activity during the corresponding behavior, but rather display an overall negative correlation with the corresponding behavior. | | | |

**Supplementary Table 3. Detailed information on statistical analyses and sample sizes.**

Statistical tests performed for each data set are shown.

In the ANOVA designs, (W) indicates a within-subjects factor, and (B) indicates a between-subjects factor. "Opsin" is used as a factor in several designs, meaning that groups have been treated (+) with and (-) without opsin expression. In the optogenetics experiments, "OFF" and "ON" are used to indicate the light conditions.

| Figure | Shapiro-Wilk test | Levene's test | Two-Way mixed ANOVA |
|---|---|---|---|
| **1C** | Total interaction time | | |
| | Unstressed Empty $p =$ 0.595 | Empty $p =$ 0.330 | Interaction F(1,14) = 0.214 , $p =$ 0.651 , partial $\eta^2 =$ 0.015 |
| | Stressed Empty $p =$ 0.178 | Mouse $p =$ 0.171 | Sociability (W) F(1,14) = 64.333 , $p =$ <0.001 , partial $\eta^2 =$ 0.821 |
| | Unstressed Mouse $p =$ 0.104 | | Stress (B) F(1,14) = 0.014 , $p =$ 0.907 , partial $\eta^2 =$ 0.001 |
| | Stressed Mouse $p =$ 0.329 | | |
| | | Box's M test | post hoc (Bonferroni) |
| | | $p =$ 0.335 | Unstressed Empty x Mouse $p =$ <0.001 |
| | | | Stressed Empty x Mouse $p =$ <0.001 |
| | Number of visits | | **Two-Way mixed ANOVA** |
| | Unstressed Empty $p =$ 0.085 | Empty $p =$ 0.082 | Interaction F(1,14) = 0.009 , $p =$ 0.924 , partial $\eta^2 =$ <0.001 |
| | Stressed Empty $p =$ 0.424 | Mouse $p =$ 0.230 | Sociability (W) F(1,14) = 15.668 , $p =$ 0.001 , partial $\eta^2 =$ 0.528 |
| | Unstressed Mouse $p =$ 0.067 | | Stress (B) F(1,14) = 0.002 , $p =$ 0.966 , partial $\eta^2 =$ <0.001 |
| | Stressed Mouse $p =$ 0.535 | | |
| | | Box's M test | post hoc (Bonferroni) |
| | | $p =$ 0.632 | Unstressed Empty x Mouse $p =$ 0.016 |
| | | | Stressed Empty x Mouse $p =$ 0.012 |
| | Interaction time per visit | | **Two-Way mixed ANOVA** |
| | Unstressed Empty $p =$ 0.386 | Empty $p =$ 0.580 | Interaction F(1,14) = 0.081 , $p =$ 0.780 , partial $\eta^2 =$ 0.006 |
| | Stressed Empty $p =$ 0.111 | Mouse $p =$ 0.265 | Sociability (W) F(1,14) = 12.206 , $p =$ 0.004 , partial $\eta^2 =$ 0.466 |
| | Unstressed Mouse $p =$ 0.111 | | Stress (B) F(1,14) = 0.478 , $p =$ 0.501 , partial $\eta^2 =$ 0.033 |
| | Stressed Mouse $p =$ 0.610 | | |
| | **Sample Size** | Box's M test | post hoc (Bonferroni) |
| | Unstressed $n =$ 8 | $p =$ 0.038 | Unstressed Empty x Mouse $p =$ 0.018 |
| | Stressed $n =$ 8 | | Stressed Empty x Mouse $p =$ 0.040 |
| **Figure** | **Shapiro-Wilk test** | **Levene's test** | **Student t-Test Independent samples, 2-tailed** |
| **1C (Socialbility index)** | Total interaction time | | |
| | Unstressed $p =$ 0.991 | $p =$ 0.302 | $t$ (14) = -0.331 , $p =$ 0.746 |
| | Stressed $p =$ 0.600 | | Cohen's d = -0.165 |
| | Number of visits | | **Student t-Test Independent samples, 2-tailed** |
| | Unstressed $p =$ 0.293 | $p =$ 0.493 | $t$ (14) = -0.127 , $p =$ 0.900 |
| | Stressed $p =$ 0.356 | | Cohen's d = -0.064 |
| | Interaction time per visit | | **Student t-Test Independent samples, 2-tailed** |
| | Unstressed $p =$ 0.092 | $p =$ 0.537 | $t$ (14) = 0.215 , $p =$ 0.833 |
| | Stressed $p =$ 0.268 | | Cohen's d = 0.108 |
| | **Sample Size** | | |
| | Unstressed $n =$ 8 | | |
| | Stressed $n =$ 8 | | |
| **Figure** | **Shapiro-Wilk test** | **Levene's test** | **Two-Way mixed ANOVA** |
| **1E** | Total interaction time | | |
| | Unstressed Familiar $p =$ 0.245 | Familiar $p =$ 0.316 | Interaction F(1,14) = 90.513 , $p =$ <0.001 , partial $\eta^2 =$ 0.866 |
| | Stressed Familiar $p =$ 0.058 | Novel $p =$ 0.934 | Social novelty (W) F(1,14) = 71.536 , $p =$ <0.001 , partial $\eta^2 =$ 0.836 |
| | Unstressed Novel $p =$ 0.517 | | Stress (B) F(1,14) = 0.567 , $p =$ 0.464 , partial $\eta^2 =$ 0.039 |
| | Stressed Novel $p =$ 0.901 | | |
| | | Box's M test | post hoc (Bonferroni) |
| | | $p =$ 0.048 | Unstressed Familiar x Novel $p =$ <0.001 |
| | | | Stressed Familiar x Novel $p =$ 0.468 |
| | Number of visits | | **Two-Way mixed ANOVA** |
| | Unstressed Familiar $p =$ 0.117 | Familiar $p =$ 0.405 | Interaction F(1,14) = 0.003 , $p =$ 0.960 , partial $\eta^2 =$ <0.001 |
| | Stressed Familiar $p =$ 0.861 | Novel $p =$ 0.211 | Social novelty (W) F(1,14) = 0.064 , $p =$ 0.803 , partial $\eta^2 =$ 0.005 |
| | Unstressed Novel $p =$ 0.164 | | Stress (B) F(1,14) = 0.344 , $p =$ 0.567 , partial $\eta^2 =$ 0.024 |
| | Stressed Novel $p =$ 0.551 | | |
| | | Box's M test | post hoc (Bonferroni) |
| | | $p =$ 0.496 | Unstressed Familiar x Novel $p =$ 0.833 |
| | | | Stressed Familiar x Novel $p =$ 0.888 |

| | | | |
|---|---|---|---|
| | Interaction time per visit<br>Unstressed Familiar *p* = 0.098<br>Stressed Familiar *p* = 0.387<br>Unstressed Novel *p* = 0.050<br>Stressed Novel *p* = 0.062<br>**Sample Size**<br>Unstressed *n* = 8<br>Stressed *n* = 8 | Familiar *p* = 0.141<br>Novel *p* = 0.616<br><br><br><br>Box's M test<br>*p* = 0.249 | **Two-Way mixed ANOVA**<br>Interaction F(1,14) = 6.425 , *p* = 0.024 , partial $\eta^2$ = 0.315<br>Social novelty (W) F(1,14) = 7.481 , *p* = 0.016 , partial $\eta^2$ = 0.348<br>Stress (B) F(1,14) = 0.001 , *p* = 0.979 , partial $\eta^2$ = <0.001<br><br>post hoc (Bonferroni)<br>Unstressed Familiar x Novel *p* = 0.002<br>Stressed Familiar x Novel *p* = 0.889 |
| **Figure<br>1E (social novelty index)** | **Shapiro-Wilk test**<br>Total interaction time<br>Unstressed *p* = 0.771<br>Stressed *p* = 0.292<br><br>Number of visits<br>Unstressed *p* = 0.813<br>Stressed *p* = 0.745<br><br>Interaction time per visit<br>Unstressed *p* = 0.767<br>Stressed *p* = 0.659<br>**Sample Size**<br>Unstressed *n* = 8<br>Stressed *n* = 8 | **Levene's test**<br><br>*p* = 0.094<br><br><br><br>*p* = 0.797<br><br><br><br>*p* = 0.248 | **Student t-Test Independent samples, 2-tailed**<br>*t* (14) = 9.531 , *p* = <0.001<br>Cohen's d = 4.766<br><br>**Student t-Test Independent samples, 2-tailed**<br>*t* (14) = 0.1373 , *p* = 0.893<br>Cohen's d = 0.0687<br><br>**Student t-Test Independent samples, 2-tailed**<br>*t* (14) = 2.9561 , *p* = 0.010<br>Cohen's d = 1.478 |
| **Figure<br>1G** | **Shapiro-Wilk test**<br>Unstressed Vglut *p* = 0.501<br>Stressed Vglut *p* = 0.343<br>Unstressed Vgat *p* = 0.192<br>Stressed Vgat *p* = 0.145<br>**Sample Size**<br>Unstressed *n* = 8<br>Stressed *n* = 8 | **Levene's test**<br>Vglut *p* = 0.970<br>Vgat *p* = 0.823<br><br>Box's M test<br>*p* = 0.618 | **Two-Way mixed ANOVA**<br>Interaction F(1,14) = 36.148 , *p* = <0.001 , partial $\eta^2$ = 0.721<br>Marker (W) F(1,14) = 150.36 , *p* = <0.001 , partial $\eta^2$ = 0.915<br>Stress (B) F(1,14) = 34.523 , *p* = <0.001 , partial $\eta^2$ = 0.711<br><br>post hoc (Bonferroni)<br>Vglut Unstressed x Stressed *p* = <0.001<br>Vgat Unstressed x Stressed *p* = 0.699 |
| **Figure<br>1I** | **Shapiro-Wilk test**<br>Unstressed Vglut *p* = 0.699<br>Stressed Vglut *p* = 0.127<br>Unstressed Vgat *p* = 0.607<br>Stressed Vgat *p* = 0.222<br>**Sample Size**<br>Unstressed *n* = 8<br>Stressed *n* = 8 | **Levene's test**<br>Vglut *p* = 0.823<br>Vgat *p* = 0.266<br><br>Box's M test<br>*p* = 0.112 | **Two-Way mixed ANOVA**<br>Interaction F(1,14) = 55.763 , *p* = <0.001 , partial $\eta^2$ = 0.799<br>Marker (W) F(1,14) = 112.21 , *p* = <0.001 , partial $\eta^2$ = 0.889<br>Stress (B) F(1,14) = 67.6 , *p* = <0.001 , partial $\eta^2$ = 0.828<br><br>post hoc (Bonferroni)<br>Vglut Unstressed x Stressed *p* = <0.001<br>Vgat Unstressed x Stressed *p* = 0.233 |
| **Figure<br>1K** | **Shapiro-Wilk test**<br>Unstressed Vglut *p* = 0.139<br>Stressed Vglut *p* = 0.350<br>Unstressed Vgat *p* = 0.851<br>Stressed Vgat *p* = 0.318<br>**Sample Size**<br>Unstressed *n* = 8<br>Stressed *n* = 8 | **Levene's test**<br>Vglut *p* = 0.198<br>Vgat *p* = 0.608<br><br>Box's M test<br>*p* = 0.657 | **Two-Way mixed ANOVA**<br>Interaction F(1,14) = 7.8963 , *p* = 0.013 , partial $\eta^2$ = 0.345<br>Marker (W) F(1,14) = 7.8963 , *p* = 0.013 , partial $\eta^2$ = 0.345<br>Stress (B) F(1,14) = 12.165 , *p* = 0.003 , partial $\eta^2$ = 0.448<br><br>post hoc (Bonferroni)<br>Vglut Unstressed x Stressed *p* = <0.001<br>Vgat Unstressed x Stressed *p* = 0.864 |
| **Figure<br>2E** | **Shapiro-Wilk test**<br>Total interaction time<br>mCherry OFF *p* = 0.061<br>eNPHR OFF *p* = 0.511<br>mCherry ON *p* = 0.623<br>eNPHR ON *p* = 0.716 | **Levene's test**<br>OFF *p* = 0.358<br>ON *p* = 0.441<br><br>Box's M test<br>*p* = 0.365 | **Two-Way mixed ANOVA**<br>Interaction F(1,14) = 4.528 , *p* = 0.052 , partial $\eta^2$ = 0.244<br>Light (W) F(1,14) = 14.698 , *p* = 0.002 , partial $\eta^2$ = 0.512<br>Opsin (B) F(1,14) = 18.049 , *p* = <0.001 , partial $\eta^2$ = 0.563<br><br>post hoc (Bonferroni)<br>mCherry ON x OFF *p* = 0.248<br>eNPHR ON x OFF *p* = <0.001 |

| | | | |
|---|---|---|---|
| | Number of visits<br>mCherry OFF *p* = 0.795<br>eNPHR OFF *p* = 0.269<br>mCherry ON *p* = 0.040<br>eNPHR ON *p* = 0.358 | OFF *p* = 0.658<br>ON *p* = 0.450 | **Wilcoxon signed-rank test**<br>mCherry ON x OFF , *p* = 0.735 , *z* = -0.338<br>*r* = -0.085<br>eNPHR ON x OFF , *p* = 0.575 , *z* = -0.560<br>*r* = -0.140 |
| | Interaction time per visit<br>mCherry OFF *p* = 0.214<br>eNPHR OFF *p* = 0.744<br>mCherry ON *p* = 0.891<br>eNPHR ON *p* = 0.706 | OFF *p* = 0.656<br>ON *p* = 0.393<br>Box's M test<br>*p* = 0.531 | **Two-Way mixed ANOVA**<br>Interaction F(1,14) = 2.8073 , *p* = 0.116 , partial $\eta^2$ = 0.167<br>Light (W) F(1,14) = 8.7977 , *p* = 0.010 , partial $\eta^2$ = 0.386<br>Opsin (B) F(1,14) = 6.8209 , *p* = 0.021 , partial $\eta^2$ = 0.328<br>post hoc (Bonferroni)<br>mCherry ON x OFF *p* = 0.377<br>eNPHR ON x OFF *p* = 0.005 |
| | **Sample Size**<br>mCherry *n* = 8<br>eNPHR *n* = 8 | | |
| **Figure 2G** | **Shapiro-Wilk test**<br>Total interaction time<br>tdTomato OFF p = 0.860<br>Chrimson OFF p = 0.643<br>tdTomato ON p = 0.907<br>Chrimson ON p = 0.525 | **Levene's test**<br>OFF p = 0.032<br>ON p = 0.527 | **Wilcoxon signed-rank test**<br>tdTomato ON x OFF *p* = 0.674 *z* = 0.420<br>*r* = 0.105<br>Chrimson ON x OFF *p* = 0.012 *z* = 2.521<br>*r* = 0.630 |
| | Number of visits<br>tdTomato OFF *p* = 0.138<br>Chrimson OFF *p* = 0.744<br>tdTomato ON *p* = 0.102<br>Chrimson ON *p* = 0.435 | OFF *p* = 0.107<br>ON *p* = 0.627<br>Box's M test<br>*p* = 0.012 | **Two-Way mixed ANOVA**<br>Interaction F(1,14) = 0.1500 , *p* = 0.704 , partial $\eta^2$ = 0.011<br>Light (W) F(1,14) = 1.8462 , *p* = 0.196 , partial $\eta^2$ = 0.117<br>Opsin (B) F(1,14) = 0.0022 , *p* = 0.963 , partial $\eta^2$ = <0.001<br>post hoc (Bonferroni)<br>tdTomato ON x OFF *p* = 0.237<br>Chrimson ON x OFF *p* = 0.503 |
| | Interaction time per visit<br>tdTomato OFF *p* = 0.234<br>Chrimson OFF *p* = 0.071<br>tdTomato ON *p* = 0.650<br>Chrimson ON *p* = 0.383 | OFF *p* = 0.323<br>ON *p* = 0.034 | **Wilcoxon signed-rank test**<br>tdTomato ON x OFF *p* = 0.123 *z* = 1.542<br>*r* = 0.386<br>Chrimson ON x OFF *p* = 0.025 *z* = 2.240<br>*r* = 0.56 |
| | **Sample Size**<br>tdTomato *n* = 8<br>Chrimson *n* = 8 | | |
| **Figure 2M** | **Sample Size (# cells)**<br>Unstressed mCherry = 972<br>Stressed tdTomato = 945<br>**Sample Size ( #animals)**<br>tdTomato *n* = 8<br>mCherry *n* = 8 | **Expected Count <5**<br>0% | **Chi-square test of homogeneity**<br>tdtomato x mCherry *p* = 0.842 $\chi^2(2)$ = 0.343<br>Phi = 0.013 |
| **Figure 2N** | **Sample Size (# cells)**<br>Unstressed mCherry = 972<br>Stressed tdTomato = 945<br>**Sample Size ( #animals)**<br>tdTomato *n* = 8<br>mCherry *n* = 8 | **Expected Count <5**<br>0% | **Chi-square test of homogeneity**<br>tdtomato x mCherry *p* = <0.001 $\chi^2(2)$ = 24.7<br>Phi = 0.114 |
| **Figure 2O** | **Sample Size (# cells)**<br>Unstressed mCherry = 972<br>Unstressed eNPHR = 972<br>Stressed tdTomato = 945<br>Stressed Chrimson = 1042<br>**Sample Size ( #animals)**<br>tdTomato *n* = 8<br>Chrimson *n* = 8 | **Expected Count <5**<br>0% | **Chi-square test of homogeneity**<br>Unstressed mCherry ON x OFF *p* = 0.673<br>Unstressed eNPHR ON x OFF *p* = <0.001<br>Stressed tdTomato ON x OFF *p* = 0.320<br>Stressed Chrimson ON x OFF *p* = <0.001 |
| **Figure 3B** | **Sample Size (# cells)**<br>Unstressed mCherry = 972<br>Stressed tdTomato = 945<br>**Sample Size ( #animals)**<br>tdTomato *n* = 8<br>mCherry *n* = 8 | **Expected Count <5**<br>0% | **Chi-square test of homogeneity**<br>tdtomato x mCherry *p* = <0.001 $\chi^2(8)$ = 34.67<br>Phi = 0.134 |

| Figure | Sample Size / Normality | Expected Count / Variance | Test |
|---|---|---|---|
| **Figure** | **Sample Size (# cells)** | **Expected Count <5** | **Chi-square test of homogeneity** |
| **3D** | Unstressed mCherry = 972 | 0% | eNPHR x mCherry $p$ = <0.001 $\chi^2(8)$ = 35.49 |
| | Unstressed eNPHR = 972 | | Phi = 0.135 |
| | **Sample Size ( #animals)** | | |
| | mCherry $n$ = 8 | | |
| | eNPHR $n$ = 8 | | |
| **Figure** | **Sample Size (# cells)** | **Expected Count <5** | **Chi-square test of homogeneity** |
| **3E** | Stressed tdTomato = 972 | 0% | tdTomato x mCherry $p$ = <0.001 $\chi^2(8)$ = 34.29 |
| | Stressed Chrimson = 1042 | | Phi = 0.131 |
| | **Sample Size ( #animals)** | | |
| | tdTomato $n$ = 8 | | |
| | Chrimson $n$ = 8 | | |
| **Figure** | **Shapiro-Wilk test** | **Levene's test** | **Wilcoxon signed-rank test** |
| **4C** | Total interaction time | | |
| | Exploration OFF $p$ = 0.827 | OFF $p$ = 0.322 | Exploration ON x OFF $p$ = 0.401 z = 0.84 |
| | Interaction OFF $p$ = 0.290 | ON $p$ = 0.179 | $r$ = 0.210 |
| | Exploration ON $p$ = 0.046 | | Interaction ON x OFF $p$ = 0.036 z = 2.10 |
| | Interaction ON $p$ = 0.590 | | $r$ = 0.525 |
| | Number of visits | | **Two-Way mixed ANOVA** |
| | Exploration OFF $p$ = 0.530 | OFF $p$ = 0.817 | Interaction F(1,14) = 0.3836 , $p$ = 0.546 , partial $\eta^2$ = 0.027 |
| | Interaction OFF $p$ = 0.375 | ON $p$ = 0.260 | Light (W) F(1,14) = 1.1049 , $p$ = 0.311 , partial $\eta^2$ = 0.073 |
| | Exploration ON $p$ = 0.655 | | Condition (B) F(1,14) = 0.0843 , $p$ = 0.776 , partial $\eta^2$ = 0.006 |
| | Interaction ON $p$ = 0.708 | | |
| | | Box's M test | post hoc (Bonferroni) |
| | | $p$ = 0.749 | Exploration ON x OFF $p$ = 0.257 |
| | | | Interaction ON x OFF $p$ = 0.765 |
| | Interaction time per visit | | **Two-Way mixed ANOVA** |
| | Exploration OFF $p$ = 0.300 | OFF $p$ = 0.514 | Interaction F(1,14) = 9.772 , $p$ = 0.007 , partial $\eta^2$ = 0.411 |
| | Interaction OFF $p$ = 0.165 | ON $p$ = 0.460 | Light (W) F(1,14) = 4.8626 , $p$ = 0.045 , partial $\eta^2$ = 0.258 |
| | Exploration ON $p$ = 0.463 | | Condition (B) F(1,14) = 1.6049 , $p$ = 0.226 , partial $\eta^2$ = 0.103 |
| | Interaction ON $p$ = 0.388 | | |
| | **Sample Size** | Box's M test | post hoc (Bonferroni) |
| | Exploration $n$ = 8 | $p$ = 0.402 | Exploration ON x OFF $p$ = 0.525 |
| | Interaction $n$ = 8 | | Interaction ON x OFF $p$ = 0.002 |
| **Figure** | **Shapiro-Wilk test** | **Levene's test** | **Two-Way mixed ANOVA** |
| **4E** | Total interaction time | | |
| | Exploration OFF $p$ = 0.445 | OFF $p$ = 0.948 | Interaction F(1,14) = 10.735 , $p$ = 0.006 , partial $\eta^2$ = 0.434 |
| | Interaction OFF $p$ = 0.598 | ON $p$ = 0.641 | Light (W) F(1,14) = 13.664 , $p$ = 0.002 , partial $\eta^2$ = 0.494 |
| | Exploration ON $p$ = 0.795 | | Condition (B) F(1,14) = 13.766 , $p$ = 0.002 , partial $\eta^2$ = 0.496 |
| | Interaction ON $p$ = 0.631 | | |
| | | Box's M test | post hoc (Bonferroni) |
| | | $p$ = 0.983 | Exploration ON x OFF $p$ = 0.771 |
| | | | Interaction ON x OFF $p$ = <0.001 |
| | Number of visits | | **Two-Way mixed ANOVA** |
| | Exploration OFF $p$ = 0.275 | OFF $p$ = 0.498 | Interaction F(1,14) = 2.236 , $p$ = 0.157 , partial $\eta^2$ = 0.138 |
| | Interaction OFF $p$ = 0.886 | ON $p$ = 0.219 | Light (W) F(1,14) = 0.1076 , $p$ = 0.748 , partial $\eta^2$ = 0.008 |
| | Exploration ON $p$ = 0.407 | | Condition (B) F(1,14) = 0.9752 , $p$ = 0.340 , partial $\eta^2$ = 0.065 |
| | Interaction ON $p$ = 0.837 | | |
| | | Box's M test | post hoc (Bonferroni) |
| | | $p$ = 0.626 | Exploration ON x OFF $p$ = 0.423 |
| | | | Interaction ON x OFF $p$ = 0.218 |
| | Interaction time per visit | | **Two-Way mixed ANOVA** |
| | Exploration OFF $p$ = 0.191 | OFF $p$ = 0.389 | Interaction F(1,14) = 1.2811 , $p$ = 0.277 , partial $\eta^2$ = 0.084 |
| | Interaction OFF $p$ = 0.097 | ON $p$ = 0.490 | Light (W) F(1,14) = 6.1802 , $p$ = 0.026 , partial $\eta^2$ = 0.306 |
| | Exploration ON $p$ = 0.560 | | Condition (B) F(1,14) = 2.8065 , $p$ = 0.116 , partial $\eta^2$ = 0.167 |
| | Interaction ON $p$ = 0.331 | | |
| | **Sample Size** | Box's M test | post hoc (Bonferroni) |
| | Exploration $n$ = 8 | $p$ = 0.720 | Exploration ON x OFF $p$ = 0.355 |
| | Interaction $n$ = 8 | | Interaction ON x OFF $p$ = 0.023 |
| **Figure** | **Sample Size (# cells)** | **Expected Count <5** | **Chi-square test of homogeneity** |
| **4F** | eNPHR/Exploration = 1021 | 0% | Exploration eNPHR ON x OFF $p$ = 0.551 |
| | eNPHR/Interaction = 1111 | | Interaction eNPHR ON x OFF $p$ = 0.001 |
| | Chrimson/Exploration = 997 | | Exploration Chrimson ON x OFF $p$ = 0.327 |
| | Chrimson/Interaction = 998 | | Interaction Chrimson ON x OFF $p$ = 0.002 |

| | | | |
|---|---|---|---|
| | **Sample Size ( #animals)** | | |
| | eNPHR Exploration *n* = 8 | | |
| | eNPHR Interaction *n* = 8 | | |
| | Chrimson Exploration *n* = 8 | | |
| | Chrimson Interaction *n* = 8 | | |
| **Figure 5C** | **Shapiro-Wilk test** | **Levene's test** | **One-way ANOVA** |
| | EGFP expression level | | |
| | Week 1 *p* = 0.179 | *p* = 0.587 | Week (B) F(2,11) = 96.155 , *p* = <0.001 , partial $\eta^2$ = 0.955 |
| | Week 2 *p* = 0.319 | | |
| | Week 3 *p* = 0.460 | | post hoc (Bonferroni) |
| | | | Week 1 x Week 2 *p* = 0.039 |
| | | | Week 1 x Week 3 *p* = <0.001 |
| | | | Week 2 x Week 3 *p* = <0.001 |
| | GR expression level | | **Welch's ANOVA** |
| | Week 1 *p* = 0.703 | *p* = 0.001 | |
| | Week 2 *p* = 0.554 | | Week (B) F(2,5) = 86.756 , *p* = <0.001 , partial $\eta^2$ = 0.966 |
| | Week 3 *p* = 0.292 | | |
| | | | post hoc (Games-Howell) |
| | | | Week 1 x Week 2 *p* = 0.012 |
| | | | Week 1 x Week 3 *p* = 0.001 |
| | | | Week 2 x Week 3 *p* = 0.001 |
| | **Sample Size** | | |
| | Week 1 *n* = 4 | | |
| | Week 2 *n* = 4 | | |
| | Week 3 *n* = 4 | | |
| **Figure 5D** | **Shapiro-Wilk test** | **Levene's test** | **Two-Way ANOVA** |
| | Total interaction time | | |
| | Unstressed Control *p* = 0.787 | *p* = 0.302 | Interaction F(3,20) = 3.418 , *p* = 0.079 , partial $\eta^2$ = 0.146 |
| | Unstressed GR-KO *p* = 0.698 | | Stress (B) F(3,20) = 1.531 , *p* = 0.230 , partial $\eta^2$ = 0.071 |
| | Stressed Control *p* = 0.082 | | KO (B) F(3,20) = 7.821 , *p* = 0.011 , partial $\eta^2$ = 0.281 |
| | Stressed GR-KO *p* = 0.354 | | |
| | | | post hoc (Bonferroni) |
| | | | Unstressed Control x GR-KO *p* = 0.510 |
| | | | Stressed Control x GR-KO *p* = 0.004 |
| | Number of visits | | **Mann Whitney U test** |
| | Unstressed Control *p* = 0.135 | *p* = 0.044 | Unstressed Control x GR-KO *p* = 0.699 *U* = 15.00 |
| | Unstressed GR-KO *p* = 0.056 | | *r* = -0.12 *z* = -0.48 |
| | Stressed Control *p* = 0.261 | | Stressed Control x GR-KO *p* = 0.589 *U* = 14.00 |
| | Stressed GR-KO *p* = 0.727 | | *r* = -0.16 *z* = -0.64 |
| | Interaction time per visit | | **Two-Way ANOVA** |
| | Unstressed Control *p* = 0.996 | *p* = 0.475 | Interaction F(3,20) = 4.2815 , *p* = 0.052 , partial $\eta^2$ = 0.176 |
| | Unstressed GR-KO *p* = 0.600 | | Stress (B) F(3,20) = 3.1581 , *p* = 0.091 , partial $\eta^2$ = 0.136 |
| | Stressed Control *p* = 0.260 | | KO (B) F(3,20) = 9.2584 , *p* = 0.006 , partial $\eta^2$ = 0.316 |
| | Stressed GR-KO *p* = 0.395 | | |
| | | | post hoc (Bonferroni) |
| | | | Unstressed Control x GR-KO *p* = 0.499 |
| | | | Stressed Control x GR-KO *p* = 0.002 |
| | **Sample Size** | | |
| | Unstressed Control *n* = 6 | | |
| | Unstressed GR-KO *n* = 6 | | |
| | Stressed Control *n* = 6 | | |
| | Stressed GR-KO *n* = 6 | | |
| **Figure 5F** | **Shapiro-Wilk test** | **Levene's test** | **Two-Way ANOVA** |
| | AI | | |
| | Unstressed Control p = 0.669 | *p* = 0.309 | Interaction F(3,20) = 10.027 , *p* = 0.005 , partial $\eta^2$ = 0.334 |
| | Unstressed GR-KO p = 0.520 | | Stress (B) F(3,20) = 17.218 , *p* = <0.001 , partial $\eta^2$ = 0.463 |
| | Stressed Control p = 0.454 | | KO (B) F(3,20) = 7.720 , *p* = 0.012 , partial $\eta^2$ = 0.279 |
| | Stressed GR-KO p = 0.315 | | |
| | | | post hoc (Bonferroni) |
| | | | Unstressed Control x GR-KO *p* = 0.787 |
| | | | Stressed Control x GR-KO *p* = <0.001 |
| | | | Control Unstressed x Stressed *p* = <0.001 |
| | PrL | | **Two-Way ANOVA** |

| | | | |
|---|---|---|---|
| | Unstressed Control $p =$ 0.570 | $p =$ 0.231 | Interaction F(3,20) = 8.251 , $p =$ 0.009 , partial $\eta^2 =$ 0.292 |
| | Unstressed GR-KO $p =$ 0.767 | | Stress (B) F(3,20) = 26.178 , $p =$ <0.001 , partial $\eta^2 =$ 0.567 |
| | Stressed Control $p =$ 0.461 | | KO (B) F(3,20) = 9.619 , $p =$ 0.006 , partial $\eta^2 =$ 0.325 |
| | Stressed GR-KO $p =$ 0.320 | | |
| | **Sample Size** | | post hoc (Bonferroni) |
| | Unstressed Control $n =$ 6 | | Unstressed Control x GR-KO $p =$ 0.873 |
| | Unstressed GR-KO $n =$ 6 | | Stressed Control x GR-KO $p =$ <0.001 |
| | Stressed Control $n =$ 6 | | Control Unstressed x Stressed $p =$ <0.001 |
| | Stressed GR-KO $n =$ 6 | | |
| **Figure S1B** | **Shapiro-Wilk test** | **Levene's test** | **Student t-Test Independent samples, 2-tailed** |
| | Total interaction time | | |
| | Unstressed $p =$ 0.571 | $p =$ 0.326 | $t$ (14) = 0.918 , $p =$ 0.374 |
| | Stressed $p =$ 0.526 | | Cohen's d = 0.459 |
| | Number of visits | | **Student t-Test Independent samples, 2-tailed** |
| | Unstressed $p =$ 0.076 | $p =$ 0.840 | $t$ (14) = 0.0215 , $p =$ 0.983 |
| | Stressed $p =$ 0.176 | | Cohen's d = 0.0108 |
| | Interaction time per visit | | **Student t-Test Independent samples, 2-tailed** |
| | Unstressed $p =$ 0.566 | $p =$ 0.466 | $t$ (14) = 0.9261 , $p =$ 0.370 |
| | Stressed $p =$ 0.633 | | Cohen's d = 0.4631 |
| | **Sample Size** | | |
| | Unstressed $n =$ 8 | | |
| | Stressed $n =$ 8 | | |
| **Figure S1C** | **Shapiro-Wilk test** | **Levene's test** | **Student t-Test Independent samples, 2-tailed** |
| | Total interaction time | | |
| | Unstressed $p =$ 0.479 | $p =$ 0.572 | $t$ (14) = 0.3837 , $p =$ 0.707 |
| | Stressed $p =$ 0.704 | | Cohen's d = 0.1919 |
| | Number of visits | | **Student t-Test Independent samples, 2-tailed** |
| | Unstressed $p =$ 0.302 | $p =$ 0.548 | $t$ (14) = 0.1412 , $p =$ 0.890 |
| | Stressed $p =$ 0.364 | | Cohen's d = 0.0706 |
| | Interaction time per visit | | **Student t-Test Independent samples, 2-tailed** |
| | Unstressed $p =$ 0.813 | $p =$ 0.999 | $t$ (14) = 0.3646 , $p =$ 0.721 |
| | Stressed $p =$ 0.824 | | Cohen's d = 0.1823 |
| | **Sample Size** | | |
| | Unstressed $n =$ 8 | | |
| | Stressed $n =$ 8 | | |
| **Figure S1D** | **Shapiro-Wilk test** | **Levene's test** | **Student t-Test Independent samples, 2-tailed** |
| | PrL | | |
| | Unstressed $p =$ 0.413 | $p =$ 0.062 | $t$ (14) = 3.345 , $p =$ 0.005 |
| | Stressed $p =$ 0.820 | | Cohen's d = 1.672 |
| | **Sample Size** | | |
| | Unstressed $n =$ 8 | | |
| | Stressed $n =$ 8 | | |
| **Figure S1G** | **Shapiro-Wilk test** | | **Paired Student t-Test, 2-tailed** |
| | Area | | |
| | MO Male ♂ $p =$ 0.055 | | $t$ (3) = 2.375 , $p =$ 0.098 , Cohen's d = 1.188 |
| | VO Male ♂ $p =$ 0.628 | | $t$ (3) = 6.662 , $p =$ 0.007 , Cohen's d = 3.331 |
| | LO Male ♂ $p =$ 0.714 | | $t$ (3) = 12.880 , $p =$ 0.001 , Cohen's d = 6.440 |
| | AI Male ♂ $p =$ 0.233 | | $t$ (3) = 3.660 , $p =$ 0.035 , Cohen's d = 1.830 |
| | M1 Male ♂ $p =$ 0.186 | | $t$ (3) = 4.993 , $p =$ 0.015 , Cohen's d = 2.496 |
| | M2 Male ♂ $p =$ 0.074 | | $t$ (3) = 20.048 , $p =$ <0.001 , Cohen's d = 10.024 |
| | Cg Male ♂ $p =$ 0.597 | | $t$ (3) = 14.311 , $p =$ <0.001 , Cohen's d = 7.156 |
| | PrL Male ♂ $p =$ 0.515 | | $t$ (3) = -18.585 , $p =$ <0.001 , Cohen's d = -9.292 |
| | IL Male ♂ $p =$ 0.079 | | $t$ (3) = 3.884 , $p =$ 0.030 , Cohen's d = 1.942 |
| | AM Male ♂ $p =$ 0.293 | | $t$ (3) = 2.980 , $p =$ 0.059 , Cohen's d = 1.490 |
| | VM Male ♂ $p =$ 0.156 | | $t$ (3) = 4.079 , $p =$ 0.027 , Cohen's d = 2.040 |
| | RSA Male ♂ $p =$ 0.115 | | $t$ (3) = 3.155 , $p =$ 0.051 , Cohen's d = 1.577 |
| | V2 Male ♂ $p =$ 0.130 | | $t$ (3) = 13.462 , $p =$ <0.001 , Cohen's d = 6.731 |
| | V1 Male ♂ $p =$ 0.262 | | $t$ (3) = 7.577 , $p =$ 0.005 , Cohen's d = 3.789 |
| | MD Male ♂ $p =$ 0.572 | | $t$ (3) = 37.044 , $p =$ <0.001 , Cohen's d = 18.522 |
| | RE Male ♂ $p =$ 0.144 | | $t$ (3) = 2.307 , $p =$ 0.104 , Cohen's d = 1.154 |
| | DEN Male ♂ $p =$ 0.345 | | $t$ (3) = 2.359 , $p =$ 0.099 , Cohen's d = 1.180 |
| | BMA Male ♂ $p =$ 0.567 | | $t$ (3) = 7.920 , $p =$ 0.004 , Cohen's d = 3.960 |
| | BLA Male ♂ $p =$ 0.283 | | $t$ (3) = 2.550 , $p =$ 0.084 , Cohen's d = 1.275 |
| | ECT Male ♂ $p =$ 0.301 | | $t$ (3) = 3.146 , $p =$ 0.051 , Cohen's d = 1.573 |
| | PRh Male ♂ $p =$ 0.354 | | $t$ (3) = 3.060 , $p =$ 0.055 , Cohen's d = 1.530 |
| | LEnt Male ♂ $p =$ 0.860 | | $t$ (3) = 6.663 , $p =$ 0.007 , Cohen's d = 3.332 |

| Figure | Area | Group | Shapiro-Wilk test | Levene's | Area | Test | Statistic | p | Effect size |
|---|---|---|---|---|---|---|---|---|---|
| | Pir | Male ♂ | $p$ = 0.245 | | | $t$ (3) | = 4.522 | $p$ = 0.020 | Cohen's d = 2.261 |
| | S | Male ♂ | $p$ = 0.354 | | | $t$ (3) | = 2.831 | $p$ = 0.066 | Cohen's d = 1.416 |
| | CA1 | Male ♂ | $p$ = 0.065 | | | $t$ (3) | = 2.602 | $p$ = 0.080 | Cohen's d = 1.301 |
| | VTA | Male ♂ | $p$ = 0.105 | | | $t$ (3) | = 1.360 | $p$ = 0.267 | Cohen's d = 0.680 |
| | MO | Female ♀ | $p$ = 0.368 | | | $t$ (3) | = 2.441 | $p$ = 0.092 | Cohen's d = 1.220 |
| | VO | Female ♀ | $p$ = 0.572 | | | $t$ (3) | = 8.560 | $p$ = 0.003 | Cohen's d = 4.280 |
| | LO | Female ♀ | $p$ = 0.650 | | | $t$ (3) | = 2.011 | $p$ = 0.138 | Cohen's d = 1.005 |
| | AI | Female ♀ | $p$ = 0.679 | | | $t$ (3) | = 2.891 | $p$ = 0.063 | Cohen's d = 1.445 |
| | M1 | Female ♀ | $p$ = 0.580 | | | $t$ (3) | = 6.088 | $p$ = 0.009 | Cohen's d = 3.044 |
| | M2 | Female ♀ | $p$ = 0.114 | | | $t$ (3) | = 5.394 | $p$ = 0.012 | Cohen's d = 2.697 |
| | Cg | Female ♀ | $p$ = 0.558 | | | $t$ (3) | = 6.209 | $p$ = 0.008 | Cohen's d = 3.105 |
| | PrL | Female ♀ | $p$ = 0.794 | | | $t$ (3) | = -5.914 | $p$ = 0.010 | Cohen's d = -2.957 |
| | IL | Female ♀ | $p$ = 0.936 | | | $t$ (3) | = 3.296 | $p$ = 0.046 | Cohen's d = 1.648 |
| | AM | Female ♀ | $p$ = 0.237 | | | $t$ (3) | = 4.029 | $p$ = 0.027 | Cohen's d = 2.015 |
| | VM | Female ♀ | $p$ = 0.798 | | | $t$ (3) | = 3.050 | $p$ = 0.055 | Cohen's d = 1.525 |
| | RSA | Female ♀ | $p$ = 0.192 | | | $t$ (3) | = 3.530 | $p$ = 0.039 | Cohen's d = 1.765 |
| | V2 | Female ♀ | $p$ = 0.513 | | | $t$ (3) | = 10.226 | $p$ = 0.002 | Cohen's d = 5.113 |
| | V1 | Female ♀ | $p$ = 0.279 | | | $t$ (3) | = 2.901 | $p$ = 0.062 | Cohen's d = 1.450 |
| | S2 | Female ♀ | $p$ = 0.644 | | | $t$ (3) | = 7.165 | $p$ = 0.006 | Cohen's d = 3.582 |
| | MD | Female ♀ | $p$ = 0.458 | | | $t$ (3) | = 10.561 | $p$ = 0.002 | Cohen's d = 5.280 |
| | RE | Female ♀ | $p$ = 0.104 | | | $t$ (3) | = 1.311 | $p$ = 0.281 | Cohen's d = 0.655 |
| | DEN | Female ♀ | $p$ = 0.985 | | | $t$ (3) | = 2.782 | $p$ = 0.069 | Cohen's d = 1.391 |
| | BMA | Female ♀ | $p$ = 0.561 | | | $t$ (3) | = 5.063 | $p$ = 0.015 | Cohen's d = 2.532 |
| | BLA | Female ♀ | $p$ = 0.629 | | | $t$ (3) | = 2.805 | $p$ = 0.068 | Cohen's d = 1.403 |
| | ECT | Female ♀ | $p$ = 0.304 | | | $t$ (3) | = 2.834 | $p$ = 0.066 | Cohen's d = 1.417 |
| | PRh | Female ♀ | $p$ = 0.056 | | | $t$ (3) | = 2.647 | $p$ = 0.077 | Cohen's d = 1.323 |
| | LEnt | Female ♀ | $p$ = 0.367 | | | $t$ (3) | = 5.561 | $p$ = 0.011 | Cohen's d = 2.781 |
| | Pir | Female ♀ | $p$ = 0.761 | | | $t$ (3) | = 2.857 | $p$ = 0.065 | Cohen's d = 1.429 |
| | S | Female ♀ | $p$ = 0.662 | | | $t$ (3) | = 3.855 | $p$ = 0.031 | Cohen's d = 1.928 |
| | CA1 | Female ♀ | $p$ = 0.374 | | | $t$ (3) | = 3.312 | $p$ = 0.045 | Cohen's d = 1.656 |
| | DR | Female ♀ | $p$ = 0.062 | | | $t$ (3) | = 4.265 | $p$ = 0.024 | Cohen's d = 2.132 |
| | VTA | Female ♀ | $p$ = 0.676 | | | $t$ (3) | = 1.982 | $p$ = 0.142 | Cohen's d = 0.991 |
| | | | | | | **Wilcoxon signed-rank test** | | | |
| | S2 | Male ♂ | $p$ = 0.017 | | | Ipsi x Contra | | $p$ = 0.068 | $z$ = 1.826 |
| | | | | | | | | | $r$ = 0.646 |
| | DR | Male ♂ | $p$ = 0.049 | | | Ipsi x Contra | | $p$ = 0.068 | $z$ = 1.825 |
| | | **Sample Size** | | | | | | | $r$ = 0.645 |
| | | Male | $n$ = 4 | | | | | | |
| | | Female | $n$ = 4 | | | | | | |
| **Figure** | **Shapiro-Wilk test** | | | **Levene's** | **Student t-Test Independent samples, 2-tailed** | | | | |
| **S1H** | Area | | | | Area | | | | |
| | MO | Unstressed | $p$ = 0.420 | $p$ = 0.054 | MO | $t$ (14) | = 0.4489 | $p$ = 0.660 | Cohen's d = 0.224 |
| | VO | Unstressed | $p$ = 0.951 | $p$ = 0.078 | VO | $t$ (14) | = -0.2627 | $p$ = 0.797 | Cohen's d = -0.131 |
| | LO | Unstressed | $p$ = 0.661 | $p$ = 0.799 | LO | $t$ (14) | = -0.1005 | $p$ = 0.921 | Cohen's d = -0.050 |
| | AI | Unstressed | $p$ = 0.896 | $p$ = 0.076 | AI | $t$ (14) | = 7.2071 | $p$ = <0.001 | Cohen's d = 3.604 |
| | M1 | Unstressed | $p$ = 0.242 | $p$ = 0.815 | M1 | $t$ (14) | = 0.7030 | $p$ = 0.494 | Cohen's d = 0.352 |
| | M2 | Unstressed | $p$ = 0.846 | $p$ = 0.272 | M2 | $t$ (14) | = 0.5075 | $p$ = 0.620 | Cohen's d = 0.254 |
| | Cg | Unstressed | $p$ = 0.679 | $p$ = 0.944 | Cg | $t$ (14) | = 1.1508 | $p$ = 0.269 | Cohen's d = 0.575 |
| | IL | Unstressed | $p$ = 0.431 | $p$ = 0.763 | IL | $t$ (14) | = 2.1112 | $p$ = 0.053 | Cohen's d = 1.056 |
| | AM | Unstressed | $p$ = 0.711 | $p$ = 0.692 | AM | $t$ (14) | = 1.0403 | $p$ = 0.316 | Cohen's d = 0.520 |
| | VM | Unstressed | $p$ = 0.358 | $p$ = 0.523 | VM | $t$ (14) | = -1.0253 | $p$ = 0.323 | Cohen's d = -0.513 |
| | RSA | Unstressed | $p$ = 0.849 | $p$ = 0.904 | RSA | $t$ (14) | = -0.6915 | $p$ = 0.501 | Cohen's d = -0.346 |
| | V2 | Unstressed | $p$ = 0.061 | $p$ = 0.059 | V2 | $t$ (14) | = 1.1576 | $p$ = 0.266 | Cohen's d = 0.579 |
| | V1 | Unstressed | $p$ = 0.384 | $p$ = 0.682 | V1 | $t$ (14) | = 0.3981 | $p$ = 0.697 | Cohen's d = 0.199 |
| | S2 | Unstressed | $p$ = 0.409 | $p$ = 0.264 | S2 | $t$ (14) | = 0.2366 | $p$ = 0.816 | Cohen's d = 0.118 |
| | MD | Unstressed | $p$ = 0.613 | $p$ = 0.239 | MD | $t$ (14) | = -1.8322 | $p$ = 0.088 | Cohen's d = -0.916 |
| | RE | Unstressed | $p$ = 0.294 | $p$ = 0.435 | RE | $t$ (14) | = -0.2929 | $p$ = 0.774 | Cohen's d = -0.146 |
| | DEN | Unstressed | $p$ = 0.562 | $p$ = 0.165 | DEN | $t$ (14) | = 0.4471 | $p$ = 0.662 | Cohen's d = 0.224 |
| | BMA | Unstressed | $p$ = 0.578 | $p$ = 0.822 | BMA | $t$ (14) | = -0.9130 | $p$ = 0.377 | Cohen's d = -0.456 |
| | BLA | Unstressed | $p$ = 0.596 | $p$ = 0.051 | BLA | $t$ (14) | = 2.8513 | $p$ = 0.013 | Cohen's d = 1.426 |
| | ECT | Unstressed | $p$ = 0.955 | $p$ = 0.299 | ECT | $t$ (14) | = 0.2354 | $p$ = 0.817 | Cohen's d = 0.118 |
| | PRh | Unstressed | $p$ = 0.085 | $p$ = 0.246 | PRh | $t$ (14) | = 0.0110 | $p$ = 0.991 | Cohen's d = 0.006 |
| | LEnt | Unstressed | $p$ = 0.523 | $p$ = 0.940 | LEnt | $t$ (14) | = -1.9096 | $p$ = 0.077 | Cohen's d = -0.955 |
| | Pir | Unstressed | $p$ = 0.237 | $p$ = 0.245 | Pir | $t$ (14) | = -0.4888 | $p$ = 0.633 | Cohen's d = -0.244 |
| | CA1 | Unstressed | $p$ = 0.723 | $p$ = 0.536 | CA1 | $t$ (14) | = 0.5443 | $p$ = 0.595 | Cohen's d = 0.272 |
| | DR | Unstressed | $p$ = 0.903 | $p$ = 0.114 | DR | $t$ (14) | = -0.1640 | $p$ = 0.872 | Cohen's d = -0.082 |
| | VTA | Unstressed | $p$ = 0.624 | $p$ = 0.517 | VTA | $t$ (14) | = -0.6980 | $p$ = 0.496 | Cohen's d = -0.349 |
| | MO | Stressed | $p$ = 0.880 | | | | | | |
| | VO | Stressed | $p$ = 0.782 | | | | | | |

| | | | |
|---|---|---|---|
| | LO Stressed $p$ = 0.130 | | |
| | AI Stressed $p$ = 0.561 | | |
| | M1 Stressed $p$ = 0.886 | | |
| | M2 Stressed $p$ = 0.411 | | |
| | Cg Stressed $p$ = 0.134 | | |
| | IL Stressed $p$ = 0.721 | | |
| | AM Stressed $p$ = 0.909 | | |
| | VM Stressed $p$ = 0.841 | | |
| | RSA Stressed $p$ = 0.463 | | |
| | V2 Stressed $p$ = 0.879 | | |
| | V1 Stressed $p$ = 0.476 | | |
| | S2 Stressed $p$ = 0.964 | | |
| | MD Stressed $p$ = 0.308 | | |
| | RE Stressed $p$ = 0.109 | | |
| | DEN Stressed $p$ = 0.234 | | |
| | BMA Stressed $p$ = 0.152 | | |
| | BLA Stressed $p$ = 0.354 | | |
| | ECT Stressed $p$ = 0.719 | | |
| | PRh Stressed $p$ = 0.389 | | |
| | LEnt Stressed $p$ = 0.204 | | |
| | Pir Stressed $p$ = 0.636 | | |
| | CA1 Stressed $p$ = 0.330 | | |
| | DR Stressed $p$ = 0.784 | | |
| | VTA Stressed $p$ = 0.337 | | |
| | | | **Mann Whitney U test** |
| | S Unstressed $p$ = 0.026 | $p$ = 0.932 | Unstressed x Stressed $p$ = 0.195 $U$ = 45.00 |
| | S Stressed $p$ = 0.888 | | $r$ = 0.341 $z$ = 1.37 |
| | **Sample Size** | | |
| | Unstressed $n$ = 8 | | |
| | Stressed $n$ = 8 | | |
| **Figure S1J** | **Shapiro-Wilk test** | **Levene's test** | **Two-Way mixed ANOVA** |
| | Male ♂ Vglut $p$ = 0.967 | Vglut $p$ = 0.769 | Interaction F(1,10) = 0.3396 , $p$ = 0.573 , partial $\eta^2$ = 0.033 |
| | Male ♂ Vgat $p$ = 0.796 | Vgat $p$ = 0.126 | Marker (W) F(1,10) = 829.12 , $p$ = <0.001 , partial $\eta^2$ = 0.988 |
| | Female ♀ Vglut $p$ = 0.922 | | Sex (B) F(1,10) = 0.0003 , $p$ = 0.987 , partial $\eta^2$ = <0.001 |
| | Female ♀ Vgat $p$ = 0.153 | | |
| | | Box's M test | post hoc (Bonferroni) |
| | | $p$ = 0.555 | Male ♂ Vglut x Vgat $p$ = <0.001 |
| | | | Female ♀ Vglut x Vgat $p$ = <0.001 |
| | **Sample Size** | | |
| | Unstressed $n$ = 6 | | |
| | Stressed $n$ = 6 | | |
| **Figure S2** | **NA** | **NA** | **NA** |
| | **Sample Size** | | |
| | Female B6J mice $n$ = 4 | | |
| **Figure S3A** | **Shapiro-Wilk test** | **Sample Size (# cells)** | **Wilcoxon signed rank test (One Sample)** |
| | Modulation index | | Against 0.5 |
| | Unstressed mCherry $p$ = <0.001 | N= 512 | $p$ = 0.692 , $z$ = 0.396 $r$ = 0.018 |
| | Unstressed eNPHR $p$ = <0.001 | N= 618 | $p$ = <0.001 , $z$ = -8.283 $r$ = -0.333 |
| | Stressed tdTomato $p$ = 0.001 | N= 708 | $p$ = 0.714 , $z$ = -0.367 $r$ = -0.014 |
| | Stressed Chrimson $p$ = <0.001 | N= 658 | $p$ = <0.001 , $z$ = 6.784 $r$ = 0.264 |
| | **Sample Size ( # animals)** | | |
| | Unstressed mCherry $n$ = 6 | | |
| | Unstressed eNPHR $n$ = 6 | | |
| | Stressed tdTomato $n$ = 6 | | |
| | Stressed Chrimson $n$ = 6 | | |
| **Figure S3A (Modulation index)** | **Shapiro-Wilk test** | **Levene's test** | **Mann Whitney U test** |
| | Modulation index | | |
| | Unstressed mCherry $p$ = <0.001 | $p$ = 0.029 | mCherry x eNPHR $p$ = <0.001 $U$ = >1200 |
| | Unstressed eNPHR $p$ = <0.001 | | $r$ = 0.163 $z$ = 5.46 |
| | Modulation index | | **Mann Whitney U test** |
| | Stressed tdTomato $p$ = 0.001 | $p$ = 0.175 | tdTomato x Chrimson $p$ = <0.001 $U$ = >1200 |
| | Stressed Chrimson $p$ = <0.001 | | $r$ = 0.134 $z$ = 4.97 |
| | **Sample Size ( # animals)** | **Sample Size (# cells)** | |
| | Unstressed mCherry $n$ = 6 | N= 512 | |
| | Unstressed eNPHR $n$ = 6 | N= 618 | |
| | Stressed tdTomato $n$ = 6 | N= 708 | |
| | Stressed Chrimson $n$ = 6 | N= 658 | |

| | | | |
|---|---|---|---|
| **Figure** | **Shapiro-Wilk test** | **Sample Size (# cells)** | **Wilcoxon signed rank test (One Sample)** |
| **S3B** | Relative Peak Amplitude | | Against 1.0 |
| | Unstressed mCherry *p* = <0.001 | N= 484 | *p* = 0.235 , *z* = -1.188 *r* = -0.054 |
| | Unstressed eNPHR *p* = <0.001 | N= 609 | *p* = 0.008 , *z* = -2.640 *r* = -0.107 |
| | Stressed tdTomato *p* = <0.001 | N= 672 | *p* = 0.492 , *z* = 0.687 *r* = 0.027 |
| | Stressed Chrimson *p* = <0.001 | N= 633 | *p* = 0.006 , *z* = 2.742 *r* = 0.109 |
| | **Sample Size ( # animals)** | | |
| | Unstressed mCherry *n* = 6 | | |
| | Unstressed eNPHR *n* = 6 | | |
| | Stressed tdTomato *n* = 6 | | |
| | Stressed Chrimson *n* = 6 | | |
| **Figure** | **Shapiro-Wilk test** | **Levene's test** | **Two-Way mixed ANOVA** |
| **S3D** | Total interaction time | | |
| | mCherry OFF *p* = 0.177 | OFF *p* = 0.976 | Interaction F(1,14) = 0.0909 , *p* = 0.768 , partial $\eta^2$ = 0.006 |
| | eNPHR OFF *p* = 0.521 | ON *p* = 0.591 | Light (W) F(1,14) = 2.4759 , *p* = 0.138 , partial $\eta^2$ = 0.150 |
| | mCherry ON *p* = 0.970 | | Opsin (B) F(1,14) = 4.3559 , *p* = 0.056 , partial $\eta^2$ = 0.237 |
| | eNPHR ON *p* = 0.540 | | |
| | | Box's M test | post hoc (Bonferroni) |
| | | *p* = 0.827 | mCherry ON x OFF *p* = 0.384 |
| | | | eNPHR ON x OFF *p* = 0.206 |
| | Number of visits | | **Two-Way mixed ANOVA** |
| | mCherry OFF *p* = 0.490 | OFF *p* = 0.456 | Interaction F(1,14) = 0.0020 , *p* = 0.965 , partial $\eta^2$ = 1E-04 |
| | eNPHR OFF *p* = 0.065 | ON *p* = 0.856 | Light (W) F(1,14) = 1.1265 , *p* = 0.306 , partial $\eta^2$ = 0.074 |
| | mCherry ON *p* = 0.603 | | Opsin (B) F(1,14) = 0.2207 , *p* = 0.646 , partial $\eta^2$ = 0.016 |
| | eNPHR ON *p* = 0.772 | | |
| | | Box's M test | post hoc (Bonferroni) |
| | | *p* = 0.327 | mCherry ON x OFF *p* = 0.484 |
| | | | eNPHR ON x OFF *p* = 0.447 |
| | Interaction time per visit | | **Two-Way mixed ANOVA** |
| | mCherry OFF *p* = 0.328 | OFF *p* = 0.573 | Interaction F(1,14) = 0.2059 , *p* = 0.657 , partial $\eta^2$ = 0.014 |
| | eNPHR OFF *p* = 0.675 | ON *p* = 0.100 | Light (W) F(1,14) = 0.0000 , *p* = 0.999 , partial $\eta^2$ = <0.001 |
| | mCherry ON *p* = 0.201 | | Opsin (B) F(1,14) = 5.5826 , *p* = 0.033 , partial $\eta^2$ = 0.285 |
| | eNPHR ON *p* = 0.583 | | |
| | | Box's M test | post hoc (Bonferroni) |
| | | *p* = 0.357 | mCherry ON x OFF *p* = 0.754 |
| | | | eNPHR ON x OFF *p* = 0.752 |
| | **Sample Size** | | |
| | mCherry *n* = 8 | | |
| | eNPHR *n* = 8 | | |
| **Figure** | **Shapiro-Wilk test** | **Levene's test** | **Two-Way mixed ANOVA** |
| **S3E** | Total interaction time | | |
| | tdTomato OFF *p* = 0.310 | OFF *p* = 0.791 | Interaction F(1,14) = 0.7861 , *p* = 0.390 , partial $\eta^2$ = 0.053 |
| | Chrimson OFF *p* = 0.755 | ON *p* = 0.734 | Light (W) F(1,14) = 0.0000 , *p* = 0.998 , partial $\eta^2$ = <0.001 |
| | tdTomato ON *p* = 0.116 | | Opsin (B) F(1,14) = 0.5529 , *p* = 0.469 , partial $\eta^2$ = 0.038 |
| | Chrimson ON *p* = 0.894 | | |
| | | Box's M test | post hoc (Bonferroni) |
| | | *p* = 0.735 | tdTomato ON x OFF *p* = 0.540 |
| | | | Chrimson ON x OFF *p* = 0.542 |
| | Number of visits | | **Wilcoxon signed-rank test** |
| | tdTomato OFF *p* = 0.285 | OFF *p* = 0.916 | tdTomato ON x OFF , *p* = 0.499 , *z* = 0.676 |
| | Chrimson OFF *p* = 0.314 | ON *p* = 0.380 | *r* = 0.169 |
| | tdTomato ON *p* = 0.008 | | Chrimson ON x OFF , *p* = 0.091 , *z* = 0.091 |
| | Chrimson ON *p* = 0.198 | | *r* = 0.023 |
| | | Box's M test | |
| | | *p* = 0.217 | |
| | Interaction time per visit | | **Two-Way mixed ANOVA** |
| | tdTomato OFF *p* = 0.336 | OFF *p* = 0.885 | Interaction F(1,14) = 2.8794 , *p* = 0.112 , partial $\eta^2$ = 0.171 |
| | Chrimson OFF *p* = 0.719 | ON *p* = 0.863 | Light (W) F(1,14) = 0.6306 , *p* = 0.440 , partial $\eta^2$ = 0.043 |
| | tdTomato ON *p* = 0.802 | | Opsin (B) F(1,14) = 0.1131 , *p* = 0.742 , partial $\eta^2$ = 0.008 |
| | Chrimson ON *p* = 0.528 | | |
| | | Box's M test | post hoc (Bonferroni) |
| | | *p* = 0.870 | tdTomato ON x OFF *p* = 0.534 |
| | | | Chrimson ON x OFF *p* = 0.100 |
| | **Sample Size** | | |

| | | | |
|---|---|---|---|
| | tdTomato $n$ = 8<br>Chrimson $n$ = 8 | | |
| **Figure S3G** | **Shapiro-Wilk test**<br>Total interaction time<br>Unstressed $p$ = 0.809<br>Unstressed + Opsin $p$ = 0.749<br>Stressed $p$ = 0.753<br>Stressed + Opsin $p$ = 0.622<br><br>**Sample Size**<br>Unstressed $n$ = 8<br>Unstressed + Opsin $n$ = 8<br>Stressed $n$ = 8<br>Stressed + Opsin $n$ = 8 | **Levene's test**<br>$p$ = 0.662 | **Two-Way ANOVA**<br>Interaction F(3,32) = 37.3786 , $p$ = <0.001 , partial $\eta^2$ = 0.572<br>Stress (B) F(3,32) = 0.0986 , $p$ = 0.756 , partial $\eta^2$ = 0.004<br>Opsin+ (B) F(3,32) = 0.0246 , $p$ = 0.876 , partial $\eta^2$ = <0.001<br><br>post hoc (Bonferroni)<br>Unstressed No Opsin x Opsin $p$ = <0.001<br>Stressed No Opsin x Opsin $p$ = <0.001 |
| **Figure S4A** | **Sample Size (# cells)**<br>Unstressed mCherry = 1165<br>Stressed tdTomato = 1196<br>**Sample Size ( # animals)**<br>tdTomato $n$ = 8<br>mCherry $n$ = 8 | **Expected Count <5**<br>0% | **Chi-square test of homogeneity**<br>tdtomato x mCherry $p$ = 0.758 $\chi^2(2)$ = 0.555<br>Phi = 0.015 |
| **Figure S4B** | **Sample Size (# cells)**<br>Unstressed mCherry = 1165<br>Stressed tdTomato = 1196<br>**Sample Size ( # animals)**<br>tdTomato $n$ = 8<br>mCherry $n$ = 8 | **Expected Count <5**<br>0% | **Chi-square test of homogeneity**<br>tdtomato x mCherry $p$ = 0.102 $\chi^2(2)$ = 4.568<br>Phi = 0.044 |
| **Figure S4C** | **Sample Size (# cells)**<br>Unstressed mCherry = 1165<br>Unstressed eNPHR = 1175<br>Stressed tdTomato = 1196<br>Stressed Chrimson = 1087<br>**Sample Size ( # animals)**<br>Unstressed mCherry $n$ = 8<br>Unstressed eNPHR $n$ = 8<br>Stressed tdTomato $n$ = 8<br>Stressed Chrimson $n$ = 8 | **Expected Count <5**<br>0% | **Chi-square test of homogeneity**<br>Unstressed mCherry ON x OFF $p$ = 0.489<br>Unstressed eNPHR ON x OFF $p$ = 0.206<br>Stressed tdTomato ON x OFF $p$ = 0.509<br>Stressed Chrimson ON x OFF $p$ = 0.451 |
| **Figure S4D** | **Sample Size (# cells)**<br>Unstressed mCherry = 1165<br>Unstressed eNPHR = 1175<br>Stressed tdTomato = 1196<br>Stressed Chrimson = 1087<br>**Sample Size ( # animals)**<br>Unstressed mCherry $n$ = 8<br>Unstressed eNPHR $n$ = 8<br>Stressed tdTomato $n$ = 8<br>Stressed Chrimson $n$ = 8 | **Expected Count <5**<br>0% | **Chi-square test of homogeneity**<br>Unstressed mCherry ON x OFF $p$ = 0.471<br>Unstressed eNPHR ON x OFF $p$ = 0.916<br>Stressed tdTomato ON x OFF $p$ = 0.347<br>Stressed Chrimson ON x OFF $p$ = 0.517 |
| **Figure S4E** | **Sample Size (# cells)**<br>Unstressed mCherry = 972<br>Unstressed eNPHR = 972<br>Stressed tdTomato = 945<br>Stressed Chrimson = 1042<br>**Sample Size ( # animals)**<br>Unstressed mCherry $n$ = 8<br>Unstressed eNPHR $n$ = 8<br>Stressed tdTomato $n$ = 8<br>Stressed Chrimson $n$ = 8 | **Expected Count <5**<br>0% | **Chi-square test of homogeneity**<br>Unstressed mCherry ON x OFF $p$ = 0.306<br>Unstressed eNPHR ON x OFF $p$ = 0.697<br>Stressed tdTomato ON x OFF $p$ = 0.606<br>Stressed Chrimson ON x OFF $p$ = 0.239 |
| **Figure S5B** | **Shapiro-Wilk test**<br>Total interaction time $p$ = 0.696<br>Number of visits $p$ = 0.462<br>Interaction time per visit $p$ = 0.914<br>**Sample Size**<br>Virgin with SILA $n$ = 8<br>Virgin without SILA $n$ = 8 | | **Paired Student t-Test, 2-tailed**<br>$t$ (7) = 5.559 , $p$ = 0.001 Cohen's d = 1.965<br>$t$ (7) = 2.457 , $p$ = 0.044 Cohen's d = 0.869<br>$t$ (7) = 2.811 , $p$ = 0.026 Cohen's d = 0.994 |
| **Figure S5C** | **Shapiro-Wilk test**<br>Total interaction time $p$ = 0.733<br>Number of visits $p$ = 0.658<br><br>Interaction time per visit $p$ = 0.041<br>**Sample Size** | | **Paired Student t-Test, 2-tailed**<br>$t$ (7) = 3.570 , $p$ = 0.009 Cohen's d = 1.262<br>$t$ (7) = 1.014 , $p$ = 0.344 Cohen's d = 0.359<br><br>**Wilcoxon signed-rank test**<br>Novel x Familiar $p$ = 0.017 z = 2.38<br>$r$ = 0.841 |

| | | | |
|---|---|---|---|
| | Virgin with SILA *n* = 8<br>Virgin without SILA *n* = 8 | | |
| **Figure<br>S5D** | **Sample Size (# cells)**<br>Virgin with SILA = 1085<br>Virgin without SILA = 1088<br>**Sample Size ( # animals)**<br>Virgin with SILA *n* = 8<br>Virgin without SILA *n* = 8 | **Expected Count <5**<br>0% | **Chi-square test of homogeneity**<br>Empty cage Unstressed x Stressed *p* = 0.403<br><br>Mouse cage Unstressed x Stressed *p* = 0.330 |
| **Figure<br>S5E** | **Sample Size (# cells)**<br>Virgin with SILA = 1086<br>Virgin without SILA = 1076<br>**Sample Size ( # animals)**<br>Virgin with SILA *n* = 8<br>Virgin without SILA *n* = 8 | **Expected Count <5**<br>0% | **Chi-square test of homogeneity**<br>Familiar Unstressed x Stressed *p* = 0.708<br><br>Novel Unstressed x Stressed *p* = 0.954 |
| **Figure<br>S6B** | **Shapiro-Wilk test**<br><br>Unstressed Vglut *p* = 0.941<br>Stressed Vglut *p* = 0.425<br><br>Unstressed Vgat *p* = 0.645<br>Stressed Vgat *p* = 0.336<br>**Sample Size**<br>Unstressed Vglut *n* = 8<br>Stressed Vglut *n* = 8<br>Unstressed Vgat *n* = 8<br>Stressed Vgat *n* = 8 | **Levene's test**<br><br>Vglut *p* = 0.094<br><br><br>Vgat *p* = 0.046 | **Student t-Test Independent samples, 2-tailed**<br><br>Vglut *t* (14) = -4.083 , *p* = 0.001<br>Cohen's d = -2.042<br><br>**Welch t-Test Independent samples, 2-tailed**<br>Vgat *t* (14) = 0.978 , *p* = 0.349<br>Cohen's d = 0.489 |
| **Figure<br>S6D** | **Shapiro-Wilk test**<br><br>Male ♀ Vglut *p* = 0.716<br>Female ♀ Vglut *p* = 0.140<br>Male ♀ Vgat *p* = 0.396<br>Female ♀ Vgat *p* = 0.936<br><br>**Sample Size**<br>Male ♀ Vglut *n* = 6<br>Female ♀ Vglut *n* = 6<br>Male ♀ Vgat *n* = 6<br>Female ♀ Vgat *n* = 6 | **Levene's test**<br><br>Vglut *p* = 0.811<br>Vgat *p* = 0.880<br><br><br>Box's M test<br>*p* = 0.950 | **Two-Way mixed ANOVA**<br><br>Interaction F(1,14) = 1.4555 , *p* = 0.255 , partial $\eta^2$ = 0.127<br>Marker (W) F(1,14) = 559.76 , *p* = <0.001 , partial $\eta^2$ = 0.982<br>Sex (B) F(1,14) = 1.3831 , *p* = 0.267 , partial $\eta^2$ = 0.122<br><br>post hoc (Bonferroni)<br>Male ♂ Vglut x Vgat *p* = <0.001<br>Female ♀ Vglut x Vgat *p* = <0.001 |
| **Figure<br>S6G** | **Shapiro-Wilk test**<br>Total interaction time<br>eNPHR OFF *p* = 0.562<br>Chrimson OFF *p* = 0.782<br>eNPHR ON *p* = 0.975<br>Chrimson ON *p* = 0.583<br><br><br><br><br>Number of visits<br>eNPHR OFF *p* = 0.499<br>Chrimson OFF *p* = 0.253<br>eNPHR ON *p* = 0.714<br>Chrimson ON *p* = 0.304<br><br>Interaction time per visit<br>eNPHR OFF *p* = 0.104<br>Chrimson OFF *p* = 0.815<br>eNPHR ON *p* = 0.415<br>Chrimson ON *p* = 0.546<br>**Sample Size**<br>Unstressed eNPHR *n* = 8<br>Stressed Chrimson *n* = 8 | **Levene's test**<br><br>OFF *p* = 0.981<br>ON *p* = 0.214<br><br><br>Box's M test<br>*p* = 0.465<br><br><br><br>OFF *p* = 0.244<br>ON *p* = 0.030<br><br><br><br><br>OFF *p* = 0.773<br>ON *p* = 0.007 | **Two-Way mixed ANOVA**<br><br>Interaction F(1,14) = 0.0407 , *p* = 0.843 , partial $\eta^2$ = 0.003<br>Light (W) F(1,14) = 0.4551 , *p* = 0.511 , partial $\eta^2$ = 0.031<br>Opsin (B) F(1,14) = 0.1751 , *p* = 0.682 , partial $\eta^2$ = 0.012<br><br>post hoc (Bonferroni)<br>eNPHR ON x OFF *p* = 0.743<br>Chrimson ON x OFF *p* = 0.545<br><br>**Wilcoxon Signed-rank test**<br>eNPHR ON x OFF *p* = 0.779 z = -0.280<br>*r* = -0.07<br>Chrimson ON x OFF *p* = 0.735 z = -0.018<br>*r* = -0.004<br><br>**Wilcoxon Signed-rank test**<br>eNPHR ON x OFF *p* = 0.779 z = -0.280<br>*r* = -0.07<br>Chrimson ON x OFF *p* = 0.208 z = -1.26<br>*r* = -0.315 |

| Figure | Shapiro-Wilk test | Levene's test | Two-Way mixed ANOVA |
|---|---|---|---|
| **Figure**<br>**S6H** | Total interaction time<br>eNPHR OFF $p$ = 0.985<br>Chrimson OFF $p$ = 0.370<br>eNPHR ON $p$ = 0.865<br>Chrimson ON $p$ = 0.302 | OFF $p$ = 0.490<br>ON $p$ = 0.849<br>Box's M test<br>$p$ = 0.428 | Interaction F(1,14) = 0.0024 , $p$ = 0.962 , partial $\eta^2$ = <0.001<br>Light (W) F(1,14) = 0.2363 , $p$ = 0.634 , partial $\eta^2$ = 0.017<br>Opsin (B) F(1,14) = 26.033 , $p$ = <0.001 , partial $\eta^2$ = 0.650<br>post hoc (Bonferroni)<br>eNPHR ON x OFF $p$ = 0.711<br>Chrimson ON x OFF $p$ = 0.762 |
| | Number of visits<br>eNPHR OFF $p$ = 0.661<br>Chrimson OFF $p$ = 0.656<br>eNPHR ON $p$ = 0.132<br>Chrimson ON $p$ = 0.566 | OFF $p$ = 0.097<br>ON $p$ = 0.543<br>Box's M test<br>$p$ = 0.742 | **Two-Way mixed ANOVA**<br>Interaction F(1,14) = 0.015 , $p$ = 0.904 , partial $\eta^2$ = 0.001<br>Light (W) F(1,14) = 0.1738 , $p$ = 0.683 , partial $\eta^2$ = 0.012<br>Opsin (B) F(1,14) = 0.9663 , $p$ = 0.342 , partial $\eta^2$ = 0.065<br>post hoc (Bonferroni)<br>eNPHR ON x OFF $p$ = 0.838<br>Chrimson ON x OFF $p$ = 0.709 |
| | Interaction time per visit<br>eNPHR OFF $p$ = 0.526<br>Chrimson OFF $p$ = 0.237<br>eNPHR ON $p$ = 0.947<br>Chrimson ON $p$ = 0.922<br>**Sample Size**<br>Unstressed eNPHR $n$ = 8<br>Stressed Chrimson $n$ = 8 | OFF $p$ = 0.577<br>ON $p$ = 0.888<br>Box's M test<br>$p$ = 0.293 | **Two-Way mixed ANOVA**<br>Interaction F(1,14) = 0.0007 , $p$ = 0.980 , partial $\eta^2$ = <0.001<br>Light (W) F(1,14) = 0.3774 , $p$ = 0.549 , partial $\eta^2$ = 0.026<br>Opsin (B) F(1,14) = 20.889 , $p$ = <0.001 , partial $\eta^2$ = 0.599<br>post hoc (Bonferroni)<br>eNPHR ON x OFF $p$ = 0.684<br>Chrimson ON x OFF $p$ = 0.658 |
| **Figure**<br>**S6I** | **Sample Size (# cells)**<br>Unstressed eNPHR = 1032<br>Stressed Chrimson = 1217<br>**Sample Size (# animals)**<br>Unstressed eNPHR $n$ = 8<br>Stressed Chrimson $n$ = 8 | **Expected Count <5**<br>0% | **Chi-square test of homogeneity**<br>eNPHR Empty ON x OFF $p$ = 0.965<br>Chrimson Empty ON x OFF $p$ = 0.175<br>eNPHR Mouse ON x OFF $p$ = 0.692<br>Chrimson Mouse ON x OFF $p$ = 0.509 |
| **Figure**<br>**S6J** | **Sample Size (# cells)**<br>Unstressed eNPHR = 1082<br>Stressed Chrimson = 1152<br>**Sample Size (# animals)**<br>Unstressed eNPHR $n$ = 8<br>Stressed Chrimson $n$ = 8 | **Expected Count <5**<br>0% | **Chi-square test of homogeneity**<br>eNPHR Familiar ON x OFF $p$ = 0.865<br>Chrimson Familiar ON x OFF $p$ = 0.416<br>eNPHR Novel ON x OFF $p$ = 0.936<br>Chrimson Novel ON x OFF $p$ = 0.847 |
| **Figure**<br>**S7** | **Sample Size (# cells)**<br>Unstressed eNPHR = 972<br>Stressed Chrimson = 1042<br>**Sample Size (# animals)**<br>Unstressed eNPHR $n$ = 8<br>Stressed Chrimson $n$ = 8 | **Expected Count <5**<br>0% | **Chi-square test of homogeneity**<br>Chrimson x eNPHR $p$ = 0.846 |
| **Figure**<br>**S8A** | **Sample Size (# cells)**<br>Unstressed mCherry = 1165<br>Stressed tdTomato = 1196<br>Unstressed eNPHR = 1175<br>Stressed Chrimson = 1087<br>**Sample Size (# animals)**<br>Unstressed mCherry $n$ = 8<br>Stressed tdTomato $n$ = 8<br>Unstressed eNPHR $n$ = 8<br>Stressed Chrimson $n$ = 8 | **Expected Count <5**<br>0% | **Chi-square test of homogeneity**<br>eNPHR x mCherry $p$ = 0.658 $\chi^2(8)$ = 5.9<br>Phi = 0.05<br>Chrimson x tdTomato $p$ = 0.609 $\chi^2(8)$ = 6.342<br>Phi = 0.053 |
| **Figure**<br>**S8B** | **Sample Size (# cells)**<br>Unstressed mCherry = 1165<br>Stressed tdTomato = 1196<br>Unstressed eNPHR = 1175<br>Stressed Chrimson = 1087<br>**Sample Size (# animals)**<br>Unstressed mCherry $n$ = 8<br>Stressed tdTomato $n$ = 8<br>Unstressed eNPHR $n$ = 8<br>Stressed Chrimson $n$ = 8 | **Expected Count <5**<br>0% | **Chi-square test of homogeneity**<br>eNPHR x mCherry $p$ = 0.318 $\chi^2(8)$ = 9.291<br>Phi = 0.063<br>Chrimson x tdTomato $p$ = 0.304 $\chi^2(8)$ = 9.467<br>Phi = 0.064 |

| Figure | | | |
|---|---|---|---|
| **Figure S8C** | **Sample Size (# cells)** | **Expected Count <5** | **Chi-square test of homogeneity** |
| | Unstressed mCherry = 972 | 0% | eNPHR x mCherry $p$ = 0.745 $\chi^2(8)$ = 5.12 |
| | Stressed tdTomato = 945 | | Phi = 0.051 |
| | Unstressed eNPHR = 972 | | |
| | Stressed Chrimson = 1042 | | Chrimson x tdTomato $p$ = 0.165 $\chi^2(8)$ = 11.71 |
| | **Sample Size (# animals)** | | Phi = 0.077 |
| | Unstressed mCherry $n$ = 8 | | |
| | Stressed tdTomato $n$ = 8 | | |
| | Unstressed eNPHR $n$ = 8 | | |
| | Stressed Chrimson $n$ = 8 | | |
| **Figure S9B** | **Shapiro-Wilk test** | **Levene's test** | **Two-Way mixed ANOVA** |
| | Total interaction time | | |
| | mCherry OFF $p$ = 0.877 | OFF $p$ = 0.433 | Interaction F(1,14) = 2.125 , $p$ = 0.167 , partial $\eta^2$ = 0.132 |
| | eNPHR OFF $p$ = 0.387 | ON $p$ = 0.603 | Light (W) F(1,14) = 0.066 , $p$ = 0.801 , partial $\eta^2$ = 0.005 |
| | mCherry ON $p$ = 0.729 | | Opsin (B) F(1,14) = 0.010 , $p$ = 0.922 , partial $\eta^2$ = <0.001 |
| | eNPHR ON $p$ = 0.700 | | |
| | | Box's M test | post hoc (Bonferroni) |
| | | $p$ = 0.702 | mCherry ON x OFF $p$ = 0.410 |
| | | | eNPHR ON x OFF $p$ = 0.245 |
| | Number of visits | | **Two-Way mixed ANOVA** |
| | mCherry OFF $p$ = 0.633 | OFF $p$ = 0.0905 | Interaction F(1,14) = 0.5086 , $p$ = 0.487 , partial $\eta^2$ = 0.035 |
| | eNPHR OFF $p$ = 0.872 | ON $p$ = 0.9041 | Light (W) F(1,14) = 0.5361 , $p$ = 0.476 , partial $\eta^2$ = 0.037 |
| | mCherry ON $p$ = 0.889 | | Opsin (B) F(1,14) = 0.0255 , $p$ = 0.875 , partial $\eta^2$ = 0.002 |
| | eNPHR ON $p$ = 0.506 | | |
| | | Box's M test | post hoc (Bonferroni) |
| | | $p$ = 0.3328 | mCherry ON x OFF $p$ = 0.989 |
| | | | eNPHR ON x OFF $p$ = 0.324 |
| | Interaction time per visit | | **Related Sampes Sign Test** |
| | mCherry OFF $p$ = 0.686 | OFF $p$ = 0.0182 | |
| | eNPHR OFF $p$ = 0.706 | ON $p$ = 0.0853 | mCherry ON x OFF $p$ = 0.289 $z$ = -1.061 |
| | mCherry ON $p$ = 0.466 | | $r$ = -0.265 |
| | eNPHR ON $p$ = 0.725 | | eNPHR ON x OFF $p$ = 1.000 $z$ = 0.00 |
| | **Sample Size** | | $r$ = 0.00 |
| | mCherry $n$ = 8 | | |
| | eNPHR $n$ = 8 | | |
| **Figure S9C** | **Shapiro-Wilk test** | **Levene's test** | **Two-Way mixed ANOVA** |
| | Total interaction time | | |
| | mCherry OFF p = 0.470 | OFF p = 0.568 | Interaction F(1,14) = 1.794 , $p$ = 0.202 , partial $\eta^2$ = 0.114 |
| | eNPHR OFF p = 0.879 | ON p = 0.110 | Light (W) F(1,14) = 0.651 , $p$ = 0.433 , partial $\eta^2$ = 0.044 |
| | mCherry ON p = 0.715 | | Opsin (B) F(1,14) = 0.015 , $p$ = 0.903 , partial $\eta^2$ = 0.001 |
| | eNPHR ON p = 0.924 | | |
| | | Box's M test | post hoc (Bonferroni) |
| | | $p$ = 0.391 | mCherry ON x OFF $p$ = 0.712 |
| | | | eNPHR ON x OFF $p$ = 0.151 |
| | Number of visits | | **Two-Way mixed ANOVA** |
| | mCherry OFF $p$ = 0.051 | OFF $p$ = 0.7447 | Interaction F(1,14) = 1.2134 , $p$ = 0.289 , partial $\eta^2$ = 0.080 |
| | eNPHR OFF $p$ = 0.209 | ON $p$ = 0.3767 | Light (W) F(1,14) = 0.1482 , $p$ = 0.706 , partial $\eta^2$ = 0.010 |
| | mCherry ON $p$ = 0.910 | | Opsin (B) F(1,14) = 0.241 , $p$ = 0.631 , partial $\eta^2$ = 0.017 |
| | eNPHR ON $p$ = 0.677 | | |
| | | Box's M test | post hoc (Bonferroni) |
| | | $p$ = 0.950 | mCherry ON x OFF $p$ = 0.311 |
| | | | eNPHR ON x OFF $p$ = 0.620 |
| | Interaction time per visit | | **Two-Way mixed ANOVA** |
| | mCherry OFF $p$ = 0.756 | OFF $p$ = 0.1665 | Interaction F(1,14) = 0.5804 , $p$ = 0.459 , partial $\eta^2$ = 0.040 |
| | eNPHR OFF $p$ = 0.973 | ON $p$ = 0.3728 | Light (W) F(1,14) = 2.9871 , $p$ = 0.106 , partial $\eta^2$ = 0.176 |
| | mCherry ON $p$ = 0.302 | | Opsin (B) F(1,14) = 0.7484 , $p$ = 0.402 , partial $\eta^2$ = 0.051 |
| | eNPHR ON $p$ = 0.190 | | |
| | | Box's M test | post hoc (Bonferroni) |
| | **Sample Size** | $p$ = 0.0446 | mCherry ON x OFF $p$ = 0.506 |
| | mCherry $n$ = 8 | | eNPHR ON x OFF $p$ = 0.100 |
| | eNPHR $n$ = 8 | | |

| **Figure** | **Shapiro-Wilk test** | **Levene's test** | **Two-Way mixed ANOVA** |
|---|---|---|---|
| **S9D** | Total interaction time | | |
| | mCherry OFF *p* = 0.723 | OFF *p* = 0.606 | Interaction F(1,14) = 11.390 , *p* = 0.005 , partial $\eta^2$ = 0.449 |
| | eNPHR OFF *p* = 0.487 | ON *p* = 0.642 | Light (W) F(1,14) = 2.045 , *p* = 0.175 , partial $\eta^2$ = 0.127 |
| | mCherry ON *p* = 0.707 | | Opsin (B) F(1,14) = 10.686 , *p* = 0.006 , partial $\eta^2$ = 0.433 |
| | eNPHR ON *p* = 0.274 | | |
| | | Box's M test | post hoc (Bonferroni) |
| | | *p* = 0.431 | mCherry ON x OFF *p* = 0.191 |
| | | | eNPHR ON x OFF *p* = 0.004 |
| | Number of visits | | **Wilcoxon signed-rank test** |
| | mCherry OFF *p* = 0.014 | OFF *p* = 0.514 | mCherry ON x OFF *p* = 0.161 *z* = 1.400 |
| | eNPHR OFF *p* = 0.883 | ON *p* = 0.7806 | *r* = 0.495 |
| | mCherry ON *p* = 0.155 | | eNPHR ON x OFF *p* = 0.674 *z* = 0.420 |
| | eNPHR ON *p* = 0.689 | | *r* = 0.148 |
| | Interaction time per visit | | **Two-Way mixed ANOVA** |
| | mCherry Light OFF p = 0.051 | Empty p = 0.249 | Interaction F(1,14) = 4.518 , p = 0.052 , partial $\eta^2$ = 0.244 |
| | eNPHR Light OFF p = 0.817 | Mouse p = 0.725 | Light (W) F(1,14) = 2.983 , p = 0.106 , partial $\eta^2$ = 0.176 |
| | mCherry Light ON p = 0.952 | | Opsin (B) F(1,14) = 4.782 , p = 0.046 , partial $\eta^2$ = 0.255 |
| | eNPHR Light ON p = 0.121 | | |
| | **Sample Size** | Box's M test | post hoc (Bonferroni) |
| | mCherry *n* = 8 | p = 0.907 | mCherry ON x OFF p = 0.782 |
| | eNPHR *n* = 8 | | eNPHR ON x OFF p = 0.016 |
| **Figure** | **Shapiro-Wilk test** | **Levene's test** | **Wilcoxon signed-rank test** |
| **S9E** | Total interaction time | | |
| | mCherry OFF *p* = 0.024 | OFF *p* = 0.200 | mCherry ON x OFF *p* = 0.263 *z* = 1.120 |
| | eNPHR OFF *p* = 0.335 | ON *p* = 0.745 | *r* = 0.396 |
| | mCherry ON *p* = 0.007 | | eNPHR ON x OFF *p* = 0.017 *z* = -2.380 |
| | eNPHR ON *p* = 0.476 | | *r* = -0.841 |
| | Number of visits | | **Wilcoxon signed-rank test** |
| | mCherry OFF *p* = 0.195 | OFF *p* = 0.351 | mCherry ON x OFF *p* = 0.484 *z* = 0.700 |
| | eNPHR OFF *p* = 0.491 | ON *p* = 0.253 | *r* = 0.247 |
| | mCherry ON *p* = 0.042 | | eNPHR ON x OFF *p* = 0.674 *z* = -0.420 |
| | eNPHR ON *p* = 0.622 | | *r* = -0.148 |
| | Interaction time per visit | | **Two-Way mixed ANOVA** |
| | mCherry OFF *p* = 0.863 | Empty *p* = 0.280 | Interaction F(1,14) = 21.425 , *p* = <0.001 , partial $\eta^2$ = 0.605 |
| | eNPHR OFF *p* = 0.329 | Mouse *p* = 0.928 | Light (W) F(1,14) = 13.94 , *p* = 0.002 , partial $\eta^2$ = 0.499 |
| | mCherry ON *p* = 0.560 | | Opsin (B) F(1,14) = 3.11 , *p* = 0.100 , partial $\eta^2$ = 0.182 |
| | eNPHR ON *p* = 0.668 | | |
| | **Sample Size** | Box's M test | post hoc (Bonferroni) |
| | mCherry *n* = 8 | *p* = 0.7559 | mCherry ON x OFF *p* = 0.537 |
| | eNPHR *n* = 8 | | eNPHR ON x OFF *p* = <0.001 |
| **Figure** | **Sample Size (# cells)** | **Expected Count <5** | **Chi-square test of homogeneity** |
| **S10A** | Virgin ♂ mCherry = 1060 | 0% | Virgin Male ♂ mCherry ON x OFF *p* = 0.532 |
| | Virgin ♂ eNPHR = 1052 | | Virgin Male ♂ eNPHR ON x OFF *p* = 0.623 |
| | Virgin ♀ mCherry = 1088 | | Virgin Female ♀ mCherry ON x OFF *p* = 0.248 |
| | Virgin ♀ eNPHR = 1113 | | Virgin Female ♀ eNPHR ON x OFF *p* = 0.150 |
| | **Sample Size (# animals)** | | |
| | Virgin ♂ mCherry *n* = 8 | | |
| | Virgin ♂ eNPHR *n* = 8 | | |
| | Virgin ♀ mCherry *n* = 8 | | |
| | Virgin ♀ eNPHR *n* = 8 | | |
| **Figure** | **Sample Size (# cells)** | **Expected Count <5** | **Chi-square test of homogeneity** |
| **S10B** | Virgin ♂ mCherry = 1060 | 0% | Virgin Male ♂ mCherry ON x OFF *p* = 0.632 |
| | Virgin ♂ eNPHR = 1052 | | Virgin Male ♂ eNPHR ON x OFF *p* = 0.923 |
| | Virgin ♀ mCherry = 1088 | | Virgin Female ♀ mCherry ON x OFF *p* = 0.400 |
| | Virgin ♀ eNPHR = 1113 | | Virgin Female ♀ eNPHR ON x OFF *p* = 0.587 |
| | **Sample Size (# animals)** | | |
| | Virgin ♂ mCherry *n* = 8 | | |
| | Virgin ♂ eNPHR *n* = 8 | | |
| | Virgin ♀ mCherry *n* = 8 | | |
| | Virgin ♀ eNPHR *n* = 8 | | |
| **Figure** | **Sample Size (# cells)** | **Expected Count <5** | **Chi-square test of homogeneity** |
| **S10C** | Virgin ♂ mCherry = 1167 | 0% | Virgin Male ♂ mCherry ON x OFF *p* = 0.448 |

| | | | |
|---|---|---|---|
| | Virgin ♂ eNPHR = 1095<br>Virgin ♀ mCherry = 1076<br>Virgin ♀ eNPHR = 1067<br>**Sample Size (# animals)**<br>Virgin ♂ mCherry *n* = 8<br>Virgin ♂ eNPHR *n* = 8<br>Virgin ♀ mCherry *n* = 8<br>Virgin ♀ eNPHR *n* = 8 | | Virgin Male ♂ eNPHR ON x OFF *p* = 0.136<br>Virgin Female ♀ mCherry ON x OFF *p* = 0.170<br>Virgin Female ♀ eNPHR ON x OFF *p* = 0.452 |
| **Figure S10D** | **Sample Size (# cells)**<br>Virgin ♂ mCherry = 1167<br>Virgin ♂ eNPHR = 1095<br>Virgin ♀ mCherry = 1076<br>Virgin ♀ eNPHR = 1067<br>**Sample Size (# animals)**<br>Virgin ♂ mCherry *n* = 8<br>Virgin ♂ eNPHR *n* = 8<br>Virgin ♀ mCherry *n* = 8<br>Virgin ♀ eNPHR *n* = 8 | **Expected Count <5**<br>0% | **Chi-square test of homogeneity**<br>Virgin Male ♂ mCherry ON x OFF *p* = 0.625<br>Virgin Male ♂ eNPHR ON x OFF *p* = <0.001<br>Virgin Female ♀ mCherry ON x OFF *p* = 0.267<br>Virgin Female ♀ eNPHR ON x OFF *p* = <0.001 |
| **Figure S11A** | **Sample Size (# cells)**<br>Unstressed mCherry = 1167<br>Unstressed eNPHR = 1095<br>**Sample Size (# animals)**<br>Unstressed mCherry *n* = 8<br>Unstressed eNPHR *n* = 8 | **Expected Count <5**<br>0% | **Chi-square test of homogeneity**<br>eNPHR x mCherry *p* = <0.001 $\chi^2(8)$ = 32.28<br>Phi = 0.119 |
| **Figure S11B** | **Sample Size (# cells)**<br>Unstressed mCherry = 1076<br>Unstressed eNPHR = 1067<br>**Sample Size (# animals)**<br>Unstressed mCherry *n* = 8<br>Unstressed eNPHR *n* = 8 | **Expected Count <5**<br>0% | **Chi-square test of homogeneity**<br>eNPHR x mCherry *p* = <0.001 $\chi^2(8)$ = 26.87<br>Phi = 0.112 |
| **Figure S12B** | **Shapiro-Wilk test**<br>Discrimination Index<br>Unstressed Opsin+ OFF *p* = 0.092<br>Unstressed Opsin+ ON *p* = 0.591<br>Unstressed Opsin- OFF *p* = 0.756<br>Unstressed Opsin- ON *p* = 0.461<br>Stressed Opsin+ OFF *p* = 0.807<br>Stressed Opsin+ ON *p* = 0.485<br>Stressed Opsin- OFF *p* = 0.275<br>Stressed Opsin- ON *p* = 0.307<br>**Sample Size**<br>Unstressed Opsin+ *n* = 8<br>Unstressed Opsin- *n* = 8<br>Stressed Opsin+ *n* = 8<br>Stressed Opsin- *n* = 8<br>Unstressed Opsin+ = eNPHR<br>Stressed Opsin+ = Chrimson | **Levene's test**<br>OFF *p* = 0.543<br>ON *p* = 0.184<br>Box's M test<br>*p* = 0.264 | **Three-Way mixed ANOVA**<br>Interaction F(1,28) = 0.102 , *p* = 0.752 , partial $\eta^2$ = 0.004<br>Light (W) F(1,28) = 0.002 , *p* = 0.969 , partial $\eta^2$ = <0.001<br>Stress (B) F(1,28) = 0.069 , *p* = 0.795 , partial $\eta^2$ = 0.002<br>Opsin (B) F(1,28) = 2.683 , *p* = 0.113 , partial $\eta^2$ = 0.087<br>post hoc (Bonferroni)<br>Unstressed Opsin+ OFF x ON *p* = 0.579<br>Unstressed Opsin- OFF x ON *p* = 0.752<br>Stressed Opsin+ OFF x ON *p* = 0.504<br>Stressed Opsin- OFF x ON *p* = 0.408 |
| **Figure S12C** | **Sample Size (# cells)**<br>Unstressed mCherry = 976<br>Unstressed eNPHR = 1013<br>Stressed tdTomato = 1007<br>Stressed Chrimson = 1030<br>**Sample Size (# animals)**<br>Unstressed mCherry *n* = 8<br>Unstressed eNPHR *n* = 8<br>Stressed tdTomato *n* = 8<br>Stressed Chrimson *n* = 8 | **Expected Count <5**<br>0% | **Chi-square test of homogeneity**<br>Unstressed mCherry ON x OFF *p* = 0.755<br>Unstressed eNPHR ON x OFF *p* = 0.408<br>Stressed tdTomato ON x OFF *p* = 0.824<br>Stressed Chrimson ON x OFF *p* = 0.475 |
| **Figure S12D** | **Sample Size (# cells)**<br>Unstressed mCherry = 976<br>Unstressed eNPHR = 1013<br>Stressed tdTomato = 1007<br>Stressed Chrimson = 1030 | **Expected Count <5**<br>0% | **Chi-square test of homogeneity**<br>Unstressed mCherry ON x OFF *p* = 0.795<br>Unstressed eNPHR ON x OFF *p* = 0.296<br>Stressed tdTomato ON x OFF *p* = 0.563<br>Stressed Chrimson ON x OFF *p* = 0.704 |

| | | | |
|---|---|---|---|
| | **Sample Size (# animals)**<br>Unstressed mCherry *n* = 8<br>Unstressed eNPHR *n* = 8<br>Stressed tdTomato *n* = 8<br>Stressed Chrimson *n* = 8 | | |
| **Figure S12F** | **Shapiro-Wilk test**<br>Total exploration time<br>Unstressed *p* = 0.449<br>Stressed *p* = 0.797<br><br>Number of visits<br>Unstressed *p* = 0.674<br>Stressed *p* = 0.843<br><br>Exploration time per visit<br>Unstressed *p* = 0.253<br>Stressed *p* = 0.768<br>**Sample Size**<br>Unstressed *n* = 8<br>Stressed *n* = 8 | **Levene's test**<br><br>*p* = 0.331<br><br><br><br>*p* = 0.163<br><br><br><br>*p* = 0.494 | **Student t-Test Independent samples, 2-tailed**<br><br>*t* (14) = -1.057 , *p* = 0.308<br>Cohen's d = -0.529<br><br>**Student t-Test Independent samples, 2-tailed**<br>*t* (14) = -0.045 , *p* = 0.965<br>Cohen's d = -0.023<br><br>**Student t-Test Independent samples, 2-tailed**<br>*t* (14) = -0.843 , *p* = 0.414<br>Cohen's d = -0.421 |
| **Figure S12H** | **Shapiro-Wilk test**<br>Total exploration time<br>Unstressed *p* = 0.085<br>Stressed *p* = 0.665<br><br>Number of visits<br>Unstressed *p* = 0.301<br>Stressed *p* = 0.962<br><br>Exploration time per visit<br>Unstressed *p* = 0.135<br>Stressed *p* = 0.024<br>**Sample Size**<br>Unstressed *n* = 8<br>Stressed *n* = 8 | **Levene's test**<br><br>*p* = 0.006<br><br><br><br>*p* = 0.285<br><br><br><br>*p* = 0.328 | **Welch t-Test Independent samples, 2-tailed**<br><br>*t* (14) = 0.719 , *p* = 0.487<br>Cohen's d = 0.360<br><br>**Student t-Test Independent samples, 2-tailed**<br>*t* (14) = -0.006 , *p* = 0.995<br>Cohen's d = -0.003<br><br>**Mann Whitney U test**<br>Unstressed x Stressed *p* = 0.130 *U* = 17.00<br>*r* = -0.394 *z* = -1.58 |
| **Figure S12I** | **Shapiro-Wilk test**<br>Total exploration time<br>Unstressed *p* = 0.800<br>Stressed *p* = 0.966<br><br>Number of visits<br>Unstressed *p* = 0.409<br>Stressed *p* = 0.901<br><br>Exploration time per visit<br>Unstressed *p* = 0.512<br>Stressed *p* = 0.589<br>**Sample Size**<br>Unstressed *n* = 8<br>Stressed *n* = 8 | **Levene's test**<br><br>*p* = 0.704<br><br><br><br>*p* = 0.118<br><br><br><br>*p* = 0.349 | **Student t-Test Independent samples, 2-tailed**<br><br>*t* (14) = 0.221 , *p* = 0.828<br>Cohen's d = 0.111<br><br>**Student t-Test Independent samples, 2-tailed**<br>*t* (14) = 1.715 , *p* = 0.108<br>Cohen's d = 0.857<br><br>**Student t-Test Independent samples, 2-tailed**<br>*t* (14) = -1.284 , *p* = 0.220<br>Cohen's d = -0.642 |
| **Figure S12K** | **Shapiro-Wilk test**<br>Time<br>Unstressed OFF *p* = 0.268<br>Stressed OFF *p* = 0.372<br>Unstressed ON *p* = 0.090<br>Stressed ON *p* = 0.482<br><br><br>**Sample Size**<br>Unstressed *n* = 8<br>Stressed *n* = 8 | **Levene's test**<br><br>OFF *p* = 0.858<br>ON *p* = 0.642<br><br><br>Box's M test<br>*p* = 0.097 | **Two-Way mixed ANOVA**<br><br>Interaction F(1,14) = 0.554 , *p* = 0.469 , partial $\eta^2$ = 0.038<br>Time (W) F(1,14) = 0.378 , *p* = 0.549 , partial $\eta^2$ = 0.026<br>Stress (B) F(1,14) = 0.785 , *p* = 0.391 , partial $\eta^2$ = 0.053<br><br>post hoc (Bonferroni)<br>Unstressed Empty x Mouse *p* = 0.353<br>Stressed Empty x Mouse *p* = 0.928 |
| **Figure S13B** | **Shapiro-Wilk test**<br>Total interaction time<br>Exploration OFF *p* = 0.633<br>Interaction OFF *p* = 0.047<br>Exploration ON *p* = 0.173 | **Levene's test**<br><br>OFF *p* = 0.931<br>ON *p* = 0.045 | **Wilcoxon signed-rank test**<br><br>Exploration ON x OFF *p* = 0.779 *z* = 0.280<br>*r* = 0.07<br>Interaction ON x OFF *p* = 0.674 *z* = 0.420 |

| | | | |
|---|---|---|---|
| | Interaction ON *p* = 0.334 | | *r* = 0.105 |
| | Number of visits | | **Wilcoxon signed-rank test** |
| | Exploration OFF *p* = 0.092 | OFF *p* = 0.029 | Exploration ON x OFF *p* = 0.735 *z* = -0.338 |
| | Interaction OFF *p* = 0.138 | ON *p* = 0.349 | *r* = -0.085 |
| | Exploration ON *p* = 0.498 | | Interaction ON x OFF *p* = 0.779 *z* = -0.280 |
| | Interaction ON *p* = 0.242 | | *r* = -0.07 |
| | Interaction time per visit | | **Two-Way mixed ANOVA** |
| | Exploration OFF *p* = 0.199 | OFF *p* = 0.4668 | Interaction F(1,14) = 0.1306 , *p* = 0.723 , partial $\eta^2$ = 0.009 |
| | Interaction OFF *p* = 0.653 | ON *p* = 0.4076 | Light (W) F(1,14) = 0.3935 , *p* = 0.541 , partial $\eta^2$ = 0.027 |
| | Exploration ON *p* = 0.839 | | Condition (B) F(1,14) = 1.4447 , *p* = 0.249 , partial $\eta^2$ = 0.094 |
| | Interaction ON *p* = 0.594 | | |
| | | Box's M test | post hoc (Bonferroni) |
| | **Sample Size** | *p* = 0.851 | Exploration ON x OFF *p* = 0.496 |
| | Exploration *n* = 8 | | Interaction ON x OFF *p* = 0.854 |
| | Interaction *n* = 8 | | |
| **Figure S13C** | **Shapiro-Wilk test** | **Levene's test** | **Two-Way mixed ANOVA** |
| | Total interaction time | | |
| | Exploration OFF *p* = 0.854 | OFF *p* = 0.529 | Interaction F(1,14) = 0.004 , *p* = 0.952 , partial $\eta^2$ = <0.001 |
| | Interaction OFF *p* = 0.442 | ON *p* = 0.866 | Light (W) F(1,14) = 0.010 , *p* = 0.921 , partial $\eta^2$ = <0.001 |
| | Exploration ON *p* = 0.407 | | Condition (B) F(1,14) = 0.037 , *p* = 0.851 , partial $\eta^2$ = 0.003 |
| | Interaction ON *p* = 0.648 | | |
| | | Box's M test | post hoc (Bonferroni) |
| | | *p* = 0.8334 | Exploration ON x OFF *p* = 0.978 |
| | | | Interaction ON x OFF *p* = 0.910 |
| | Number of visits | | **Two-Way mixed ANOVA** |
| | Exploration OFF *p* = 0.141 | OFF *p* = 0.9248 | Interaction F(1,14) = 0.7609 , *p* = 0.398 , partial $\eta^2$ = 0.052 |
| | Interaction OFF *p* = 0.453 | ON *p* = 0.5774 | Light (W) F(1,14) = 0.0016 , *p* = 0.968 , partial $\eta^2$ = <0.001 |
| | Exploration ON *p* = 0.191 | | Condition (B) F(1,14) = 0.1818 , *p* = 0.676 , partial $\eta^2$ = 0.013 |
| | Interaction ON *p* = 0.910 | | |
| | | Box's M test | post hoc (Bonferroni) |
| | | *p* = 0.6557 | Exploration ON x OFF *p* = 0.566 |
| | | | Interaction ON x OFF *p* = 0.529 |
| | Interaction time per visit | | **Two-Way mixed ANOVA** |
| | Exploration OFF *p* = 0.240 | OFF *p* = 0.4273 | Interaction F(1,14) = 1.4946 , *p* = 0.242 , partial $\eta^2$ = 0.096 |
| | Interaction OFF *p* = 0.509 | ON *p* = 0.1974 | Light (W) F(1,14) = 0.0029 , *p* = 0.958 , partial $\eta^2$ = <0.001 |
| | Exploration ON *p* = 0.578 | | Condition (B) F(1,14) = 0.2148 , *p* = 0.650 , partial $\eta^2$ = 0.015 |
| | Interaction ON *p* = 0.068 | | |
| | | Box's M test | post hoc (Bonferroni) |
| | | *p* = 0.8284 | Exploration ON x OFF *p* = 0.423 |
| | **Sample Size** | | Interaction ON x OFF *p* = 0.382 |
| | Exploration *n* = 8 | | |
| | Interaction *n* = 8 | | |
| **Figure S13D** | **Sample Size (# cells)** | **Expected Count <5** | **Chi-square test of homogeneity** |
| | Exploration eNPHR = 978 | 0% | Exploration eNPHR ON x OFF *p* = 0.836 |
| | Interaction eNPHR = 1076 | | Interaction eNPHR ON x OFF *p* = 0.479 |
| | Exploration Chrimson = 981 | | Exploration Chrimson ON x OFF *p* = 0.402 |
| | Interaction Chrimson = 1007 | | Interaction Chrimson ON x OFF *p* = 0.884 |
| | **Sample Size (# animals)** | | |
| | Exploration eNPHR *n* = 8 | | |
| | Interaction eNPHR *n* = 8 | | |
| | Exploration Chrimson *n* = 8 | | |
| | Interaction Chrimson *n* = 8 | | |
| **Figure S13E** | **Sample Size (# cells)** | **Expected Count <5** | **Chi-square test of homogeneity** |
| | Exploration eNPHR = 978 | 0% | Exploration eNPHR ON x OFF *p* = 0.525 |
| | Interaction eNPHR = 1076 | | Interaction eNPHR ON x OFF *p* = 0.829 |
| | Exploration Chrimson = 981 | | Exploration Chrimson ON x OFF *p* = 0.507 |
| | Interaction Chrimson = 1007 | | Interaction Chrimson ON x OFF *p* = 0.976 |
| | **Sample Size (# animals)** | | |
| | Exploration eNPHR *n* = 8 | | |
| | Interaction eNPHR *n* = 8 | | |
| | Exploration Chrimson *n* = 8 | | |
| | Interaction Chrimson *n* = 8 | | |

| | | | |
|---|---|---|---|
| **Figure S13F** | **Sample Size (# cells)**<br>Exploration eNPHR = 1021<br>Interaction eNPHR = 1111<br>Exploration Chrimson = 997<br>Interaction Chrimson = 998<br>**Sample Size (# animals)**<br>Exploration eNPHR *n* = 8<br>Interaction eNPHR *n* = 8<br>Exploration Chrimson *n* = 8<br>Interaction Chrimson *n* = 8 | **Expected Count <5**<br>0% | **Chi-square test of homogeneity**<br>Exploration eNPHR ON x OFF *p* = 0.339<br>Interaction eNPHR ON x OFF *p* = 0.127<br>Exploration Chrimson ON x OFF *p* = 0.752<br>Interaction Chrimson ON x OFF *p* = 0.162 |
| **Figure S13G** | **Sample Size (# cells)**<br>Interaction eNPHR = 1111<br>Interaction Chrimson = 998<br>**Sample Size (# animals)**<br>Interaction eNPHR *n* = 8<br>Interaction Chrimson *n* = 8 | **Expected Count <5**<br>0% | **Chi-square test of homogeneity**<br>eNPHR x Chrimson *p* = 0.485 |
| **Figure S14B** | **Shapiro-Wilk test**<br>Unstressed AI *p* = 0.337<br>Stressed AI *p* = 0.219<br>Unstressed PrL *p* = 0.878<br>Stressed PrL *p* = 0.839<br>**Sample Size**<br>Unstressed *n* = 6<br>Stressed *n* = 6 | **Levene's test**<br>Empty *p* = 0.8232<br>Mouse *p* = 0.2037<br>Box's M test<br>*p* = 0.355 | **Two-Way mixed ANOVA**<br>Interaction F(1,14) = 0.0038 , *p* = 0.952 , partial $\eta^2$ = <0.001<br>Region (W) F(1,14) = 27.355 , *p* = <0.001 , partial $\eta^2$ = 0.732<br>Stress (B) F(1,14) = 0.0445 , *p* = 0.837 , partial $\eta^2$ = 0.004<br>post hoc (Bonferroni)<br>AI Unstressed x Stressed *p* = 0.916<br>PrL Unstressed x Stressed *p* = 0.811 |
| **Figure S15C** | **Sample Size (# cells)**<br>*n* = 503<br>**Sample Size (# animals)**<br>Female B6J mice *n* = 4 | **NA** | **NA** |
| **Figure S15F** | **Sample Size (# cells)**<br>*n* = 2007<br>**Sample Size (# animals)**<br>Female B6J mice *n* = 4 | **NA** | **NA** |
| **Figure S16C** | **Sample Size (# cells)**<br>*n* = 658<br>**Sample Size (# animals)**<br>Female B6J mice *n* = 4 | **NA** | **NA** |
| **Figure S16D** | **Shapiro-Wilk test**<br>Total interaction time<br>Unstressed Control p = 0.838<br>Unstressed GR-KO p = 0.638<br>Stressed Control p = 0.808<br>Stressed GR-KO p = 0.638<br><br>Number of visits<br>Unstressed Control p = 0.022<br>Unstressed GR-KO p = 0.939<br>Stressed Control p = 0.087<br>Stressed GR-KO p = 0.491<br><br>Interaction time per visit<br>Unstressed Control p = 0.029<br>Unstressed GR-KO p = 0.557<br>Stressed Control p = 0.418<br>Stressed GR-KO p = 0.106<br>**Sample Size**<br>Unstressed Control *n* = 6<br>Unstressed GR-KO *n* = 6<br>Stressed Control *n* = 6<br>Stressed GR-KO *n* = 6 | **Levene's test**<br>*p* = 0.713<br><br>*p* = 0.6773<br><br>*p* = 0.4075 | **Two-Way ANOVA**<br>Interaction F(3,20) = 0.792 , *p* = 0.384 , partial $\eta^2$ = 0.038<br>Stress (B) F(3,20) = 0.009 , *p* = 0.927 , partial $\eta^2$ = <0.001<br>KO (B) F(3,20) = 0.741 , *p* = 0.400 , partial $\eta^2$ = 0.036<br>post hoc (Bonferroni)<br>Unstressed Control x GR-KO *p* = 0.230<br>Stressed Control x GR-KO *p* = 0.984<br><br>**Mann Whitney U test**<br>Unstressed Control x GR-KO *p* = 0.699 *U* = 15.5<br>*r* = -0.1 *z* = -0.402<br>Stressed Control x GR-KO *p* = 1.000 *U* = 18.5<br>*r* = 0.02 *z* = 0.08<br><br>**Mann Whitney U test**<br>Unstressed Control x GR-KO *p* = 0.065 *U* = 6<br>*r* = -0.48 *z* = -1.922<br>Stressed Control x GR-KO *p* = 0.699 *U* = 21<br>*r* = 0.12 *z* = 0.48 |
